# bioPROTACs establish RAS as a degradable target and provide novel RAS biology insights

**DOI:** 10.1101/2020.06.26.174565

**Authors:** Shuhui Lim, Regina Khoo, Yu-Chi Juang, Pooja Gopal, Huibin Zhang, Constance Yeo, Khong Ming Peh, Jinkai Teo, Simon Ng, Brian Henry, Anthony W. Partridge

## Abstract

Mutations to RAS proteins (H-, N-, and K-RAS) are amongst the most common oncogenic drivers and tumors harboring these lesions are some of the most difficult to treat. Although the recently discovered covalent small molecules against the KRAS^G12C^ mutant have shown promising efficacy against lung cancers, traditional barriers remain for drugging the more prevalent KRAS^G12D^ and KRAS^G12V^ mutants. Targeted degradation has emerged as an attractive alternative approach but for KRAS, identification of the required high-affinity ligands continues to be a challenge. Another significant hurdle is the discovery of a hybrid molecule that appends an E3 ligase-recruiting moiety in a manner that satisfies the precise geometries required for productive polyubiquitin transfer while maintaining favorable drug-like properties. As a tool to gain insights into the advantages and feasibility of KRAS targeted-degradation, we applied the bioPROTAC approach. This workflow centers on the intracellular expression of a chimeric protein consisting of a high-affinity target-binding domain fused to an engineered E3 ligase adapter. We generated a series of anti-RAS bioPROTACs that span different RAS isoform/nucleotide-state specificities and leverage different E3 ligases. Overall, our results provide definitive evidence for the degradability of RAS proteins. We further elucidate the functional consequences of RAS degradation, the susceptibility and degradation kinetics of various mutant KRAS, and the prevalence of different nucleotide-states in WT and mutant KRAS. Finally, if delivery challenges can be addressed, anti-RAS bioPROTACs will be exciting candidates for clinical development.

## INTRODUCTION

Mutations to RAS proteins are amongst the most frequent drivers of human cancers with approximately 30% of all clinical malignancies containing an activating RAS mutation^1^. KRAS is the most frequently mutated RAS isoform (86%), followed by NRAS (11%) and HRAS (3%)^2^. With a primary focus on KRAS, researchers have therapeutically pursued RAS oncogenes for nearly 40 years. Unfortunately, the intractability of this target to conventional approaches has impeded the identification of a clinically approved drug. However, recent advances are giving renewed hope that pharmacological inhibition of KRAS can finally be realized. In particular, recently discovered covalent inhibitors targeting the KRAS^G12C^ mutant protein are showing promising clinical efficacy^3,4^, further validating mutant KRAS as a clinically relevant oncology target. In preclinical mouse models, these inhibitors have shown robust blockade of KRAS signaling and cell proliferation^3,4^. Combinations with immunotherapy has led to increased efficacy and immune memory^3^. More importantly, early Phase I clinical data with G12C inhibitor monotherapy has recorded responses in lung and, to a lesser degree, colon cancers^3,4^. Despite these significant advances, the covalent strategy is thus far restricted to the relatively rare G12C mutation (found in 14% of non-small cell lung cancers, 5% of colorectal cancers, and 2% of pancreatic cancers).

For non-G12C mutations, traditional challenges for identifying therapeutic molecules remain. In particular, identification of high affinity non-covalent ligands against active KRAS has proven refractory - a consequence of the lack of appropriate pockets for a small molecule to bind. Removal of the covalent warhead and reinforcement of binding energies through non-covalent interactions is an approach worth considering. However, this binding pocket is occluded in the GTP-loaded state^5^ and it remains unclear if non-G12C mutants cycle between nucleotide-states rapidly enough for this approach to be effective. Overall, alternative strategies need to be considered. Amongst these, small molecule targeted-degradation approaches, such as proteolysis targeting chimeras (PROTACs), have recently generated a lot of excitement^6-10^. These bifunctional molecules consist of a target-binding moiety linked to an E3-recruiting ligand. Successfully engineered PROTAC molecules not only recruit the corresponding E3/E2 complex to the vicinity of the target-of-interest, but also form productive ternary complexes that induce the transfer of polyubiquitin to the target to result in its proteasomal degradation^7^. This strategy opens up new possibilities to tackle historically intractable targets since degradation is potentially achievable *via* engagement with a variety of binding sites - including but not restricted to those of functional consequence^8,11^. Moreover, recent examples illustrate that targeted degradation offers better efficacy, potency, and selectivity^8,12^. Finally, given the high intracellular concentration of KRAS^13-15^ (also **Supplementary Fig. 1**), achieving adequate target engagement with non-covalent stoichiometric inhibitors may be challenging.

As there are substantial challenges in identifying small molecule PROTACs, initial investigations aimed at assessing PROTAC feasibility and providing insights on optimal design strategies are warranted. Key considerations include I) target degradability through engineered polyubiquitin transfer, II) ‘fitness’ of the E3 ligases recruited, III) interfaces on the target protein that can be bound yet remain amenable to polyubiquitination, and IV) the functional consequences of target degradation. To resolve these questions, we have employed engineered fusion proteins termed bioPROTACs^16^, also known as ubiquibodies^17^, AdPROMs^18^, and deGradFP^19^. bioPROTACs consist of a target-binding domain connected to an E3 ligase (E3). A variety of polypeptide scaffolds evolved to recognize the target with high affinity and specificity can be selected as the target-binding domain^16^. Indeed, active bioPROTACs have been generated with fusions between E3s and nanobodies, monobodies, alpha-reps, DARPins, and peptides^16,17^. The choice of E3 is also flexible, with functional bioPROTACs having been engineered from both human and bacterial sequences^16,20^.

Although a recent attempt at engineering a small molecule PROTAC against KRAS^G12C^ using a covalent modifier^21^ failed to induce polyubiquitin-mediated degradation, other data suggest that RAS is indeed degradable. First, the natural turn-over of RAS proteins was reported to be proteasome-dependent and regulated by the E3 ligases LTZR1^22-24^ and βTrCP^25^. Second, the G12C covalent modifier and bioPROTAC approaches have been successful for degrading GFP-KRAS^20,21^. Third, bioPROTAC equivalents consisting of the endogenous RAS-binding-domain (RBD) fused to either VIF or CHIP E3 ligases have resulted in modest KRAS degradation^26,27^. Here, we report the discovery of a panel of novel and potent KRAS-directed bioPROTACs that build on these earlier results and provide conclusive evidence for the degradability of various RAS isoforms and mutant proteins. By utilizing a variety of E3 ligases, our study unveils the possibility of engaging novel E3 ligases for a KRAS PROTAC campaign beyond VHL and Cereblon. By exploring a variety of RAS binding moieties, we shed light on KRAS interfaces that can be exploited for the design of small molecule PROTACs. We further demonstrate that both GTP- and GDP-loaded forms of RAS proteins are amenable to targeted degradation. A bioPROTAC specific for GDP-loaded RAS (K27-SPOP) degraded wild-type and KRAS mutants (G12C, G12D, G12V and Q61H) with different efficiencies; an observation that informs on the capacity of these mutants to cycle through nucleotide-states in the cellular environment. We also show that mRNA-mediated delivery of anti-RAS bioPROTACs degraded endogenous mutant KRAS, resulting in growth inhibition and apoptosis in a KRAS-dependent cancer cell line and provide an example where targeted degradation is superior in comparison to stoichiometric inhibition.

## RESULTS

### GFP-KRAS is degraded by multiple anti-GFP bioPROTACs

As a starting point to determine if KRAS proteins can be targeted for ubiquitin-mediated proteasomal degradation, we applied our anti-GFP bioPROTAC platform^16^, which features a panel of 10 representative Cullin-RING E3 ubiquitin ligase (CRL) family members fused to the GFP-binding nanobody vhhGFP4^28,29^ (**Fig. 1a**). By tagging KRAS with GFP, we sought to recruit an assortment of ubiquitination complexes to the vicinity of KRAS and evaluate its degradability. HEK293 stable cell lines with constitutive expression of GFP or GFP-KRAS were established and the panel of anti-GFP bioPROTACs were individually transfected with mCherry as an expression reporter. Flow cytometry was used to determine GFP levels in mCherry-positive (transfected) cells (**Fig. 1b**). As noted previously^16^, GFP alone was poorly degraded by our panel of anti-GFP PROTACs (**Fig. 1c left column**). However, when fused to KRAS, GFP signal intensities were attenuated by 8 out of 10 bioPROTACs, with 6 of them (βTrCP, FBW7, SKP2, SPOP, SOCS2 and CHIP) having more than 70% of transfected cells in the GFP-negative quadrant (Q1) (**Fig. 1c right column**) 24 hours following transfection. Similar to observations against other targets^16^, both CUL4-based (CRBN and DDB2) bioPROTACs failed to degrade GFP-KRAS; we speculate this is likely due to issues related to protein engineering rather than the incompatibility of these E3 ligases. The depletion of GFP-KRAS, but not GFP, suggests that KRAS itself likely possesses the necessary traits for proteasomal degradation (i.e. solvent-exposed lysines for poly-ubiquitination and a structurally disordered segment that initiates unfolding at the 26S proteasome^30^).

**Figure 1.**
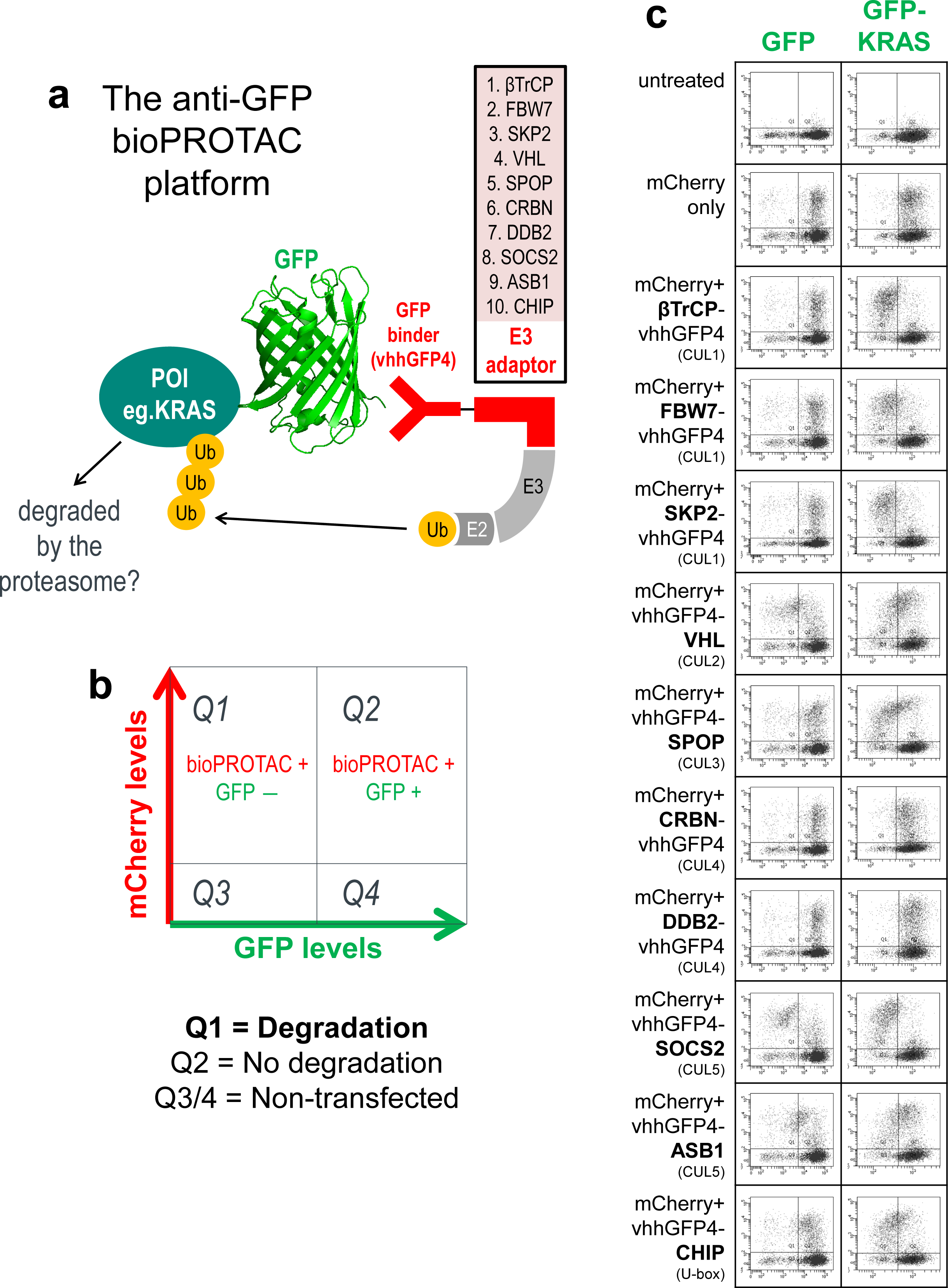
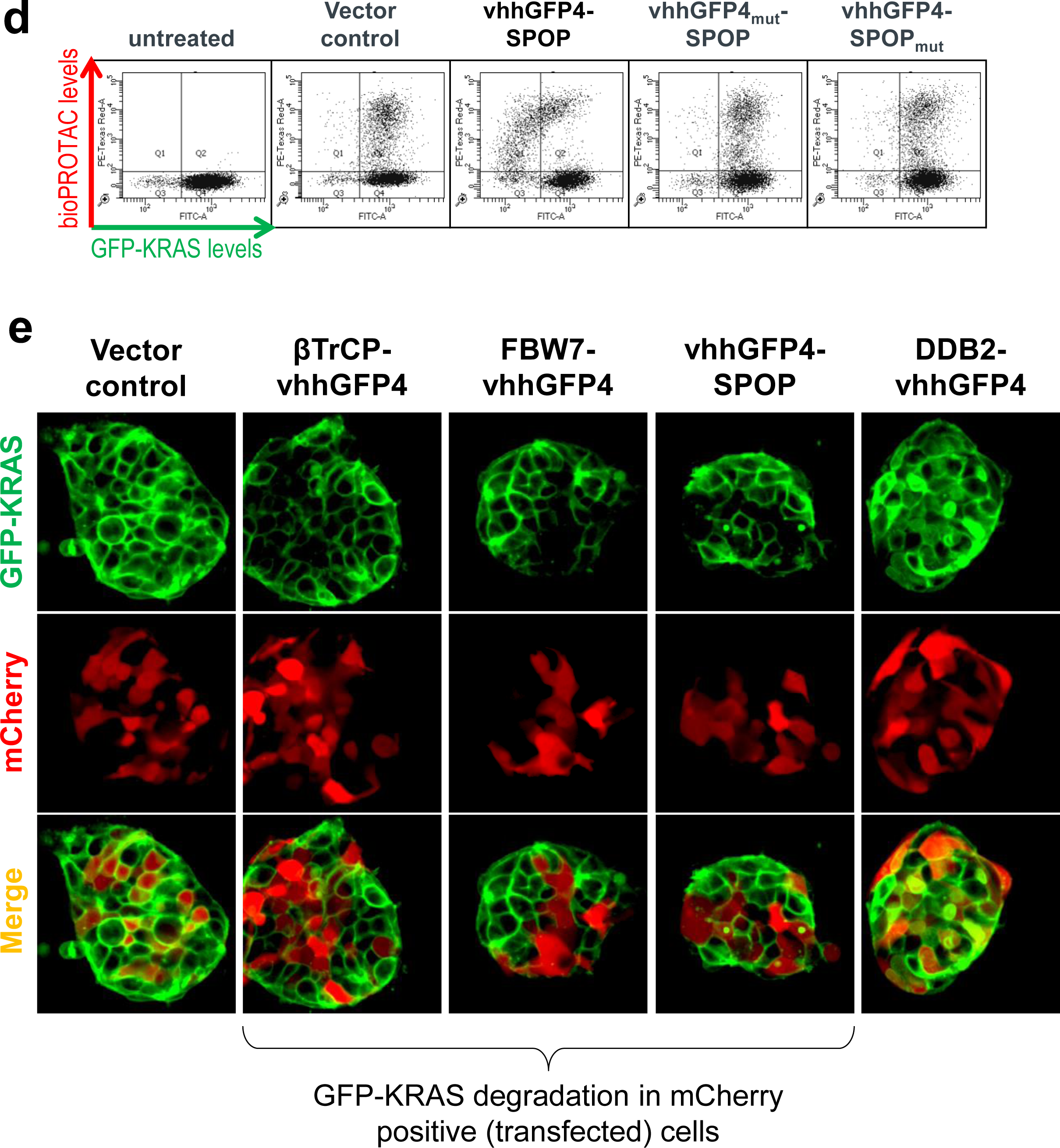
GFP-KRAS is degraded by multiple anti-GFP bioPROTACs. (**a**) Schematic of the anti-GFP bioPROTAC platform used to evaluate the degradability of a protein-of-interest (POI) fused to GFP. GFP is bound by vhhGFP4, a high-affinity anti-GFP nanobody, thereby bringing an E3 adaptor in close proximity to the POI. The collection of ten E3 adaptors span representative members of the Cullin-RING E3 ligase (CRL) family. (**b**) Flow cytometry is used to determine the levels of the GFP-tagged protein. Transfected cells that express anti-GFP bioPROTAC will be mCherry-positive and therefore reside in quadrants 1 and 2 (Q1 and Q2). Successful degradation will reduce GFP signal and cells will cumulate in Q1. Cells with no degradation will be retained in Q2. (**c**) Flow cytometric analysis of HEK293 Tet-On® 3G cells with stable integration of GFP or GFP-KRAS and transiently transfected with the panel of ten anti-GFP bioPROTACs. (**d**) Flow cytometric analysis of HEK293 Tet-On® 3G cells with stable integration of GFP-KRAS and transiently transfected with vhhGFP4-SPOP or its controls. vhhGFP4_mut_ lacks the complementarity determining region 3 (CDR3) and no longer recognizes GFP, whereas SPOP_mut_ lacks the 3-box motif responsible for recruiting CUL3 and thus cannot assemble the ubiquitination machinery. (**e**) Confocal imaging analysis of HEK293 Tet-On® 3G cells with stable integration of GFP-KRAS (green) and transiently transfected with the indicated anti-GFP bioPROTACs. mCherry (red) is a reporter of transfected cells.

For some of the active bioPROTACs such as vhhGFP4-SPOP, a characteristic hook-effect was observed (**Fig. 1c and 1d**). This is caused by excessively high PROTAC concentrations which compromises degradation by decreasing the probability of ternary complex formation in favor of substrate:PROTAC and PROTAC:E3 binary complexes^31^. Mutations to the binding domain (vhhGFP4_mut_) or the E3 ligase (SPOP_mut_) completely abrogated the downregulation of GFP-KRAS (**Fig. 1d**), suggesting that both components of the chimeric protein are essential for bioPROTAC activity. The targeted degradation of GFP-KRAS by anti-GFP bioPROTACs was further corroborated with confocal imaging. Like endogenous KRAS^32^, the subcellular localization of GFP-KRAS was predominantly membrane-bound (**Fig. 1e**). Transient expression of mCherry alone did not affect the levels and localization of GFP-KRAS (**Fig. 1e first column**). However, when co-expressed with βTrCP-vhhGFP4, FBW7-vhhGFP4 or vhhGFP4-SPOP, the membrane-localized green fluorescence was specifically lost in mCherry positive (transfected) cells (**Fig. 1e middle 3 columns**). DDB2-vhhGFP4 was identified as a non-degrader from the flow cytometric screen (**Fig. 1c**). Interestingly, upon the expression of DDB2-vhhGFP4, GFP-KRAS was redistributed to the cytoplasm/nucleus (**Fig. 1e last column**), suggesting that this bioPROTAC can bind GFP-KRAS but lacks the ability to induce its degradation. This observation also shows that a nuclear-localized E3 is still able to access a membrane-bound/cytoplasmic substrate. Overall, the anti-GFP bioPROTAC platform established GFP-KRAS as an amenable substrate and identified suitable E3s that can be employed to elicit proteasomal degradation.

### Leveraging high affinity RAS binders for endogenous RAS degradation

Having successfully demonstrated the degradability of GFP-KRAS, we were prompted to design anti-RAS bioPROTACs that can be used to directly degrade endogenous KRAS. This involves the fusion of a KRAS binder to an appropriate E3 ligase. Based on published sources, we shortlisted five KRAS binders that interact at different interfaces (**Fig. 2a**) and further validated their reported affinities and isoform/nucleotide specificities using Isothermal Titration Calorimetry (ITC). NS1 is a monobody that binds KRAS and HRAS, but not NRAS^33^ (**Supplementary Fig. 2a**). The DARPins, K27 and K55, are specific for GDP- and GTP-loaded KRAS respectively^34^ (**Supplementary Fig. 2b and 2c**). R11.1.6 is based on the ultra-stable Sso7d scaffold and was described to be mutant KRAS-selective^35^. Unfortunately, we were unable to purify sufficient quantities of recombinant R11.1.6 for biophysical analysis. We also tested the RAS-binding domain (RBD)^36^, a conserved region in RAS effector proteins (e.g. RAF, PI3K and TIAM1) that interacts specifically with activated GTP-bound RAS. The RBD of RAF1 was made and its affinity for GMPPCP-loaded KRAS^G12D^ was measured at 59 nM (**Supplementary Fig. 2d**).

**Figure 2.**
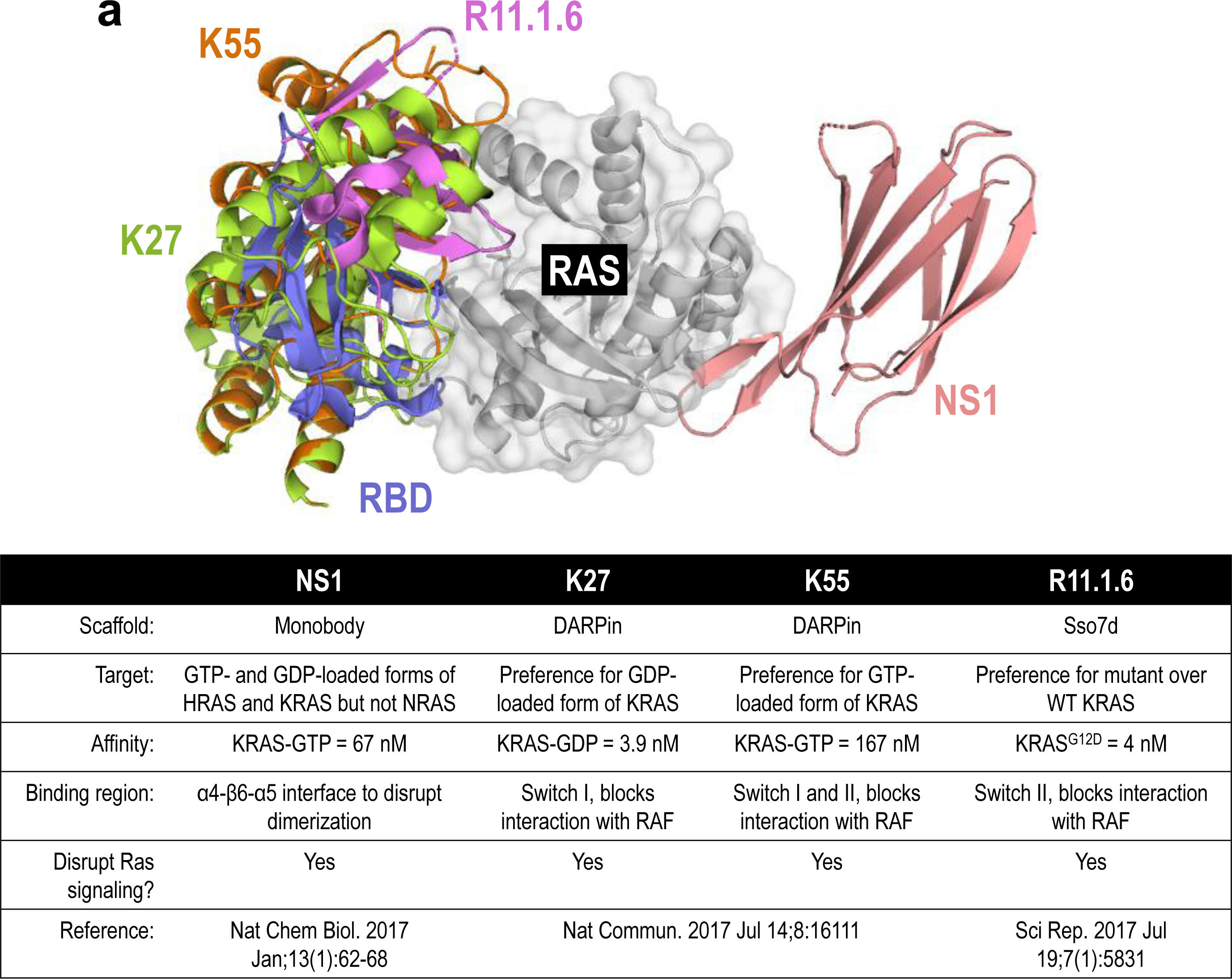
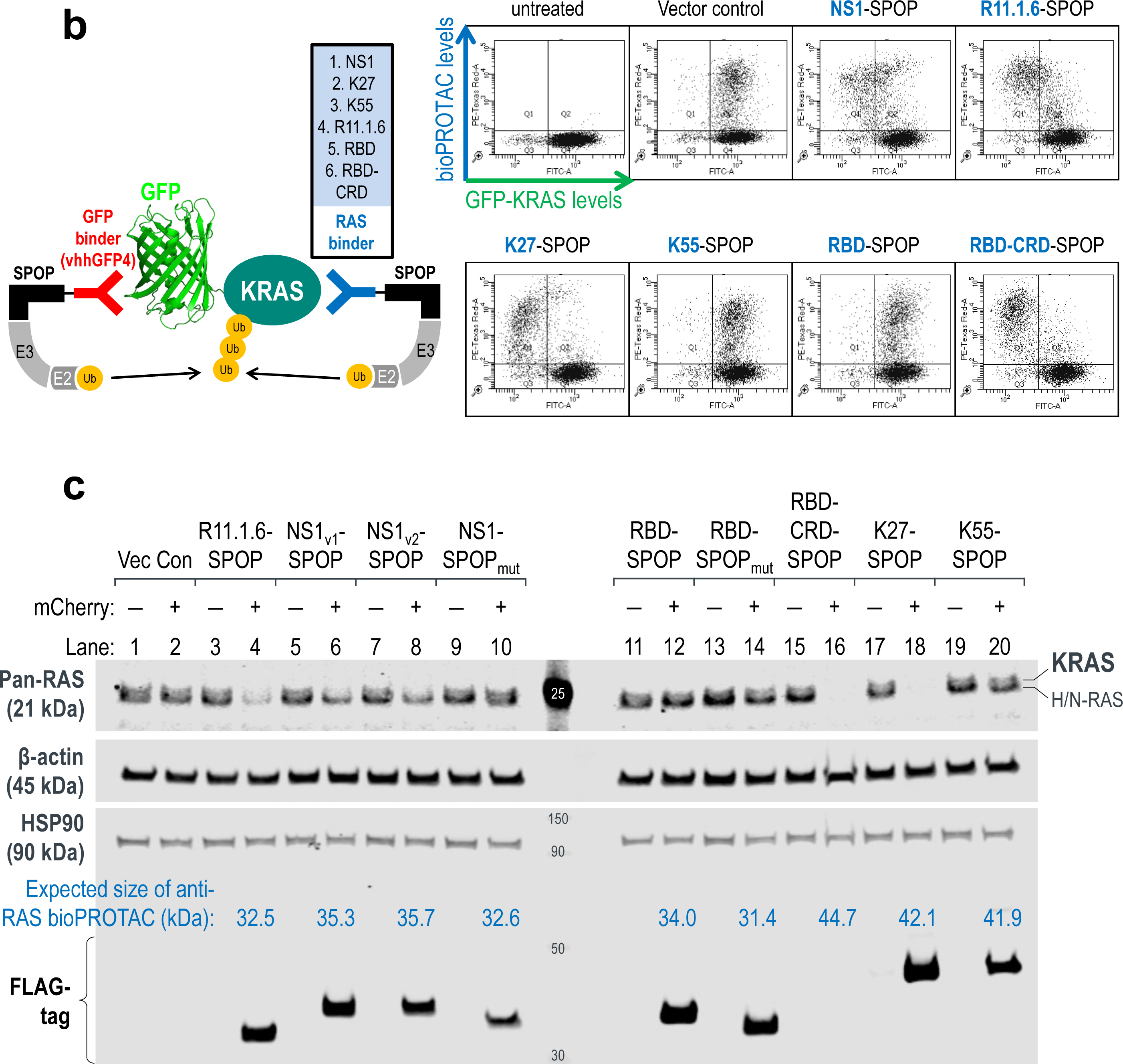
Leveraging high affinity binders for endogenous RAS degradation. (**a**) Overlay of KRAS binders from literature sources and a table summarizing their reported binding specificities and affinities. PDB structures used were: 5E95 (NS1), 5O2S (K27), 5O2T (K55), 5UFQ (R11.1.6) and 4G0N (RBD). (**b**) Flow cytometric analysis of HEK293 Tet-On® 3G cells with stable integration of GFP-KRAS and transiently transfected with anti-RAS bioPROTACs (in blue). Cells in Q1 represent successful GFP-KRAS depletion by the respective bioPROTAC. (**c**) Western blot analysis of HEK293 Tet-On® 3G cells transiently transfected with the indicated anti-RAS bioPROTACs and sorted according to the levels of mCherry (a marker of transfected cells) using FACS. Gating was set such that mCherry (-) cells have the same signal intensities as untreated cells in the mCherry channel, and anything above this basal level was assigned mCherry (+). In the pan-RAS blot, the upper band corresponds to KRAS while the lower band corresponds to HRAS and NRAS. Expression of the various anti-RAS bioPROTACs was detected using an anti-FLAG-tag antibody and the expected molecular weight of each chimeric protein is indicated in kilodaltons (kDa). β-actin and HSP90 were used as loading controls.

Our previous work^16^ and the screen described above (**Fig. 1c – 1e**) identified SPOP as a highly robust E3 ligase. Thus, we coupled each of the RAS binders to SPOP to generate anti-RAS bioPROTACs. To rapidly screen for PROTAC activity, GFP-KRAS was picked as the initial substrate. Through their abilities to directly engage KRAS, NS1-SPOP, K27-SPOP, R11.1.6-SPOP were all able to deplete the GFP signal (**Fig. 2b**). Interestingly, whilst RBD-SPOP did not degrade GFP-KRAS, the addition of the cysteine-rich domain (CRD) that was reported to anchor RAF proteins on membrane patches and stabilize RAS-RAF interactions^37,38^ yielded an active bioPROTAC (RBD-CRD-SPOP) (**Fig. 2b**). This exemplifies how increased avidity through membrane targeting could aid in the stabilization of ternary complex formation required for productive degradation of GTP-loaded KRAS. As KRAS switches to the ‘ON’ state when bound to GTP, it engages in protein-protein interactions with a multitude of effector proteins, many of which are membrane localized. Hence, bioPROTACs that target GTP-loaded KRAS might benefit from increased membrane localization. This could explain why K55-SPOP was ineffective (**Fig. 2b**) since it lacks membrane targeting. It is also worth noting that the affinity of K55 for GTP-loaded KRAS is 98 nM (**Supplementary Fig. 2c**), weaker than the endogenous RAS binder RBD, which is 59 nM (**Supplementary Fig. 2d**).

To probe for the degradation of endogenous RAS, we next transfected HEK293 cells with doxycycline-inducible DNA plasmids driving co-expression of anti-RAS bioPROTACs and mCherry reporter. Twenty-four hours post-induction, cells were sorted into mCherry-negative (non-transfected) and mCherry-positive (transfected) populations and harvested for Western blot analysis. A pan-RAS antibody was used to probe for endogenous levels of RAS family proteins: KRAS, HRAS and NRAS, which appeared as two bands in HEK293. A previous study using isoform-specific siRNAs demonstrated that the upper band corresponds to KRAS, whereas the lower band corresponds to HRAS and NRAS^39^. In our experiments, the upper KRAS band was specifically lost with the expression of NS1-SPOP (**Fig. 2c lanes 6 and 8**) but not with the non-degrading control NS1-SPOP_mut_ (**Fig. 2c lane 10**). These data suggest that it is possible to achieve selective degradation of closely-related proteins if isoform-specificities are engineered into the binders. To understand if the degradation of RAS is affected by its guanine nucleotide status, we used K27 (pan-RAS, specific for the GDP-loaded state) and RBD-CRD (pan-RAS, specific for the GTP-loaded state) as the substrate binding moieties. The expression of either K27-SPOP or RBD-CRD-SPOP led to complete disappearance of pan-RAS bands (**Fig. 2c lanes 16 and 18**), suggesting that both nucleotide-states across RAS isoforms are susceptible to degradation. Consistent with the results on GFP-KRAS (**Fig. 2b**), K55-SPOP and RBD-SPOP failed to degrade endogenous RAS (**Fig. 2c lanes 12 and 20**). R11.1.6-SPOP partially reduced pan-RAS band intensities (**Fig. 2c lane 4**). The preferential binding of R11.1.6 to mutant KRAS^35^ could explain why there was incomplete degradation in HEK293 cells where the status of RAS is wild-type. All anti-RAS bioPROTACs were FLAG-tagged and expressed according to the expected sizes and at similar levels, with the exception of RBD-CRD-SPOP (**Fig. 2c lane 16**). This bioPROTAC was also barely detectable in repeat experiments (**Supplementary Fig. 3 lane 10)**. Using cell sorting, we were able to include the mCherry-negative (non-transfected) population as an internal control for RAS levels in all cases (**Fig. 2c lanes marked as mCherry –**).

It is often challenging to achieve 100% efficiency with DNA transfection. In order to better characterize anti-RAS bioPROTACs and study the functional consequences of KRAS loss, we generated HEK293 stable cell lines with doxycycline-inducible expression of the various anti-RAS bioPROTACs. Pan-RAS deletion was achieved as early as 4 hours post-induction of K27-SPOP. This effect persisted for up to 24 hours (**Fig. 3a lanes 2 – 5, first panel**) and coincided with inhibition of phospho-ERK1/2, a downstream effector of the mitogen-activated protein kinase (MAPK) pathway (**Fig. 3a lanes 2 – 5, second panel**). With SPOP mutated, the E3 ligase activity of K27-SPOP_mut_ is disabled and such that pan-RAS protein levels were not affected (**Fig. 3a lanes 7 – 10, first panel**). However, K27 on its own was reported to have inhibitory effects on the MAPK pathway^34^ and indeed, phospho-ERK1/2 levels were reduced 4 hours after the induction of K27-SPOP_mut_ (**Fig. 3a lanes 7 – 8, second panel**). However, this inhibitory effect could not be sustained and phospho-ERK1/2 levels returned to baseline at 24 hours (**Fig. 3a lanes 9 – 10, second panel**), despite continued K27-SPOP_mut_ expression (**Fig. 3a lanes 9 – 10, third panel**). The non-binder control K27_mut_-SPOP, wherein three RAS-binding residues were replaced by alanine^34^, did not alter pan-RAS nor phospho-ERK1/2 levels as expected (**Fig. 3a lanes 11 – 15**). Stable cell lines with doxycycline-inducible expression of other anti-RAS bioPROTACs, such as R11.1.6-SPOP, NS1-SPOP and K27-VHL, were also generated (**Supplementary Fig. 4**) but K27-SPOP demonstrated the most complete RAS degradation and sustained phospho-ERK inhibition in HEK293 cells. Surprisingly, despite strong RAS knockdown, HEK293 cells expressing K27-SPOP continued to proliferate at rates similar to controls (**Fig. 3b**). Western blotting for pan-RAS confirmed that the cells proliferated in the absence of RAS proteins (**Fig. 3c**). These data suggest that HEK293 cells are not dependent on RAS proteins for survival.

**Figure 3.**
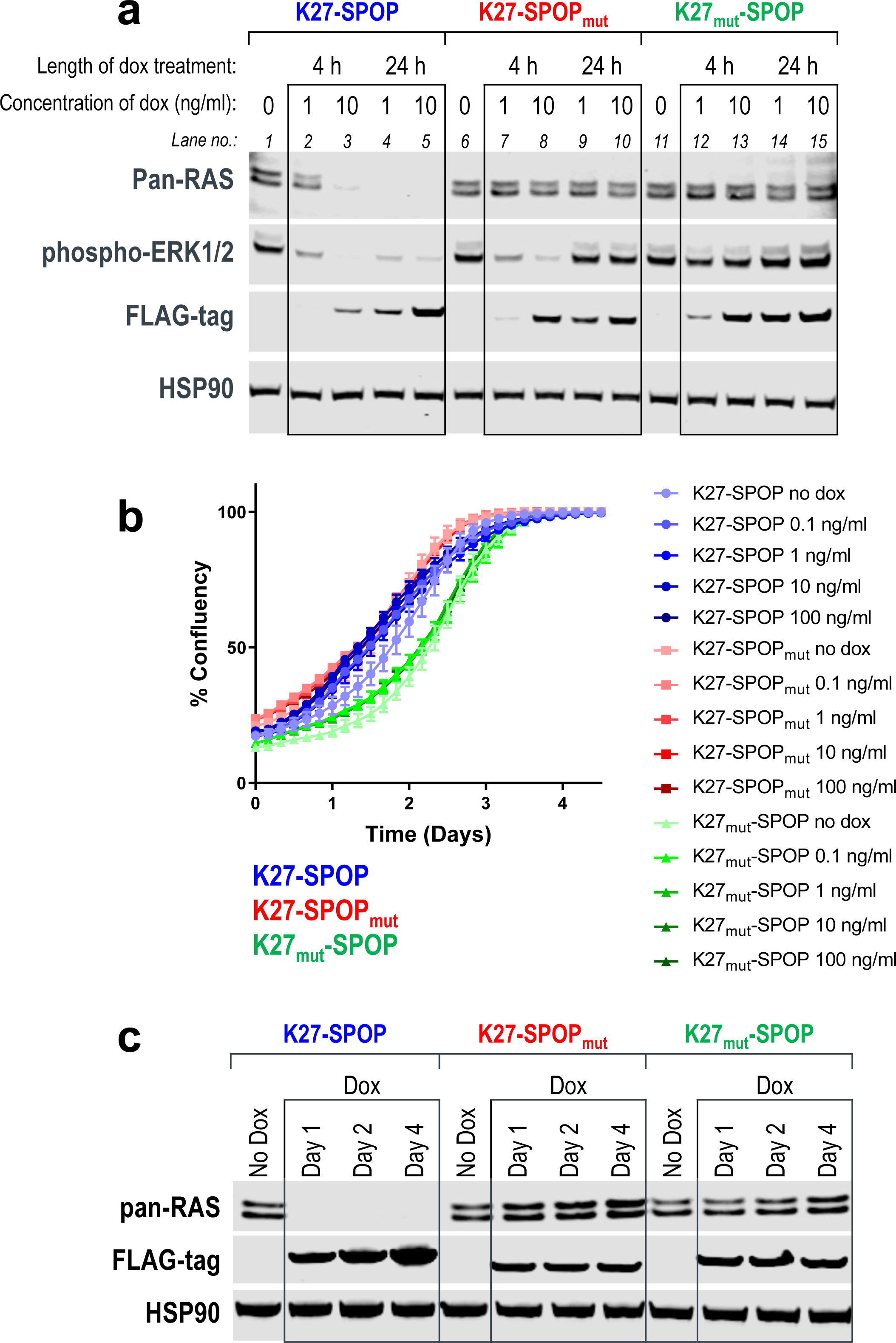
Robust RAS degradation with doxycycline-inducible anti-RAS bioPROTACs. (**a**) Western blot analysis of T-REx™-293 cells with stable integration of K27-SPOP (or its controls) under the control of a Tet-responsive promoter. Various concentrations of doxycycline (1 or 10 ng/ml) were added to the culture media for the indicated length of time (4 or 24 hours) and protein lysates were collected. Degradation of RAS was detected using a pan-RAS antibody and disruption to the MAPK pathway was measured using the levels of phospho-ERK1/2. Expression of K27-SPOP (or its controls) was detected using an anti-FLAG-tag antibody. HSP90 was used as a loading control. (**b**) Incucyte confluency measurements of T-REx™-293 cells with stable integration of K27-SPOP (or its controls) under the control of a Tet-responsive promoter. Various concentrations of doxycycline (0.1 to 100 ng/ml) were added to the culture media and the percentage confluency of the cells was tracked continuously over 4 days. (**c**) Western blot analysis as in (**a**) on protein lysates collected at 1, 2, or 4 days after treatment with 1 ng/ml doxycycline.

### Mutant KRAS degradation, inhibition of proliferation and induction of apoptosis with mRNA-mediated expression of anti-RAS bioPROTACs

To extend our study of bioPROTAC-mediated KRAS degradation to mutant KRAS-dependent cancer cells, we employed mRNA transfection to yield higher transfection rates. As an example, in AsPC-1 cells (pancreatic adenocarcinoma cell line, homozygous KRAS^G12D^), transfection efficiencies of a GFP-encoding DNA plasmid versus GFP mRNA were 1% and 90% respectively after 12 hours (**Supplementary Fig. 5**). High mRNA transfection efficiency was also seen in a panel of 14 cancer cell lines, wherein 9 of the lines were more than 80% transfected at 24 hours (**Supplementary Fig. 6**). Leveraging this workflow, we transfected AsPC-1 cells with K27-SPOP mRNA and observed pan-RAS degradation and corresponding phospho-ERK1/2 inhibition within 4 hours (**Fig. 4a**). This effect persisted for up to 24 hours and ultimately resulted in growth inhibition of AsPC-1 cells at all three mRNA concentrations tested (**Fig. 4b**). These data suggest that the KRAS^G12D^ mutant protein retains adequate intrinsic hydrolysis to cycle back to the GDP-loaded state, where it can be effectively targeted by a GDP-specific bioPROTAC such as K27-SPOP. On the contrary, although the stoichiometric inhibitor K27-SPOP_mut_ was initially successful at disrupting ERK1/2 phosphorylation, the effects were not sustained (**Fig. 4a**) and cells expressing K27-SPOP_mut_ showed similar proliferation rates as the non-binding control K27_mut_-SPOP (**Fig. 4b**). Morphologically, AsPC-1 cells transfected with the K27-SPOP bioPROTAC appeared rounded up (**Fig. 4c**) and increased cleaved caspase-3 levels revealed that they were undergoing apoptosis (**Fig. 4d**). Overall, our data highlights the superiority of employing an event-driven strategy (such as PROTAC)^40^ for inhibiting KRAS rather than an occupancy-driven stoichiometric inhibitor approach.

**Figure 4.**
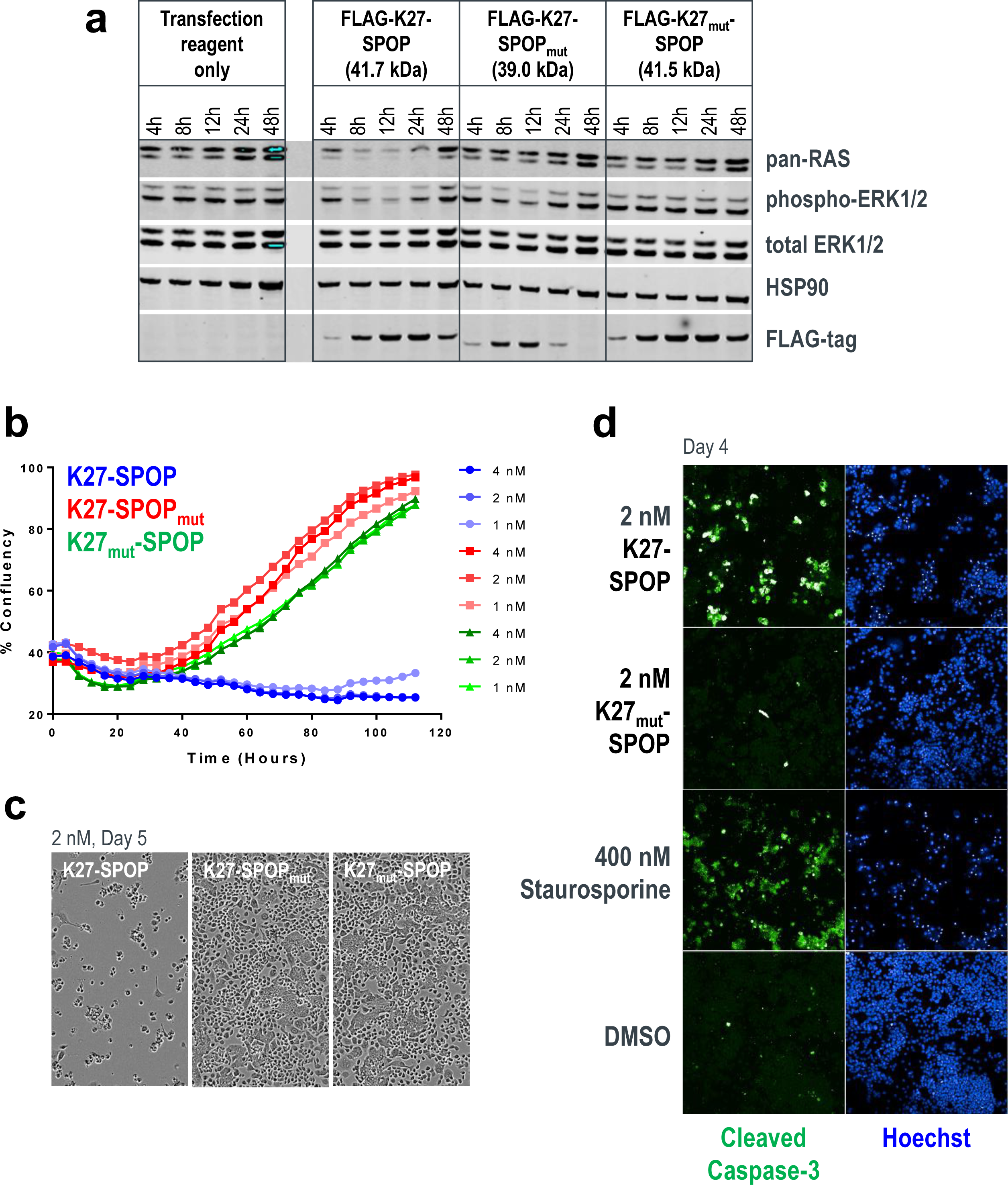
Mutant KRAS degradation, inhibition of proliferation and induction of apoptosis in AsPC-1 cells. (**a**) Western blot analysis of AsPC-1 cells transfected with mRNA encoding K27-SPOP (or its controls). Protein lysates were collected 4, 8, 12, 24, and 48 hours post-transfection. Degradation of RAS was detected using a pan-RAS antibody and disruption to the MAPK pathway was measured using the levels of phospho-ERK1/2. Expression of K27-SPOP (or its controls) was detected using an anti-FLAG-tag antibody. HSP90 was used as a loading control. (**b**) Incucyte confluency measurements of AsPC-1 cells transfected with mRNA as in (**a**) and tracked continuously over 5 days. (**c**) Phase-contrast images acquired 5 days post-transfection of AsPC-1 cells with 2 nM of K27-SPOP mRNA (or its controls). (**d**) Immunostaining for the levels of cleaved caspase-3, an indicator of apoptosis, 4 days post-transfection of AsPC-1 cells with 2 nM of K27-SPOP mRNA (or its control). Treatment with 400 nM staurosporine was used as a positive control for apoptotic cells.

### Establishment of the NanoLuc assay to inform on degradation selectivity and quantify degradation rates

We sought analytical methods to better characterize the isoform specificities and degradation efficacies of our anti-RAS bioPROTACs. Similar to the recently reported HiBiT-LgBiT platform^41^, we established a series of inducible NanoLuc-tagged RAS cell lines to track substrate levels real-time in live cells and report quantitative metrics of degradation efficiencies (**Fig. 5a**). Although the HiBiT platform has the advantage of using a smaller tag and reports on endogenous levels of the target protein, HiBiT knock-in cell lines are time-consuming to generate. Conversely, the NanoLuc approach can be established rapidly, enabling a comprehensive assessment of degradation kinetics for any RAS isoform or mutant protein in the same genetic background.

**Figure 5.**
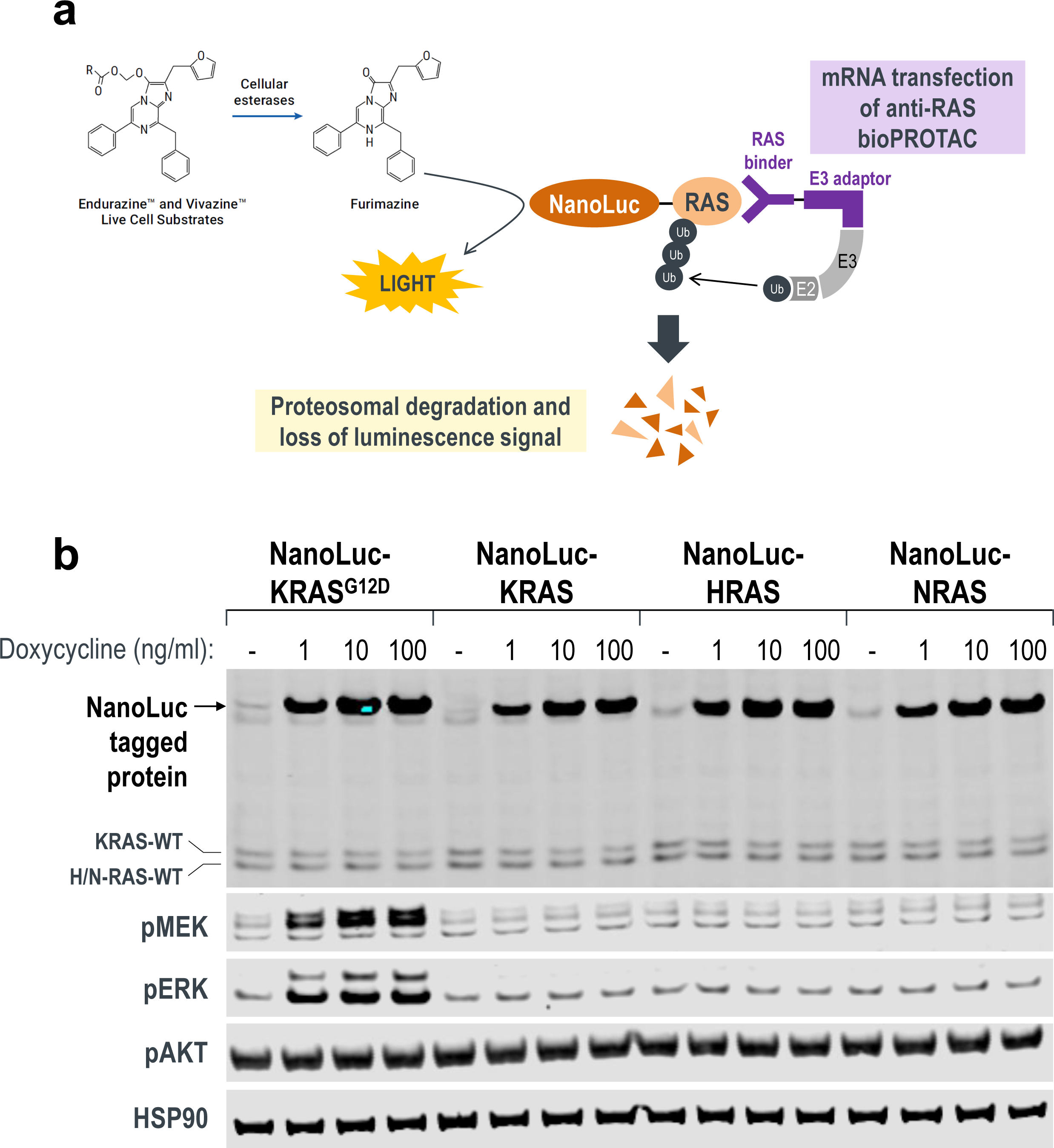
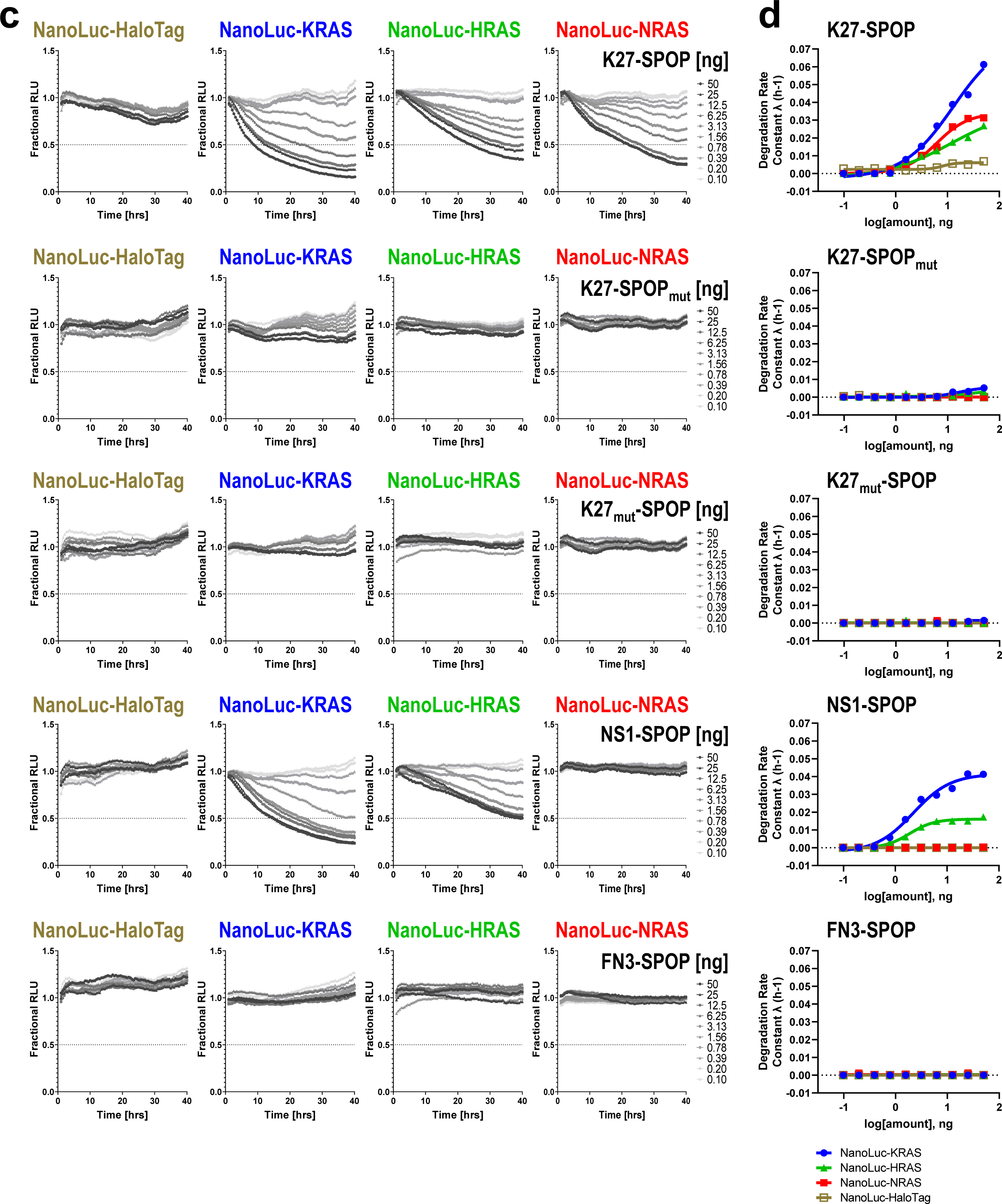
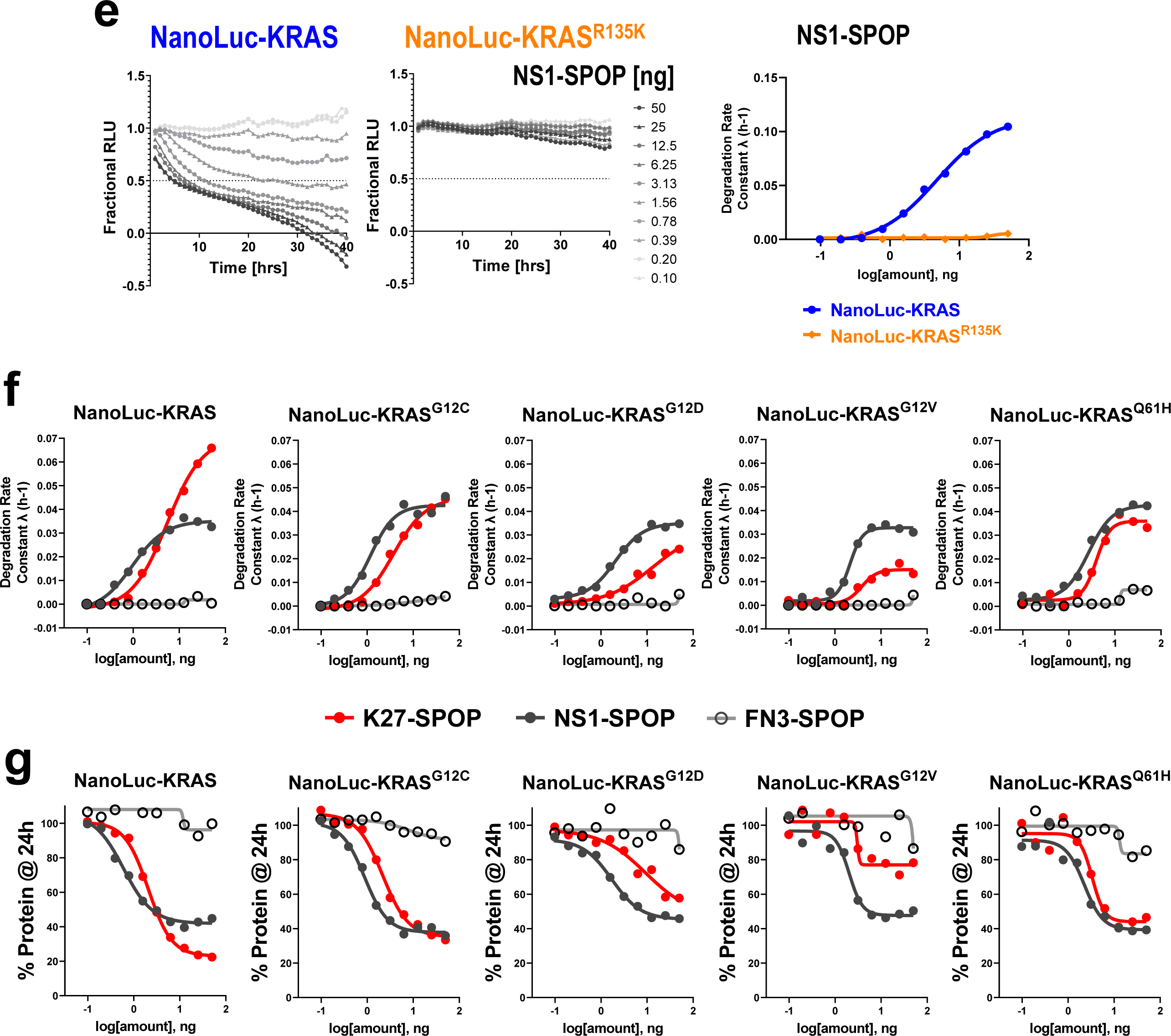
Real-time quantitative measurements of RAS degradation efficiency and selectivity using the NanoLuc assay. (**a**) Illustration of the NanoLuc degradation assay. T-REx™-293 cells with stable integration of NanoLuc-tagged RAS proteins under the control of a Tet-responsive promoter were generated. Expression was induced through a transient pulse of doxycycline, after which anti-RAS bioPROTACs were introduced through mRNA transfection. If successfully ubiquitinated and targeted for proteasomal degradation, the NanoLuc protein would not be available to react with its substrate (produced from the slow ester hydrolysis of Endurazine™) and the level of luminescence will drop. The rate of the decline in luminescence decline reflects the effectiveness of the transfected bioPROTAC. (**b**) Western blot analysis of T-REx™-293 stable cell lines as described in (**a**). Various concentrations of doxycycline (1, 10 and 100 ng/ml) were added to the culture media for 4 hours and protein lysates were collected. Fusion of a 19.7 kDa NanoLuc-tag to the RAS protein results in a slower migrating band when probed with pan-RAS antibodies. Activation of MAPK pathway was determined using the levels of phospho-MEK1/2 and phospho-ERK1/2. HSP90 was used as a loading control. (**c**) T-REx™-293 cells with doxycycline-induced expression of NanoLuc-HaloTag, NanoLuc-KRAS, NanoLuc-HRAS and NanoLuc-NRAS were transfected with a 10-point 2-fold dose-titration of the indicated bioPROTAC mRNA at time 0. Luminescence (RLU) was measured continuously every hour over a period of forty hours. Profiles were plotted as fractional RLU by normalizing to values of doxycycline induction with transfection reagent only (MAX) and no doxycycline (MIN). (**d**) Degradation rate calculated from (**c**) plotted against bioPROTAC amount in nanogram (ng). (**e**) Degradation profile and degradation rate calculated from T-REx™-293 cells with doxycycline-induced expression of NanoLuc-KRAS and NanoLuc-KRAS^R135K^, and transfected with a 10-point 2-fold dose-titration of NS1-SPOP mRNA at time 0. (**f**) Degradation rate calculated from T-REx™-293 cells with doxycycline-induced expression of various NanoLuc-tagged mutant KRAS and transfected with a 10-point 2-fold dose-titration of the indicated bioPROTAC mRNA at time 0. (**g**) Fractional RLU specifically retrieved for the 24 hours time-point from (**f**) and expressed as a percentage to represent the residual protein compared to transfection reagent only control.

HEK293 cells with stable integration of different NanoLuc-tagged RAS proteins were selected and varying concentrations of doxycycline were added to induce expression (**Fig. 5b**). Using a pan-RAS antibody, we noted that the overexpression of NanoLuc-tagged RAS proteins was significantly higher compared to endogenous levels (**Fig. 5b**). Interestingly, overexpression of NanoLuc-KRAS^G12D^ was sufficient to stimulate the MAPK pathway and result in increased phosphorylation of MEK and ERK (**Fig. 5b**). This was not observed with overexpression of the wild-type NanoLuc-RAS proteins (**Fig. 5b**), validating NanoLuc-KRAS^G12D^ as a functional and activating mutant protein.

To run this assay in a high-throughput 384-well format to accommodate a full dose-titration of bioPROTAC mRNAs, we first performed a series of optimization to select 1) type of live-cell substrate, 2) cell seeding densities, and 3) doxycycline concentrations and length of induction (**Supplementary Fig. 7**). With these conditions established, we chose K27-SPOP and NS1-SPOP as tools to evaluate if the NanoLuc assay can inform on the selectivity of bioPROTAC-mediated degradation. A previous report indicated that while K27 is specific for the GDP-loaded form of RAS (**Supplementary Fig. 2b**), it does not discriminate between RAS isoforms^34^. Accordingly, K27-SPOP degraded all RAS isoforms (NanoLuc-KRAS, NanoLuc-HRAS and NanoLuc-NRAS) in a dose-dependent manner, but not a control substrate NanoLuc-HaloTag (**Fig. 5c first panel**). Neither K27-SPOP_mut_ nor K27_mut_-SPOP degraded any of the NanoLuc-tagged proteins tested (**Fig. 5c second and third panel**). This suggested that the decline in luminescence is specific to the binding of NanoLuc-tagged substrate by an active bioPROTAC, which then induces its proteasomal turnover. Degradation rate, as described by Promega^41^, was calculated for each concentration and plotted (**Fig. 5d**). K27-SPOP was the most effective at degrading NanoLuc-KRAS, followed by NanoLuc-NRAS and finally NanoLuc-HRAS.

NS1 is a monobody that binds KRAS and HRAS, but not NRAS^33^ (**Supplementary Fig. 2a**). Using conventional Western blotting, the upper band corresponding to KRAS was preferentially lost in cells transfected with NS1-SPOP (**Fig. 2c lanes 6 and 8**). However, it was difficult to establish if other RAS isoforms were also affected since isoform-specific antibodies are lacking. Using the NanoLuc assay, it was clear that NS1-SPOP degraded NanoLuc-KRAS and NanoLuc-HRAS but not NanoLuc-NRAS (**Fig. 5c forth panel**), in line with its reported binding specificities^33^ (**Supplementary Fig. 2a**). When the substrate-binding domain of NS1-SPOP was replaced by the fibronectin type III domain (FN3), which forms the basis of the monobody scaffold, degradation was lost (**Fig. 5c fifth panel**). Interestingly, degradation rate constants suggested that NS1-SPOP degraded NanoLuc-KRAS more efficiently than NanoLuc-HRAS (**Fig. 5d forth panel**), despite the stronger affinity of NS1 for HRAS than for KRAS^33^ as determined from *in vitro* biophysical assays (**Supplementary Fig. 2a**). However, this result is consistent with the reported activity of the NS1 monobody in the cellular context, where it disrupted plasma membrane localization and RAF engagement for KRAS, but not for HRAS^33^. Overall, we have demonstrated that the NanoLuc assay is a useful tool to (1) inform on the specificity of degradation amongst closely related proteins and (2) provide quantitative measurements of degradation efficiencies inside live cells.

To further validate the NanoLuc assay, we generated a NanoLuc-KRAS^R135K^ stable cell line. R135 is a conserved residue in KRAS and HRAS but not NRAS, where it is instead a lysine. R135 makes extensive contacts with NS1 and is a major specificity determinant since its mutation to lysine greatly diminished NS1 binding^33^. Likewise, NS1-SPOP degraded NanoLuc-KRAS but was ineffective against NanoLuc-KRAS^R135K^ (**Fig. 5e**). This result clearly demonstrates how the specificity of degradation can be precisely controlled by the substrate-binding domain of bioPROTACs and the usefulness of the NanoLuc assay in providing this critical information in the cellular context.

While we have shown that the GDP-selective bioPROTAC K27-SPOP is able to degrade KRAS^G12D^ and reduce the viability of AsPC-1 cells (**Fig. 4**), it is not known if the same can be achieved with other oncogenic KRAS mutations. Specifically, it was reported that the intrinsic GTP hydrolysis rates are highly variable between KRAS mutants and therefore, the pool of GDP-loaded form available at a given time is expected to differ^42^. To determine if mutant KRAS does indeed cycle between the nucleotide-states at different rates, we generated NanoLuc-tagged lines of the most common KRAS mutations (G12C, G12D, G12V and Q61H) and compared their degradability by K27-SPOP (**Fig. 5f and Supplementary Fig. 8**). We expect that the higher the intrinsic hydrolysis rate, the greater the proportion of GDP-loaded mutant KRAS, and consequently the better the rate of degradation by K27-SPOP. NS1-SPOP was used as a normalizing comparator since it binds both the GTP- and GDP-loaded forms equally^33^. FN3-SPOP was used as a non-degrading control. Consistent with the nucleotide-state agnostic nature of NS1, the corresponding bioPROTAC NS1-SPOP degraded all five NanoLuc-tagged proteins with similar efficiencies (**Fig 5f black lines**). However, for K27-SPOP, a prominent difference in the rate of degradation was observed for each mutant (**Fig 5f red lines**). In accordance with the reported intrinsic hydrolysis rates^42^, K27-SPOP was the most effective against wildtype KRAS (even exceeding NS1-SPOP), followed by KRAS^G12C^, KRAS^G12D^ and finally KRAS^G12V^. The same trend was reproduced when we plotted the percentage of NanoLuc-tagged proteins remaining at 24 hours post-transfection of respective bioPROTAC mRNAs (**Fig 5g red lines**). KRAS^G12V^ was barely degraded by K27-SPOP while it was degraded by NS1-SPOP to a similar extent as the other mutants. One notable exception was KRAS^Q61H^. Although it was reported that Q61L and Q61H mutants exhibit the lowest intrinsic hydrolysis rates^42^, NanoLuc-KRAS^Q61H^ continued to be degraded by K27-SPOP (**Fig 5f and 5g last column**). It is currently unclear what accounts for this discrepancy.

During the preparation of this manuscript, there was a report of a KRAS-specific DARPin, K19^43^ (**Supplementary Fig. 9**). Specificity was conferred through extensive interactions with histidine 95, a residue that is unique to KRAS. We generated the K19-SPOP bioPROTAC and confirmed that it was only able to degrade NanoLuc-KRAS (and KRAS^G12D^) but not NanoLuc-HRAS and NanoLuc-NRAS (**Fig 6a**). By replacing histidine at position 95 with glutamine that is found in HRAS or leucine that is found in NRAS, K19-SPOP was no longer able to bind to and therefore degrade NanoLuc-KRAS^H95Q^ and NanoLuc-KRAS^H95L^, while its counterpart K27-SPOP continued to degrade all proteins (**Fig 6b and 6c, Supplementary Fig. 10**). Since K19 interacted with KRAS independently of the nucleotide-state^43^, K19-SPOP degraded the various NanoLuc-tagged KRAS mutants to a similar extent (**Fig 6d and Supplementary Fig. 10**). This result highlights how bioPROTACs that specifically degrade KRAS can be rapidly generated by engineering KRAS selectivity in the substrate-binding domain.

**Figure 6.**
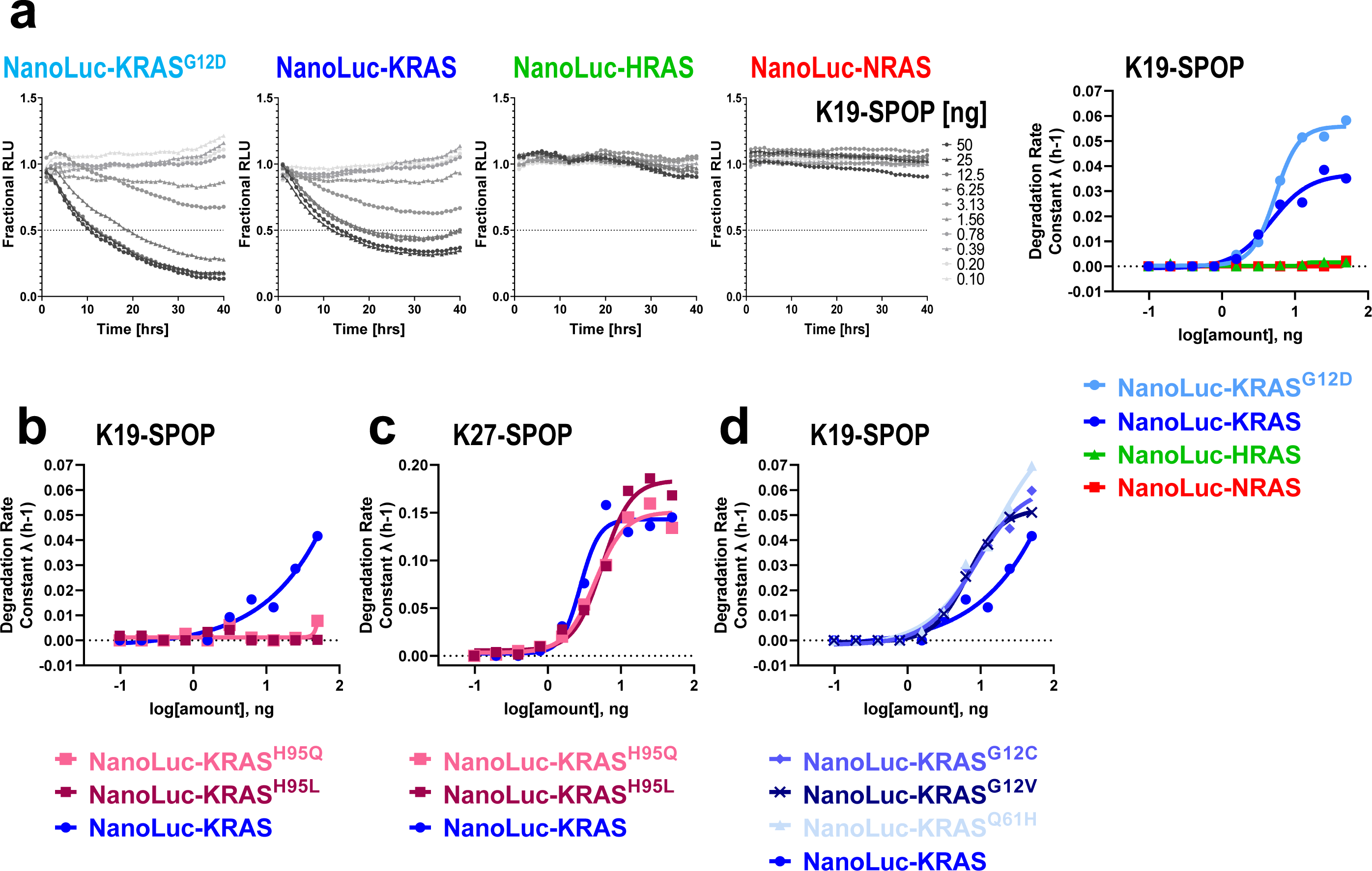
Characterization of a KRAS-specific bioPROTAC, K19-SPOP. (**a–d**) Degradation profile and degradation rate calculated from T-REx™-293 cells with doxycycline-induced expression of the indicated NanoLuc-tagged RAS protein and transfected with a 10-point 2-fold dose-titration of K19-SPOP (**a, b and d**) or K27-SPOP (**c**) mRNA at time 0.

## DISCUSSION

The work described herein advances our understanding of KRAS degradability and provides a compelling example of applying bioPROTACs as novel biological tools.

### Specificity of anti-RAS bioPROTACs

Prior to discussing how this work informs on i) the degradability of KRAS proteins and ii) KRAS biology, it is important to establish the specificity of the anti-KRAS bioPROTAC tools. As noted, the library of anti-RAS bioPROTACs were discovered using previously reported RAS binders spanning different affinities, isoform specificities, and nucleotide-state selectivities (**Fig. 2a, Supplementary Fig. 2**). Remarkably, all constructs, except for the K55-based bioPROTAC, resulted in functional degradation of endogenous RAS proteins (**Fig. 2c)**. The expected specificities of these bioPROTACs were also observed, with the clearest examples coming from the NanoLuc-RAS panel of cell lines. For example, NS1-SPOP was only able to degrade KRAS and HRAS but not NRAS (**Fig. 5c and 5d**). When a single specificity-determining residue on KRAS was mutated to the corresponding NRAS residue (KRAS^R135K^, **Fig. 5e**), it was no longer recognized by and thus cannot be degraded by NS1-SPOP, highlighting how bioPROTAC-mediated degradation is driven by precise biomolecular interactions. This point was further underscored using K19-SPOP, which was able to degrade KRAS but not HRAS nor NRAS (**Fig. 6a)**, as expected based on K19 binding specificities^43^ (**Supplementary Fig. 9**). X-ray structures have shown that the KRAS specificity of K19 is governed by its interaction with histidine 95, a residue where the equivalent amino acid is glutamine and leucine in HRAS and NRAS respectively. As predicted, K19-SPOP failed to degrade the KRAS^H95Q^ and KRAS^H95L^ point mutants (**Fig. 6b)**. The ability to engineer exquisite specificities, coupled with their ease of discovery, makes bioPROTACs valuable research tools.

### RAS degradability

To gain rapid insights into KRAS degradation, we used GFP-KRAS and the toolbox of anti-GFP bioPROTACs we developed in previous work^16^. Robust degradation was seen with most constructs (**Fig. 1c**). Amongst the E3 ligases achieving significant degradation was VHL, an important result as VHL ligands have been used extensively for small-molecule based PROTACs^7^ and therefore implies that they could be leveraged for degrading KRAS as well. Indeed, during the preparation of this manuscript, two relevant pre-print reports were released^44,45^. First, a KRAS-directed bioPROTAC was constructed employing full-length VHL fused to NS1 (which they termed VHL-aHRAS)^44^. This so-called Affinity-directed PROtein Missile (AdPROM) achieved some knockdown in A549^GFPKRAS^ cells but unfortunately did not yield significant growth inhibition in the three cancer cell lines tested – A549, HT29 and SW620. In the present study, we specifically removed the natural substrate-binding domain of VHL and demonstrated that it was highly effective at degrading both GFP-KRAS (when fused to vhhGFP4, **Fig. 1c**) and endogenous KRAS (when fused to K27 and R11.1.6, **Supplementary Fig. 3, lanes 16 and 18**). Notably, the other two RAS binders, NS1 and RBD-CRD, that had worked in combination with SPOP (**Supplementary Fig. 3, lanes 4 and 10**) failed to degrade KRAS when conjugated to VHL (**Supplementary Fig. 3, lanes 14 and 20**), suggesting that not all binder and E3 ligase combinations will produce active bioPROTACs. The second pre-print contribution^45^ appears to confirm that small molecule PROTACs which couple G12C covalent inhibitors to VHL ligands can achieve KRAS^G12C^ degradation. The current DC_50_ value (concentration to achieve 50% maximal degradation) stands at the micro-molar range. Indeed, employment of the VHL E3 ligase in a degradation strategy is a convenient starting point as PROTAC-compatible ligands are available. However, our study also uncovered other E3 ligases that gave superior GFP-KRAS degradation (**Fig. 1**), suggesting that time spent generating ligands to alternative E3 ligases could potentially yield more effective small molecule degraders.

While the case for converting an irreversible covalent inhibitor into a PROTAC molecule may not be immediately compelling, this seminal work by the Crews lab^45^ provides solid evidence for the degradation of oncogenic KRAS^G12C^ through a PROTAC approach and paves the way for future exploration in this direction. However, it is paramount to understand if the same can be applied to other KRAS mutants as they behave quite differently, both in terms of protein dynamics^46^ and ultimately, *in vivo* tumorgenicity^47,48^. Specifically, it was reported that the intrinsic GTP hydrolysis of various KRAS mutants differs in magnitude with the G12C mutant protein retaining the highest capacity to convert from the GTP-bound to the GDP-bound state^42^. The two nucleotide-states adopt distinct conformations and interact differently with the lipid bilayer^49^, which may impact PROTAC accessibility and ternary complex formation. More importantly, the binding pocket bound by covalent inhibitors is only accessible in the GDP-loaded state. Thus, PROTAC strategies that aim to (non-covalently) exploit this pocket might be limited to KRAS mutant proteins that retain sufficiently high GTPase activity. In this study, we further investigated the degradability of KRAS under different nucleotide states and containing different oncogenic mutations. By applying bioPROTACs that are either GDP-specific (K27-SPOP) or GTP-specific (RBD-CRD-SPOP), we have demonstrated that both nucleotide-states of K-, N-, and H-RAS are degradable substrates (**Fig. 2c and 5c**). We have also shown that wild-type and a spectrum of KRAS mutants (G12D, G12C, G12V, and Q61H) are degradable (**Fig 5f and 5g**).

### Cellular prevalence of the GDP-loaded state

The specificity of K27-SPOP for the GDP-loaded state of RAS has provided us with an ideal tool to probe the prevalence of the inactive state in individual KRAS mutants. The corresponding data adds to a growing body of literature challenging the dogma that oncogenic RAS proteins are “locked” in the GTP-state^50^. Instead, a more accurate view is one where the oncogenic mutations bias RAS to the GTP-state. In particular, biochemical studies have suggested that while phenotypic RAS mutations greatly compromised GAP-mediated hydrolysis of GTP, low levels of both GAP-mediated and intrinsic hydrolysis still occur, albeit with a range of rate constants across the different mutations^42^. Amongst them, KRAS^G12C^ had the highest intrinsic hydrolysis rate implying that a significant proportion of this protein may be present in the GDP-loaded (inactive) state. Indeed, the robust cellular activity demonstrated by G12C covalent inhibitors supports this notion since the corresponding binding pocket is only accessible in the GDP-state. In fact, the covalent inhibitors were able to capture more than 90% of KRAS^G12C^ proteins within one hour of treatment^4^, attesting to the significant rate of GTP hydrolysis in G12C mutants. We investigated the capacity of other KRAS mutants to cycle through the GDP/GTP states in the cellular context by using K27-SPOP as a gauge of the prevalence of the GDP-loaded state. K27-SPOP-induced degradability was WT > G12C > G12D > Q61H > G12V (from highest to lowest). Except for the Q61H mutant protein, this rank-order matches that determined previously^42^. Our study has thus corroborated the biochemical data with physiologically relevant cell-based readouts.

Previous studies have suggested that ≥75% KRAS occupancy is needed to achieve therapeutic efficacy in tumor models^51^. Irreversible inhibitory mechanisms have demonstrated the capacity to attain and sustain these levels despite the high intracellular concentration of KRAS (0.3 to 1.5 µM, **Supplementary Fig. 1**). However, for other KRAS mutants where a non-covalent inhibitor approach is required, achieving sufficient intracellular concentrations such that ≥75% stoichiometric target engagement is maintained will be challenging. A KRAS degradation approach is an attractive solution since PROTACs can potentially be recycled to catalyze multiple rounds of target degradation at sub-stoichiometric concentrations^6^. The binding pocket that is available in the GDP-state and bound by the G12C covalent inhibitors is an obvious starting point for the discovery of PROTAC molecules against other KRAS mutations. However, considering our current data and previous work^42^, leveraging this binding pocket for KRAS mutations with slower intrinsic hydrolysis may be challenging. As alternatives, our study has uncovered at least two additional RAS interfaces that might be leveraged for small molecule PROTAC strategies. The regions bound by NS1-SPOP and K19-SPOP are especially attractive since we have shown that degradation efficiencies are comparable regardless of KRAS mutational status (**Fig**.**5f and 6d**). Although the path towards the identification of small molecule ligands that bind to these sites remains challenging, our study has nonetheless shown definitively that PROTACs occupying these spaces do not obstruct poly-ubiquitination sites and proteasomal degradation of KRAS.

### Probing RAS dependency/Superiority of a degradation strategy

bioPROTACs can be used as a novel tool to probe for RAS dependency, with examples herein of (1) lack of dependency (despite complete pan-RAS degradation, HEK293 cells, **Fig. 3**) and (2) robust dependency (AsPC-1 cells, **Fig. 4**). Compared to protein knockdown using conventional siRNA where effects only occur after turnover of the pre-existing pool of proteins (for KRAS^G12C^, the reported half-life is ∼24 to 48 hours^4^), targeted protein degradation by bioPROTACs can be achieved within 4 hours following transfection (**Fig. 3a and 4a**). The present study also shows that a degradation modality outperforms the stoichiometric equivalent. For example, the bioPROTAC K27-SPOP demonstrated sustained pERK inhibition up to 24 hours post doxycycline induction, whereas pERK levels rebounded at this time point with the stoichiometric inhibitor K27-SPOP_mut_ despite its continued expression (**Fig. 3a**). It is likely that feedback mechanisms related to RAS re-activation are at play as have been reported elsewhere with inhibitors of the RAS-signaling pathway^52-54^. The superior effects of bioPROTACs were also recapitulated in functional assays where K27-SPOP resulted in complete growth arrest (**Fig. 4b**) and induction of apoptosis (**Fig. 4d**), whereas K27-SPOP_mut_ and the non-binding control K27_mut_-SPOP had no impact on AsPC-1 cells. Collectively, our study suggest that a degradation strategy can elicit a more comprehensive and durable inhibition of KRAS-dependent signaling compared to a stoichiometric approach, a finding that may have important implications for the treatment of KRAS mutant tumors.

## CONCLUDING REMARKS

This work advances the emerging field of bioPROTACs by demonstrating their specificity and utility as biological tools. Here, we have applied them to demonstrate the superiority of a degradation approach, inform on KRAS biology, and firmly establish the general degradability of RAS proteins across various isoforms, nucleotide-states, and mutant forms. This latter insight may prove useful in the design of small-molecule based degraders for KRAS, one of the most important oncogenic drivers. At the same time, this work highlights the potential therapeutic application of bioPROTACs and related intracellular biologics. Obtaining sufficient delivery and intracellular expression will be amongst the most important challenges. Encouragingly, the *in vivo* delivery of therapeutic mRNA is starting to be realized outside of the vaccine arena^55-60^.

## ACKNOWLEDGEMENTS

We thank Tomi K. Sawyer, Kaustav Biswas, Nicolas Boyer, Chunhui Huang, Alexander Stoeck, Nicole Boo, Jeff Chang, Sybil M. G. Williams, Payal Sheth, Jason E. Imbriglio, Uyen Phan, Ruban Mangadu, Mohammed Selman, CM Hsieh, Veronica Juan, Sara Zarnowski, Li Ding, Lei Chen, Amy C. Doty, Lauren A. Austin, Jeffrey S. Smith, Nicolas Solban, David P. Lane, Christopher J. Brown, Charles W. Johannes, Tsz Ying Yuen, Chandra Verma, Srinivasaraghavan Kannan, and all members of the Quantitative Biosciences team for helpful discussions and comments on the manuscript. The authors acknowledge support from the MRL Postdoctoral Research Program.

## Notes

### Competing Interest Statement

The authors have declared no competing interest.

