## Supplementry Information for "bioPROTACs establish RAS as a degradable target and provide novel RAS biology insights"

### Supplementary Figure 1

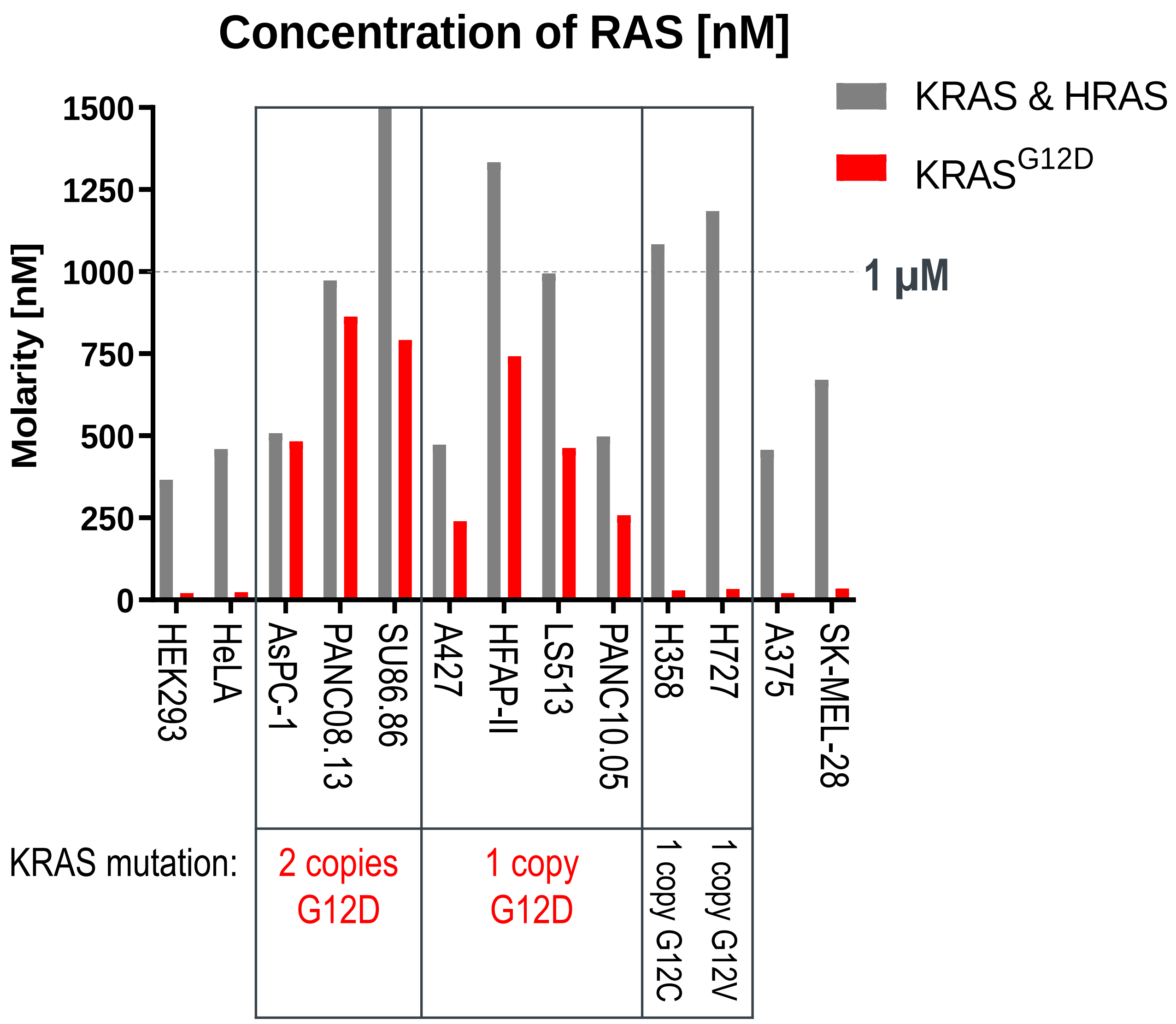


**Supplementary Figure 1| Quantitative of intracellular RAS concentration in a panel of KRAS mutant cell lines.** To prepare protein lysates, 50 µl of cell lysis buffer was added to every one million cells. Concentration was determined using the BCA assay and 2 µg lysate was loaded into Wes™ (ProteinSimple) together with a 5-point 4-fold concentration series of recombinant purified KRAS protein. By plotting a standard curve, we were able to determine the amount of RAS per µg protein for each cell type, and therefore the amount of RAS per cell. This value was converted into molarity, where the volume of each cell was calculated from its diameter measurements in suspension provided by Vi-CELL XR (Beckman Coulter).

### Supplementary Figure 2

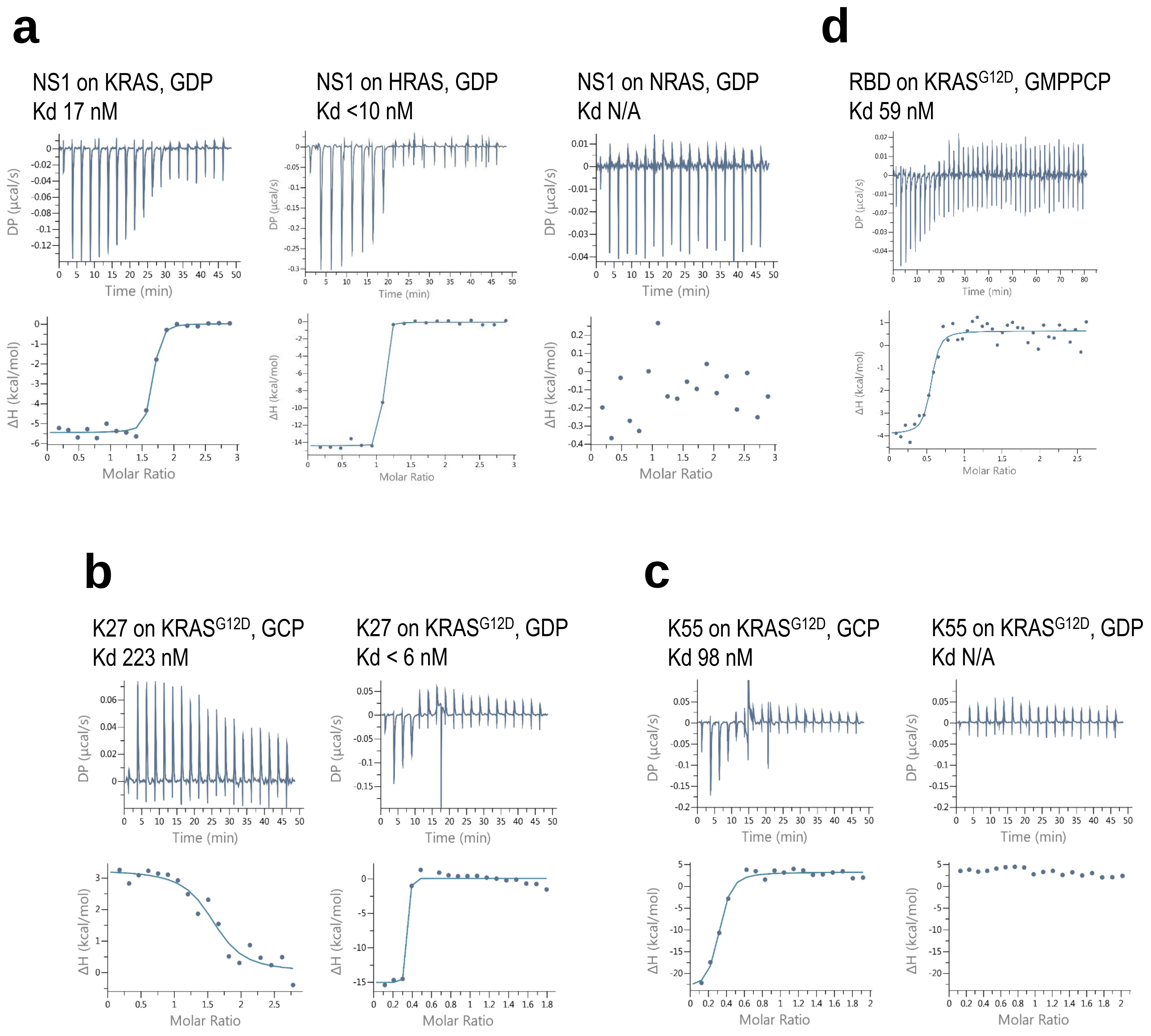


**Supplementary Figure 2| Isothermal titration calorimetric analysis (ITC).** ITC of the interaction of NS1 (a), K27 (b), K55 (c), and RBD (d) with the indicated RAS protein. Raw data (top) and binding isotherm (bottom) were obtained over a series of injections of the binder into the RAS protein. Differential power (μcal/sec) versus time (min) is presented in the form of integrated heat values. The data was fitted using a one binding site model and the calculated binding constant (Kd) is indicated.

### Supplementary Figure 3

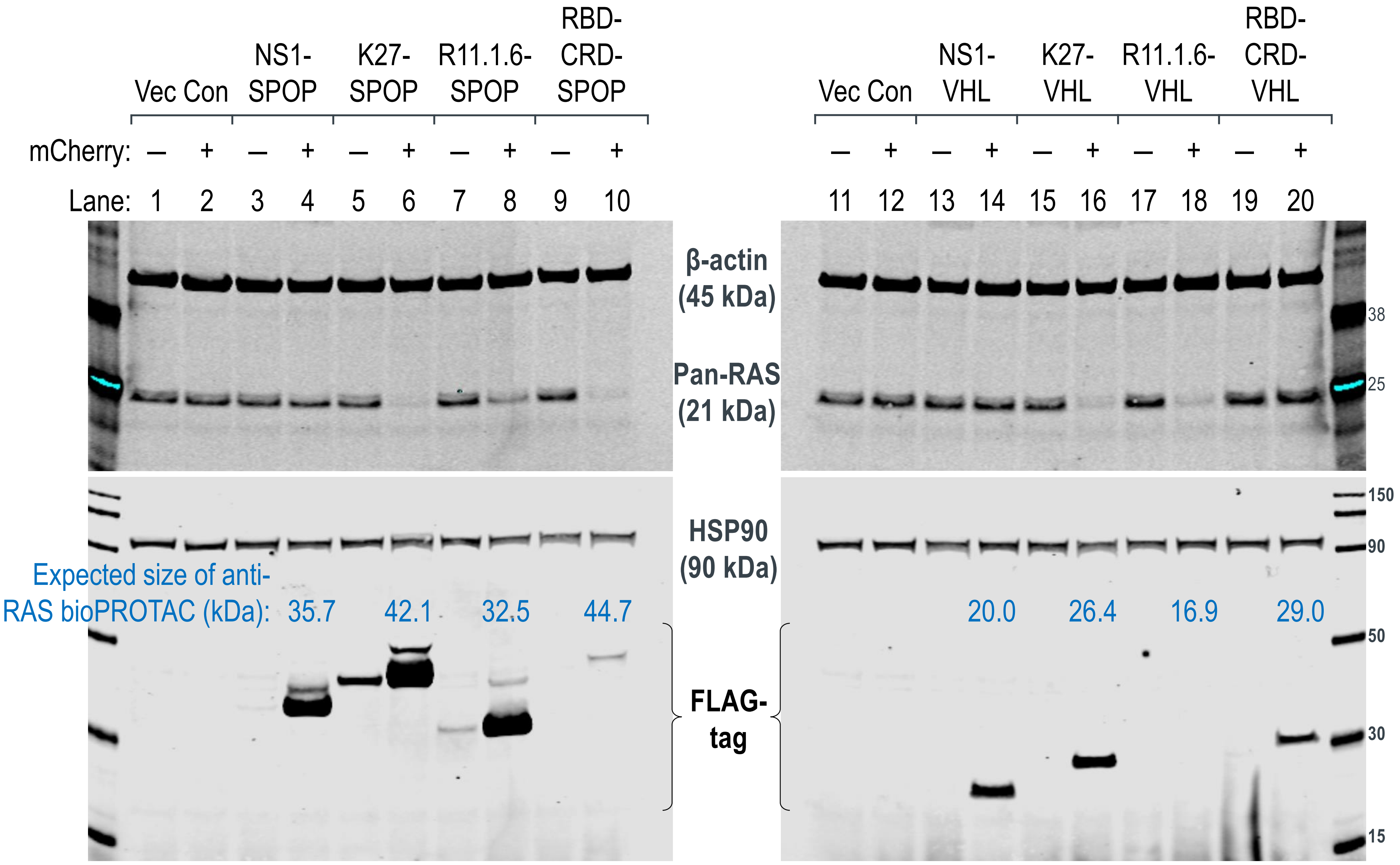


**Supplementary Figure 3| Degradation of endogenous RAS by SPOP- and VHL-based bioPROTACs.** Western blot analysis of HEK293 Tet-On® 3G cells transiently transfected with the indicated anti-RAS bioPROTACs and sorted according to the levels of mCherry (a marker of transfected cells) using FACS. Gating was set such that mCherry (-) cells have the same signal intensities as untreated cells in the mCherry channel, and anything above this basal level was assigned mCherry (+). In the pan-RAS blot, the upper band corresponds to KRAS while the lower band corresponds to HRAS and NRAS. Expression of the various anti-RAS bioPROTACs was detected using an anti-FLAG-tag antibody and the expected molecular weight of each chimeric protein is indicated in kilodaltons (kDa). β-actin and HSP90 were used as loading controls.

### Supplementary Figure 4

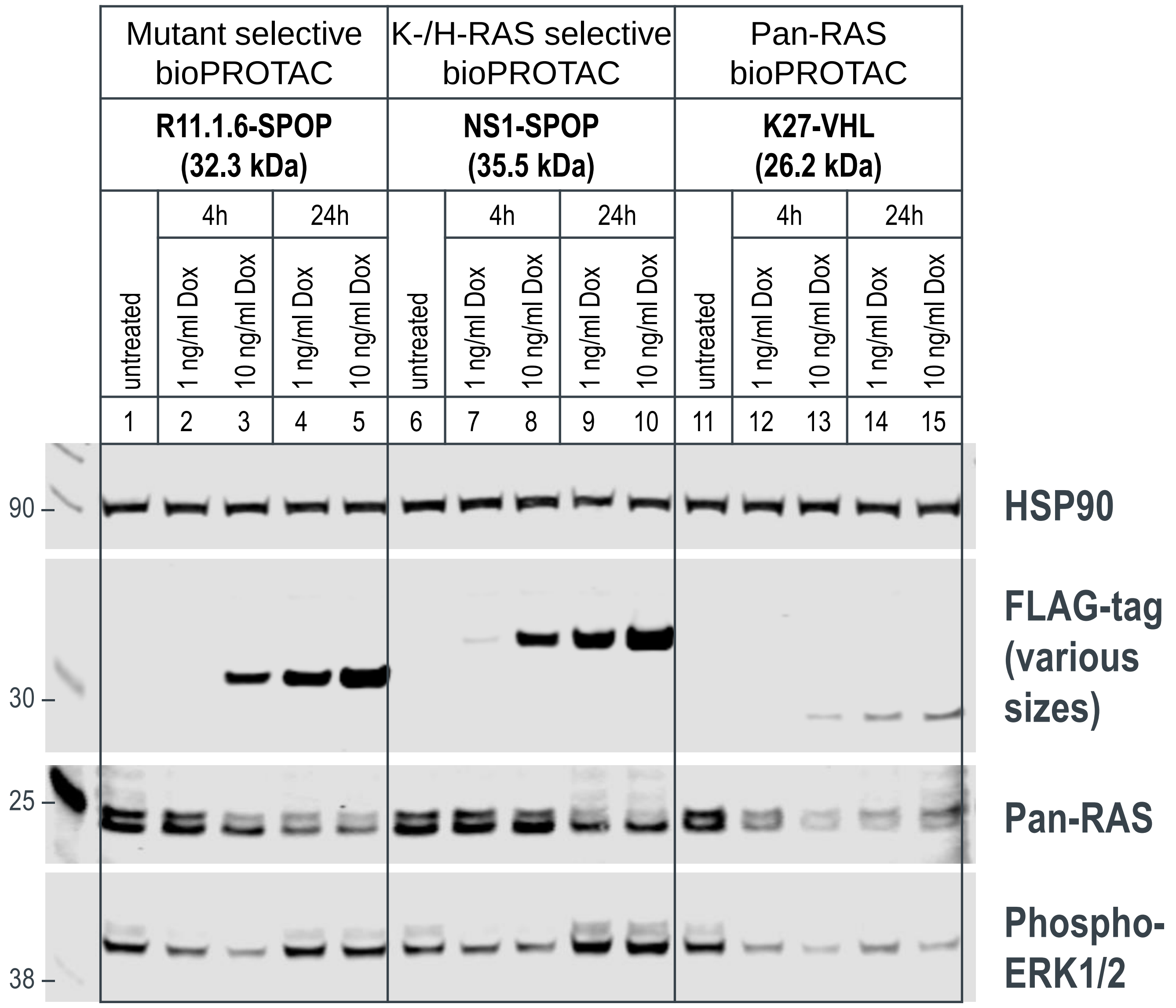


**Supplementary Figure 4| Degradation of endogenous RAS by doxycycline-inducible anti-RAS bioPROTACs.** Western blot analysis of T-REx™-293 cells with stable integration of R11.1.6-SPOP, NS1-SPOP or K27-VHL under the control of a Tet-responsive promoter. Various concentrations of doxycycline (1 or 10 ng/ml) were added to the culture media for the indicated length of time and protein lysates were collected. Degradation of RAS was detected using a pan-RAS antibody and disruption to the MAPK pathway was measured using the levels of phospho-ERK1/2. Expression of the various anti-RAS bioPROTAC was detected using an anti-FLAG-tag antibody. HSP90 was used as a loading control.

### Supplementary Figure 5

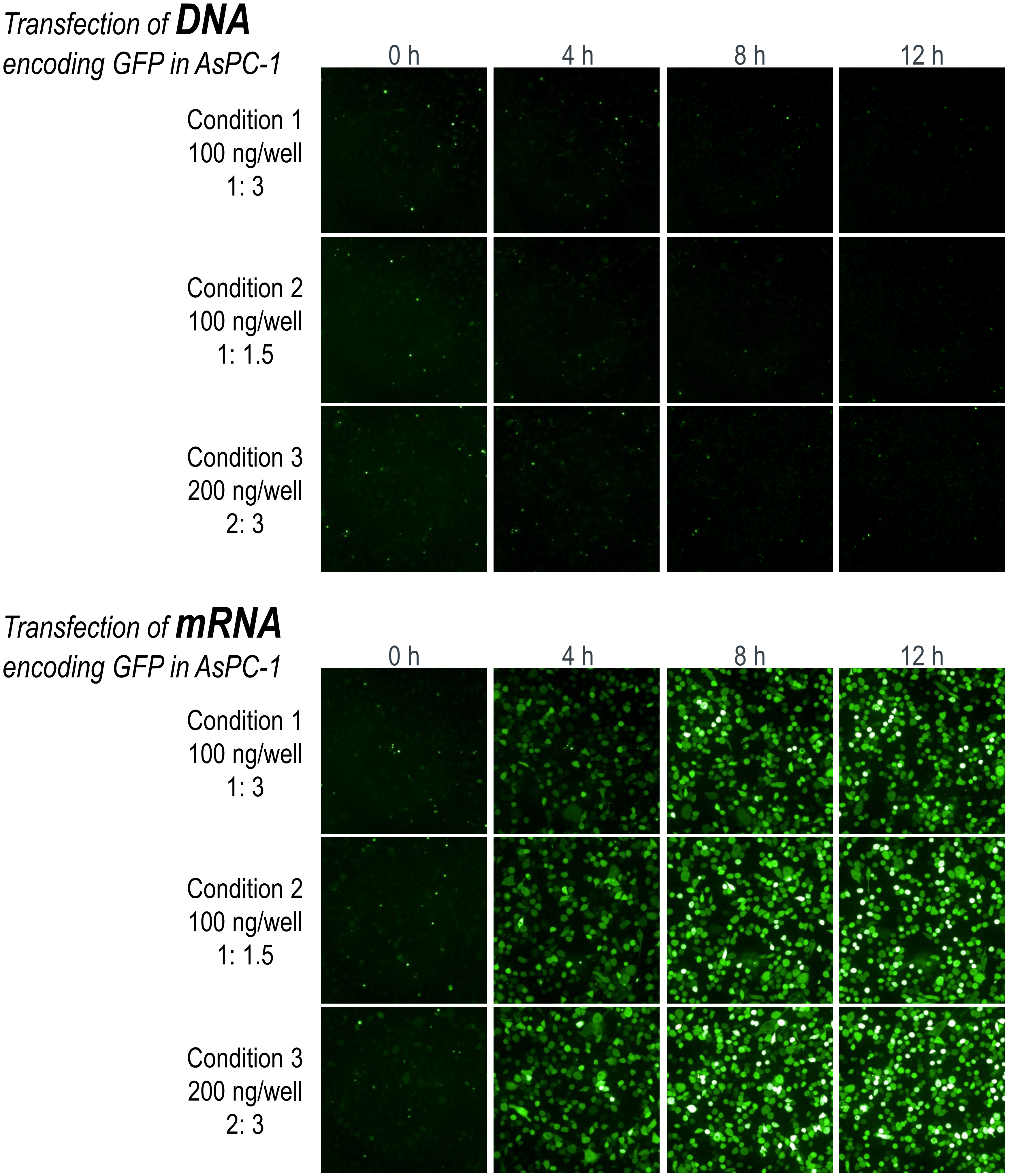


**Supplementary Figure 5| Comparison of DNA and mRNA transfection efficiencies in AsPC-1 cells.** Fluorescence images of GFP expression in AsPC-1 cells at the indicated time points following DNA or mRNA transfection. DNA transfection was performed using FuGENE® HD (Promega) and the ratio of DNA:transfection reagent is indicated. mRNA transfection was performed using Lipofectamine™ MessengerMAX™ (Life Technologies) and the ratio of mRNA:transfection reagent is indicated.

### Supplementary Figure 6

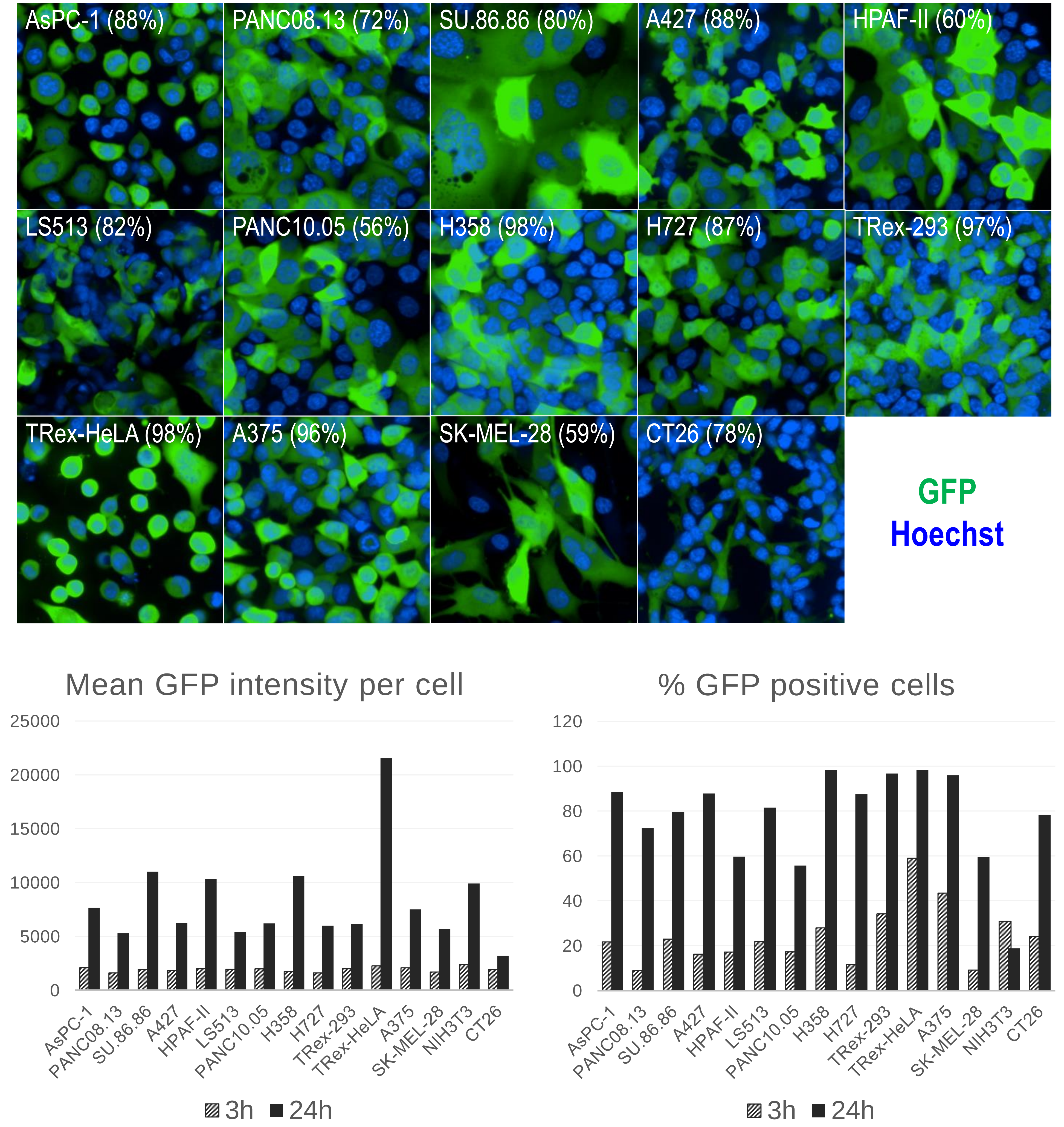


**Supplementary Figure 6| mRNA transfection is highly efficient across a panel of KRAS mutant cells.** Fluorescence images of GFP expression in the indicated cells 24 hours post-mRNA transfection. The number in brackets represent the percentage of GFP-positive cells 24 hours post-mRNA transfection. The mean GFP intensity per cell and percentage GFP-positive cells at 3 and 24 hours post-mRNA transfection were plotted.

### Supplementary Figure 7

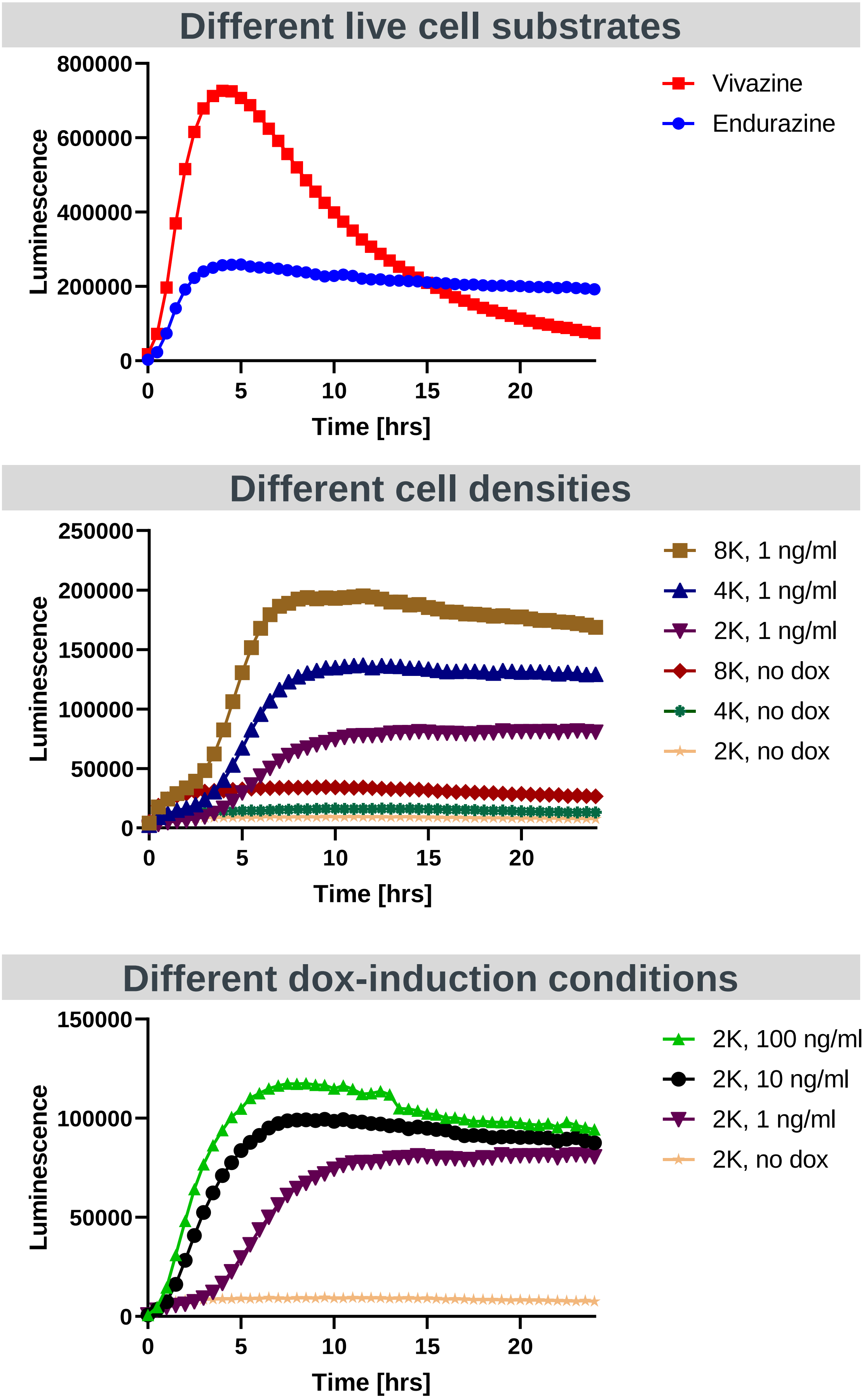


**Supplementary Figure 7| Optimization of the NanoLuc degradation assay.** Various conditions were tested for the optimization of the NanoLuc degradation assay, such as the choice of live-cell substrate, cell seeding densities and concentration of doxycycline. Raw luminescence values were plotted.

### Supplementary Figure 8

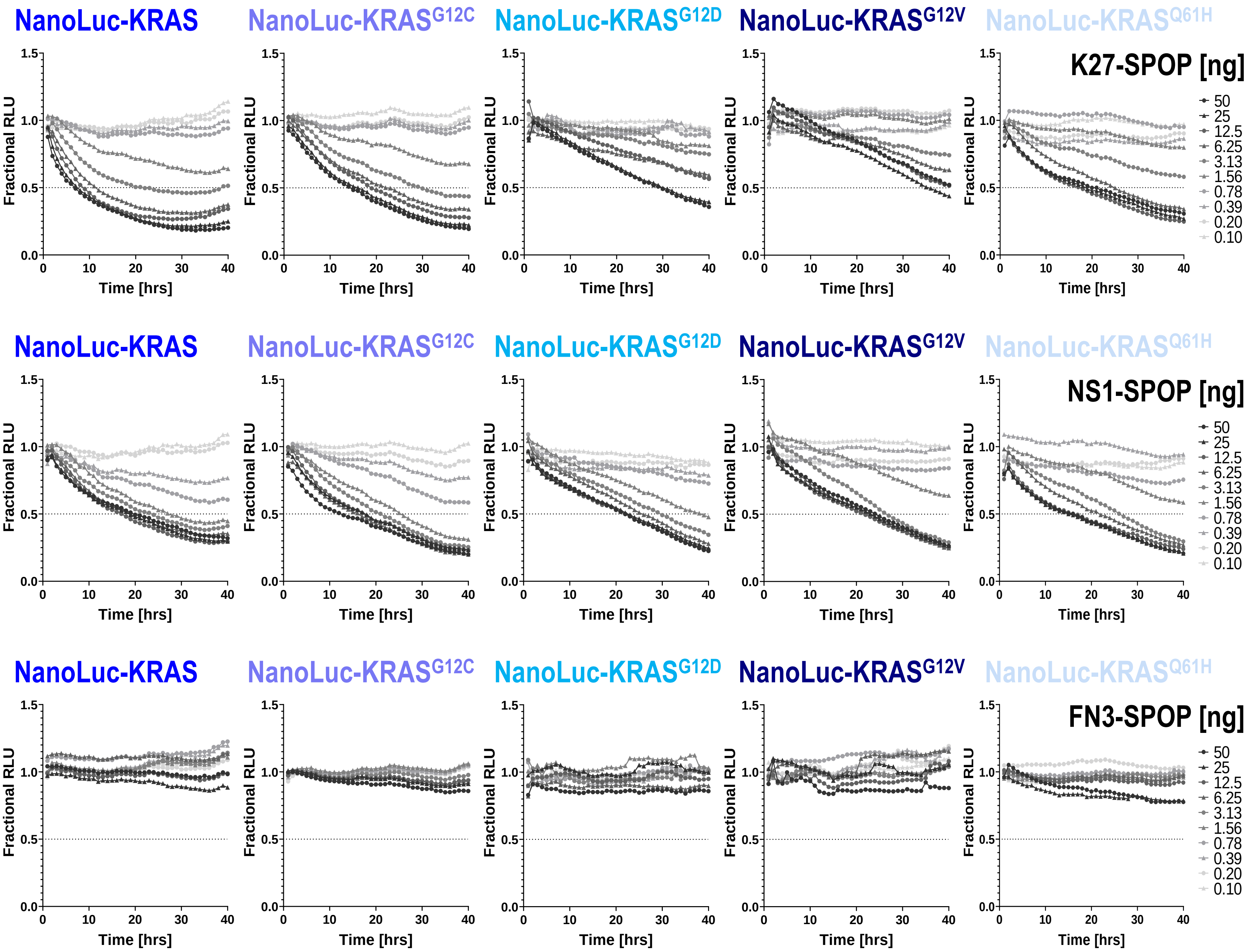


**Supplementary Figure 8| Some KRAS mutant forms are less efficiently degraded by K27-SPOP.** T-REx™-293 cells with doxycycline-induced expression of various NanoLuc-tagged mutant KRAS were transfected with a 10-point 2-fold dose-titration of the indicated bioPROTAC mRNA at time 0. Luminescence (RLU) was continuously measured at one hour intervals over a period of forty hours. Profiles were plotted as fractional RLU by normalizing to values of doxycycline induction with transfection reagent only (MAX) and no doxycycline (MIN).

### Supplementary Figure 9

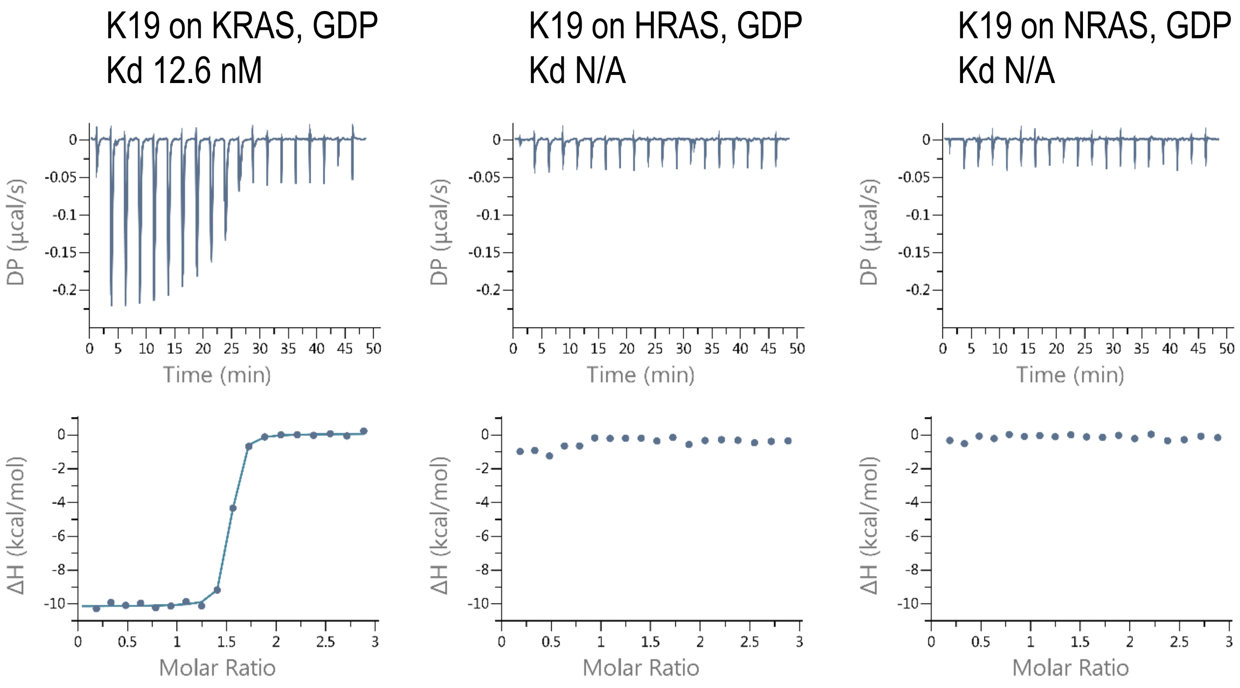


**Supplementary Figure 9| Isothermal titration calorimetric analysis (ITC).** ITC of the interaction of K19 with the indicated RAS protein. Raw data (top) and binding isotherm (bottom) were obtained over a series of injections of the binder into the RAS protein. Differential power (µcal/sec) versus time (min) is presented in the form of integrated heat values. The data was fitted using a one binding site model and the calculated binding constant (Kd) is indicated.

### Supplementary Figure 10

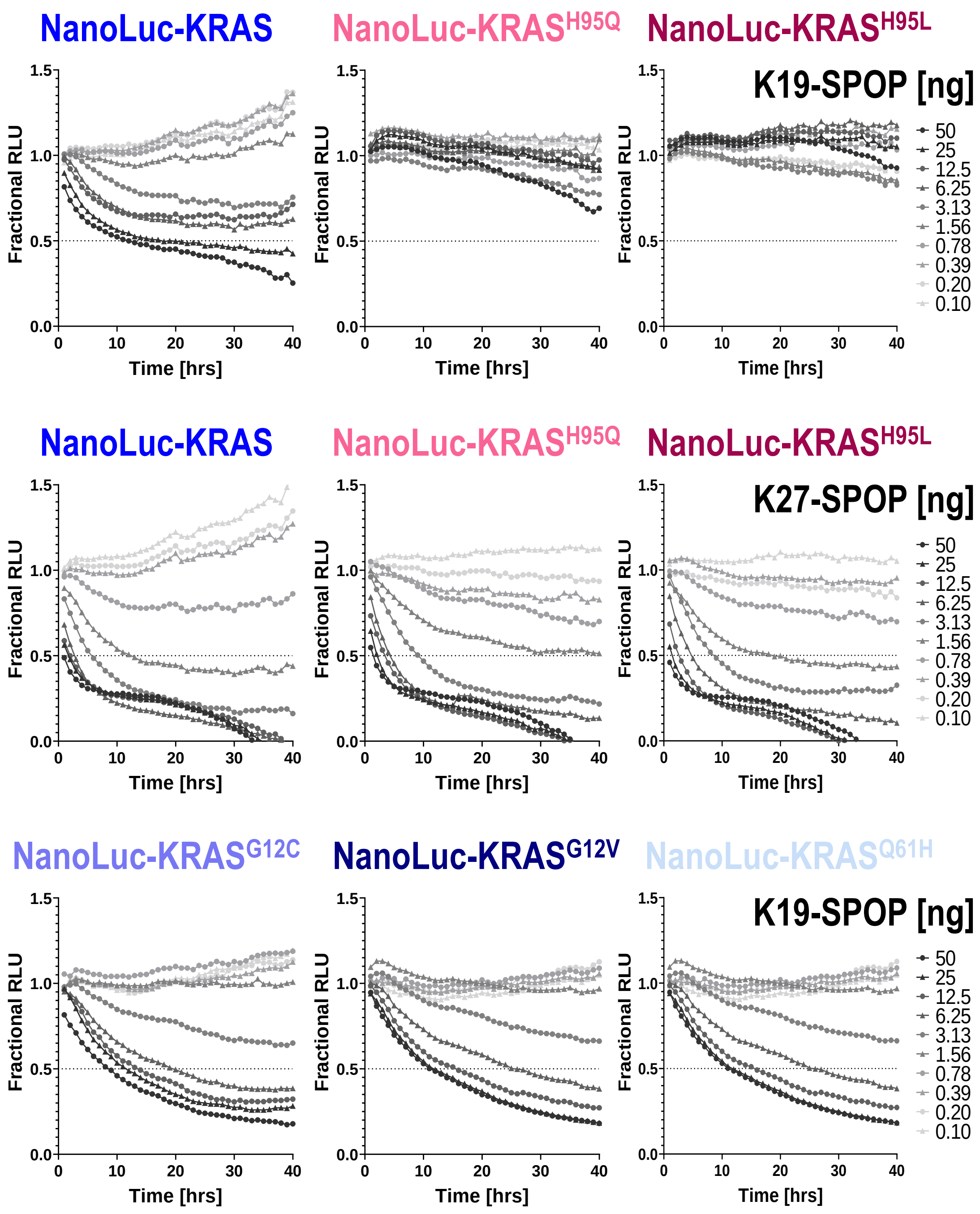


**Supplementary Figure 10| Characterization of the KRAS-specific bioPROTAC K19-SPOP.** T-REx™-293 cells with doxycycline-induced expression of the indicated NanoLuc-tagged KRAS protein were transfected with a 10-point 2-fold dose-titration of the indicated bioPROTAC mRNA at time 0. Luminescence (RLU) was continuously measured at one hour intervals over a period of forty hours. Profiles were plotted as fractional RLU by normalizing to values of doxycycline induction with transfection reagent only (MAX) and no doxycycline (MIN).

### MATERIALS AND METHODS

#### Plasmids

To generate Tet-On® 3G bidirectional inducible plasmids, gBlocks® gene fragments for the following inserts were synthesized by Integrated DNA technologies (IDT) and cloned into pTRE3G-BI-mCherry (Clontech) using BamHI (or BglII) and NotI: FLAG-vhhGFP4-$SPOP_{167-374}$, FLAG-$vhhGFP4_{mut}$-$SPOP_{167-374}$, FLAG-vhhGFP4-$SPOP_{mut}$, FLAG-β$TrCP_{2-263}$-vhhGFP4, FLAG-$FBW7_{2-293}$-vhhGFP4, FLAG-$SKP2_{2-147}$-vhhGFP4, FLAG-vhhGFP4-$VHL_{152-213}$, FLAG-$CRBN_{2-320}$-vhhGFP4, FLAG-$DDB2_{2-114}$-vhhGFP4, FLAG-vhhGFP4-$SOCS2_{143-198}$, FLAG-vhhGFP4-$ASB1_{266-335}$, FLAG-vhhGFP4-$CHIP_{128-303}$, FLAG-NS1-$SPOP_{167-374}$, FLAG-NS1-$SPOP_{mut}$, FLAG-FN3-$SPOP_{167-374}$, FLAG-K27-$SPOP_{167-374}$, FLAG-K27-$SPOP_{mut}$, FLAG-$K27_{mut}$-$SPOP_{167-374}$, FLAG-K55-$SPOP_{167-374}$, FLAG-R11.1.6-$SPOP_{167-374}$, FLAG-RBD-CRD-$SPOP_{167-374}$, FLAG-RBD-$SPOP_{167-374}$, FLAG-RBD-$SPOP_{mut}$, FLAG-NS1-$VHL_{152-213}$, FLAG-K27-$VHL_{152-213}$, FLAG-R11.1.6-$VHL_{152-213}$, FLAG-RBD-CRD-$VHL_{152-213}$ and FLAG-K19-$SPOP_{167-374}$. $vhhGFP4_{mut}$ lacks complementarity determining region 3 (ΔNVNVGFE). $SPOP_{mut}$ lacks the 3-box motif responsible for binding to Cullin (ΔAAEILILADLHSADQLKTQAVDFIN). $K27_{mut}$ is a non-binding control where 3 interfacial arginines were mutated to alanine. RBD and RBD-CRD corresponds to a.a. 52-131 and a.a. 52-220 of human RAF1 (NP_002871) respectively. FN3 is the $10^{th}$ fibronectin type III domain of human fibronectin (NP_001293058). For the establishment of stable cell lines with constitutive target gene expression, gBlocks for eGFP, eGFP-KRAS were synthesized by IDT and cloned into pEF6 (Thermo Fisher Scientific) using BamHI and NotI. For the establishment of stable cell lines with doxycycline-inducible target gene expression, gBlocks for FLAG-K27-$SPOP_{167-374}$, FLAG-K27-$SPOP_{mut}$, FLAG-$K27_{mut}$-$SPOP_{167-374}$, FLAG-R11.1.6-$SPOP_{167-374}$, FLAG-NS1-$SPOP_{167-374}$, FLAG-K27-$VHL_{152-213}$, NanoLuc-HaloTag, NanoLuc-KRAS, NanoLuc-HRAS, NanoLuc-NRAS, NanoLuc-$KRAS^{R135K}$, NanoLuc-$KRAS^{G12D}$, NanoLuc-$KRAS^{G12C}$, NanoLuc-$KRAS^{G12V}$, NanoLuc-$KRAS^{Q61H}$, NanoLuc-$KRAS^{H95Q}$ and NanoLuc-$KRAS^{H95L}$ were synthesized by IDT and cloned into pcDNA™4/TO (Thermo Fisher Scientific) using BamHI (or KpnI) and NotI. All plasmids were verified by sequencing at 1st BASE. Coding sequences of all constructs used in this study are provided in **Supplementary Table S1**.

#### Cell culture and transfection

HEK 293 Tet-On® 3G cells were purchased from Clontech and cultured in Minimum Essential Medium (MEM) GlutaMAX™ (Gibco) supplemented with 10% Tet system approved FBS (Clontech) and 100 µg/ml geneticin. T-REx™-293 cells were purchased from Thermo Fisher Scientific and cultured in MEM GlutaMAX™ supplemented with 10% Tet system approved FBS (Clontech) and 5 µg/ml blasticidin. Cells were seeded in poly-D-lysine coated plates and transfected with FuGENE® HD (Promega) for DNA plasmids or Lipofectamine™ MessengerMAX™ (Life Technologies) for mRNA the following day according to the manufacturer's protocol. To induce expression from the pTRE3G-BI-mCherry plasmids, 100 ng/ml doxycycline (Clontech) was added 24 hours post-transfection. To generate stable cell lines expressing GFP and GFP-KRAS, HEK 293 Tet-On® 3G cells were selected using 10 µg/ml blasticidin 3 days post-transfection and maintained in 5 µg/ml blasticidin once stable colonies are formed. GFP-positive cells were subsequently enriched by fluorescence-activated cell sorting (FACs) on BD FACSAria™ Fusion. To generate stable cell lines with doxycycline-inducible expression of FLAG-K27-$SPOP_{167-374}$, FLAG-K27-$SPOP_{mut}$, FLAG-$K27_{mut}$-$SPOP_{167-374}$, FLAG-R11.1.6-$SPOP_{167-374}$, FLAG-NS1-$SPOP_{167-374}$, FLAG-K27-$VHL_{152-213}$, NanoLuc-HaloTag, NanoLuc-KRAS, NanoLuc-HRAS, NanoLuc-NRAS, NanoLuc-$KRAS^{R135K}$, NanoLuc-$KRAS^{G12D}$, NanoLuc-$KRAS^{G12C}$, NanoLuc-$KRAS^{G12V}$, NanoLuc-$KRAS^{Q61H}$, NanoLuc-$KRAS^{H95Q}$ or NanoLuc-$KRAS^{H95L}$, T-REx™-293 cells were selected using 400 µg/ml Zeocin 3 days post-transfection and maintained in 200 µg/ml Zeocin once stable colonies are formed. All cells were maintained at 37ºC, 5% $CO_2$ and 90% relative humidity.

#### Flow cytometric analysis and fluorescence activated cell sorting (FACS)

Cells were seeded in 24-well poly-D-lysine coated plates and transfected as described above. 24 hours after transfection, the transfection media was removed and replaced with fresh media containing 100 ng/ml doxycycline. 24 hours after doxycycline-induction, cells were trypsinized and resuspended in cold PBS containing 10% FBS. The cell suspension was passed through a 35 µm nylon mesh to dissociate aggregates before analysis on BD LSRFortessa™ X-20. To sort cells according to mCherry or GFP expression, cells were seeded in 60 mm dishes and harvested in complete media after transfection and doxycycline-induction. A four-way sort was used on BD FACSAria™ Fusion to achieve a purity >98% and a yield >80%. 100, 000 cells were collected and processed for Western blot analysis or for further expansion in culture.

#### Fluorescence imaging and confluency measurements

Cells were seeded in 96-well poly-D-lysine coated µCLEAR® plates (Greiner) and allowed to attach overnight. The next day, transfection and doxycycline-induction were performed as described above. For immunostaining to detect apoptotic cells, cells were fixed in 4% formaldehyde in PBS for 15 minutes and blocked with 5% normal donkey serum-0.3% Triton™ X-100 in PBS for 1 hour at room temperature. Rabbit anti-cleaved caspase-3 (Asp175) antibody (Cell Signaling Technology, #9661) was diluted in 1% BSA-0.3% Triton™ X-100 in PBS and incubated overnight at 4°C. The next day, donkey anti-rabbit Alexa Fluor 488 (ThermoFisher A-21206) was added for 1 hour at room temperature. Nuclei were counterstained with Hoechst. Images were acquired using the Opera Phenix™ High Content Confocal Screening System under the 20X or 40X water immersion lenses. For live-cell imaging, chamber conditions were set to 37ºC, 5% $CO_2$. Percentage confluency of cells were tracked continuously using the IncuCyte® S3 Live-Cell Analysis System under the 4X whole well imaging objective and analyzed using the IncuCyte® software.

#### Isothermal Titration Calorimetry (ITC)

Recombinant proteins were synthesized and purified by Evotec or the Protein Production Platform (PPP) at NTU School of Biological Sciences. ITC measurements were carried out on a MicroCal PEAQ-ITC Automated System (Malvern Panalytical). Protein samples were dialyzed overnight at 4°C in buffer containing 1X PBS pH 7.4, 1 mM $MgCl_2$. 60 – 150 µM of NS1, K27, K55, RBD within the syringe were titrated into 6 – 10 µM concentrations of indicated RAS proteins in the sample cell. All binding experiments were carried out at constant temperature of 25°C. Data analysis was performed using MicroCal PEAQ-ITC Analysis Software and fitted with one site binding model.

**Western blot analysis**

Cells were lysed in ice-cold cell lysis buffer (Cell Signaling Technology) supplemented with 1 mM PMSF and cOmplete™ EDTA-free protease inhibitor cocktail (Roche) for 30 min with intermittent vortexing. Lysates were centrifuged at 18,000 g, 4°C for 15 min and supernatants were snap frozen in liquid nitrogen. Protein concentration was determined using the BCA protein assay kit (Pierce). For direct lysis, 100 µl of Bolt™ LDS sample buffer supplemented with NuPAGE® sample reducing agent was added per well of a 24-well plate. The wells were scrapped using wide orifice tips and the lysate was transferred into PCR-strip tubes and sonicated for 10 X 10 seconds in a chilled water bath sonicator (QSonica). 20 to 50 µg of protein extract was separated on 4-12% Bis-Tris plus gels, transferred onto nitrocellulose membranes using the Trans-Blot® Turbo™ semi-dry system (Bio-rad), and blocked for 1 hour at room temperature with tris-buffered saline (TBS) Odyssey blocking buffer (Li-Cor). Blots were probed with the appropriate primary antibodies overnight at 4°C in Odyssey blocking buffer supplemented with 0.1% Tween-20, followed by the secondary antibodies IRDye® 680RD donkey anti-mouse IgG or IRDye® 800CW donkey anti-rabbit IgG (Li-Cor) for 1 hour at room temperature. Fluorescent signals were imaged and quantified using Odyssey® CLx. Primary antibodies used were: pan-RAS (Cell Signaling Technology, #3339), β-actin (Santa Cruz Biotechnology, sc-8432), HSP90 (BD Transduction Laboratories, 610419), FLAG-tag (Cell Signaling Technology, #8146 and #14793), phospho-p44/42 MAPK (ERK1/2) (Thr202/Tyr204) (Cell Signaling Technology, #4370), p44/42 MAPK (ERK1/2) (Cell Signaling Technology, #4695), phospho-MEK1/2 (Ser217/221) (Cell Signaling Technology, #9154) and phospho-AKT (Ser473) (Cell Signaling Technology, #4060).

**Generation of modified mRNA**

mRNAs capped with CleanCap® and modified with 100% pseudouridine were either synthesized at TriLink Biotechnologies or *in vitro* transcribed using the mMESSAGE mMACHINE® T7 Ultra transcription kit (Ambion, AMB13455). Linearized plasmid DNA containing the target gene downstream of a T7 RNA polymerase promoter was used as the template and synthesis reactions were performed according to the manufacturer's protocol, except for substituting T7 2X NTP/ARCA with 8 mM CleanCap® Reagent AG (TriLink Biotechnologies, N-7113) and 10 mM each of pseudouridine-5'-triphosphate (TriLink Biotechnologies, N-1019), ATP, CTP and GTP.

mRNAs were subsequently purified by the RNeasy Mini Kit (Qiagen, 74104) and quantified on the NanoDrop spectrophotometer.

**NanoLuc degradation assay**

Stable cell lines expressing the various NanoLuc-tagged proteins were generated as described above. Cells were pulsed with 3 ng/ml or 10 ng/ml doxycycline for 2 hours to induce the expression of the respective NanoLuc-tagged protein. Lipofectamine™ MessengerMAX™ (Life Technologies) was diluted in opti-MEM to the desired working concentration and dispensed onto 384-well white assay plates (Greiner 781080). A source plate (Labcyte LP-0200) containing serial dilutions of the mRNAs was prepared using the Bravo liquid handler (Agilent) and a 10-point 2-fold dose-titration of each mRNA was dispensed onto the assay plate using Echo (Labcyte). After a 10 min incubation, cells with doxycycline washed-off were added followed by 20 μM Endurazine (Promega), an extended time-released live cell substrate. Luminescence was measured continuously at 1-hour intervals on the Tecan Spark 10M set to 37ºC, 5% $CO_2$.

**Supplementary Table S1.** Coding sequences of all constructs.

| | |
|---|---|
| FLAG-vhhGFP4-SPOP$_{167\text{-}374}$ | ATGGATTACAAGGACGACGACGACAAGgctagcGATCAAGTCCAACTGGTGGAGTCTGGTGGCGCT<br>TTGGTGCAGCCAGGTGGCTCTCTGCGTTTGTCCTGTGCCGCTTCTGGCTTCCCAGTGAACCGCTAT<br>TCCATGCGCTGGTATCGCCAGGCTCCAGGCAAAGAGCGTGAGTGGGTAGCCGGTATGTCCAGCGCG<br>GGTGATCGTAGCTCCTATGAAGACTCCGTGAAGGGCCGTTTCACCATCAGCCGTGACGATGCCCGT<br>AACACGGTGTATCTGCAAATGAACAGCTTGAAACCTGAAGATACGGCCGTGTATTACTGTAATGTG<br>AACGTGGGCTTCGAGTATTGGGGCCAAGGCACCCAGGTCACCGTCTCCAGCtccggaAGCGTGAAC<br>ATCTCCGGCCAGAACACAATGAACATGGTCAAGGTGCCCGAGTGCAGACTGGCCGACGAGCTGGGA<br>GGACTGTGGGAGAACTCCAGGTTTACCGACTGCTGCCTGTGCGTGGCCGGCCAAGAGTTCCAAGCC<br>CACAAAGCCATCCTGGCCGCTAGGTCCCCCGTGTTCAGCGCCATGTTCGAGCACGAGATGGAGGAG<br>TCCAAGAAGAACAGAGTGGAGATTAACGATGTGGAGCCCGAGGTGTTCAAAGAAATGATGTGCTTC<br>ATCTACACCGGCAAGGCCCCCAACCTGGATAAAATGGCCGATGACCTGCTGGCCGCCGCCGATAAG<br>TACGCCCTGGAGAGACTGAAGGTGATGTGCGAGGACGCTCTGTGTTCCAACCTGTCCGTGGAAAAT<br>GCCGCCGAGATCCTCATCCTGGCCGACCTGCATAGCGCCGACCAGCTGAAAACCCAGGCCGTGGAC<br>TTCATCAACTATCACGCTTCCGACGTGCTGGAGACCAGCGGATGGAAGAGCATGGTGGTGAGCCAT<br>CCCCATCTCGTGGCCGAAGCCTACAGGAGCCTGGCaAGCGCCCAGTGTCCCTTTCTGGGCCCTCCC<br>AGGAAGAGACTGAAACAGAGCTGA |
| FLAG-vhhGFP4$_{mut}$-SPOP$_{167\text{-}374}$ | ATGGATTACAAGGACGACGACGACAAGgctagcGATCAAGTCCAACTGGTGGAGTCTGGTGGCGCT<br>TTGGTGCAGCCAGGTGGCTCTCTGCGTTTGTCCTGTGCCGCTTCTGGCTTCCCAGTGAACCGCTAT<br>TCCATGCGCTGGTATCGCCAGGCTCCAGGCAAAGAGCGTGAGTGGGTAGCCGGTATGTCCAGCGCG<br>GGTGATCGTAGCTCCTATGAAGACTCCGTGAAGGGCCGTTTCACCATCAGCCGTGACGATGCCCGT<br>AACACGGTGTATCTGCAAATGAACAGCTTGAAACCTGAAGATACGGCCGTGTATTACTGTTATTGG<br>GGCCAAGGCACCCAGGTCACCGTCTCCAGCtccggaAGCGTGAACATCTCCGGCCAGAACACAATG<br>AACATGGTCAAGGTGCCCGAGTGCAGACTGGCCGACGAGCTGGGAGGACTGTGGGAGAACTCCAGG<br>TTTACCGACTGCTGCCTGTGCGTGGCCGGCCAAGAGTTCCAAGCCCACAAAGCCATCCTGGCCGCT<br>AGGTCCCCCGTGTTCAGCGCCATGTTCGAGCACGAGATGGAGGAGTCCAAGAAGAACAGAGTGGAG<br>ATTAACGATGTGGAGCCCGAGGTGTTCAAAGAAATGATGTGCTTCATCTACACCGGCAAGGCCCCC<br>AACCTGGATAAAATGGCCGATGACCTGCTGGCCGCCGCCGATAAGTACGCCCTGGAGAGACTGAAG<br>GTGATGTGCGAGGACGCTCTGTGTTCCAACCTGTCCGTGGAAAATGCCGCCGAGATCCTCATCCTG<br>GCCGACCTGCATAGCGCCGACCAGCTGAAAACCCAGGCCGTGGACTTCATCAACTATCACGCTTCC<br>GACGTGCTGGAGACCAGCGGATGGAAGAGCATGGTGGTGAGCCATCCCCATCTCGTGGCCGAAGCC<br>TACAGGAGCCTGGCaAGCGCCCAGTGTCCCTTTCTGGGCCCTCCCAGGAAGAGACTGAAACAGAGC<br>TGA |
| FLAG-vhhGFP4-SPOP$_{mut}$ | ATGGATTACAAGGACGACGACGACAAGgctagcGATCAAGTCCAACTGGTGGAGTCTGGTGGCGCT<br>TTGGTGCAGCCAGGTGGCTCTCTGCGTTTGTCCTGTGCCGCTTCTGGCTTCCCAGTGAACCGCTAT<br>TCCATGCGCTGGTATCGCCAGGCTCCAGGCAAAGAGCGTGAGTGGGTAGCCGGTATGTCCAGCGCG<br>GGTGATCGTAGCTCCTATGAAGACTCCGTGAAGGGCCGTTTCACCATCAGCCGTGACGATGCCCGT<br>AACACGGTGTATCTGCAAATGAACAGCTTGAAACCTGAAGATACGGCCGTGTATTACTGTAATGTG<br>AACGTGGGCTTCGAGTATTGGGGCCAAGGCACCCAGGTCACCGTCTCCAGCtccggaAGCGTGAAC<br>ATCTCCGGCCAGAACACAATGAACATGGTCAAGGTGCCCGAGTGCAGACTGGCCGACGAGCTGGGA<br>GGACTGTGGGAGAACTCCAGGTTTACCGACTGCTGCCTGTGCGTGGCCGGCCAAGAGTTCCAAGCC<br>CACAAAGCCATCCTGGCCGCTAGGTCCCCCGTGTTCAGCGCCATGTTCGAGCACGAGATGGAGGAG<br>TCCAAGAAGAACAGAGTGGAGATTAACGATGTGGAGCCCGAGGTGTTCAAAGAAATGATGTGCTTC<br>ATCTACACCGGCAAGGCCCCCAACCTGGATAAAATGGCCGATGACCTGCTGGCCGCCGCCGATAAG<br>TACGCCCTGGAGAGACTGAAGGTGATGTGCGAGGACGCTCTGTGTTCCAACCTGTCCGTGGAAAAT<br>TATCACGCTTCCGACGTGCTGGAGACCAGCGGATGGAAGAGCATGGTGGTGAGCCATCCCCATCTC<br>GTGGCCGAAGCCTACAGGAGCCTGGCaAGCGCCCAGTGTCCCTTTCTGGGCCCTCCCAGGAAGAGA<br>CTGAAACAGAGCTGA |
| FLAG-βTrCP$_{2\text{-}263}$-vhhGFP4 | ATGggaggccttGATTACAAGGACGACGACGACAAGgctagcGATCCAGCCGAGGCTGTGTTGCAA<br>GAGAAGGCCCTTAAGTTCATGAATAGCAGCGAGCGCGAGGATTGTAACAATGGTGAGCCTCCTAGA<br>AAAATCATTCCAGAGAAGAACAGCCTCAGACAAACATACAATAGTTGCGCCCGGCTTTGCTTGAAC |

| | |
|---|---|
| | CAAGAAACTGTCTGTCTTGCATCTACCGCTATGAAGACTGAAAACTGCGTCGCCAAAACCAAACTG<br>GCGAATGGCACGTCTTCCATGATTGTCCCCAAACAGAGGAAACTGAGTGCTTCCTATGAAAAAGAA<br>AAAGAGCTGTGTGTAAAATACTTTGAACAGTGGTCAGAAAGTGACCAAGTCGAGTTTGTTGAACAT<br>TTGATCTCACAAATGTGCCATTACCAACATGGACACATCAATTCTTATTTGAAGCCTATGCTCCAG<br>CGAGATTTTATCACGGCTCTGCCTGCCAGGGGGCTGGATCACATTGCGGAGAATATCCTCTCATAT<br>CTCGACGCAAAGTCCCTTTGTGCAGCCGAGCTGGTCTGCAAAGAATGGTATAGGGTGACATCTGAT<br>GGGATGCTTTGGAAGAAACTTATTGAACGAATGGTGCGGACAGACTCCCTGTGGCGCGGGCTGGCA<br>GAAAGGCGCGGATGGGGCCAGTATCTGTTCAAGAACAAACCGCCTGATGGGAACGCGCCGCCAAAC<br>AGCTTCTACCGGGCACTGTACCCCAAAATCATACAGGATATTGAAACTATTGAAAGCAATTGGCGA<br>TGTGGCCGCCATAGTCTGCAACGCATACACTGCCGGAGCGGGTCAGGTAGTGGCtccggaGATCAA<br>GTCCAACTGGTGGAGTCTGGTGGCGCTTTGGTGCAGCCAGGTGGCTCTCTGCGTTTGTCCTGTGCC<br>GCTTCTGGCTTCCCAGTGAACCGCTATTCCATGCGCTGGTATCGCCAGGCTCCAGGCAAAGAGCGT<br>GAGTGGGTAGCCGGTATGTCCAGCGCGGGTGATCGTAGCTCCTATGAAGACTCCGTGAAGGGCCGT<br>TTCACCATCAGCCGTGACGATGCCCGTAACACGGTGTATCTGCAAATGAACAGCTTGAAACCTGAA<br>GATACGGCCGTGTATTACTGTAATGTGAACGTGGGCTTCGAGTATTGGGGCCAAGGCACCCAGGTC<br>ACCGTCTCCAGCggatccTGA |
| FLAG-FBW7$_{2-293}$-vhhGFP4 | ATGggaggccttGATTACAAGGACGACGACGACAAGgctagcTGCGTGCCTCGCTCTGGGCTCATA<br>TTGTCCTGTATTTGCCTGTACTGCGGCGTTCTGCTCCCAGTACTGCTCCCAAATCTTCCATTCTTG<br>ACCTGTTTGTCCATGTCAACGCTTGAGTCTGTCACGTACTTGCCTGAGAAAGGGCTGTATTGCCAA<br>AGGCTGCCCTCTAGTCGAACCCATGGTGGTACGGAATCCCTGAAAGGCAAGAACACCGAGAACATG<br>GGCTTTTATGGACCTTGAAAATGATTTTCTACAAGATGAAAAGAAAACTTGACCATGGATCAGAG<br>GTGCGGAGCTTCAGCTTGGGTAAAAAACCCTGCAAAGTATCTGAATACACGTCTACGACCGGGTTG<br>GTTCCCTGTTCCGCCACTCCAACCACTTTCGGTGATTTGAGAGCAGCAAACGGCCAAGGGCAACAG<br>CGACGGCGCATTACTTCAGTGCAGCCTCCAACGGGGCTTCAGGAGTGGTTGAAGATGTTTCAGAGC<br>TGGAGCGGTCCAGAAAAGCTGCTTGCTCTTGATGAGCTGATTGATTCTTGTGAACCCACTCAAGTC<br>AAACACATGATGCAGGTTATTGAGCCTCAGTTTCAACGGGATTTTATCAGCCTTCTTCCTAAGGAA<br>TTGGCACTGTACGTCCTGTCTTTCCTCGAACCTAAAGACTTGCTCCAGGCCGCACAGACGTGTCGA<br>TACTGGCGAATACTTGCGGAAGACAATCTCCTGTGGCGGGAAAAGTGCAAGGAGGAGGGTATTGAT<br>GAACCGCTCCACATTAAACGAAGGAAGGTCATTAAACCCGGCTTCATTCACTCACCATGGAAGAGT<br>GCATACATCCGACAGCATAGGATAGATACTAACTGGCGGAGAGGGGAACTGAAAAGCCCCAGCGGG<br>TCAGGTAGTGGCtccggaGATCAAGTCCAACTGGTGGAGTCTGGTGGCGCTTTGGTGCAGCCAGGT<br>GGCTCTCTGCGTTTGTCCTGTGCCGCTTCTGGCTTCCCAGTGAACCGCTATTCCATGCGCTGGTAT<br>CGCCAGGCTCCAGGCAAAGAGCGTGAGTGGGTAGCCGGTATGTCCAGCGCGGGTGATCGTAGCTCC<br>TATGAAGACTCCGTGAAGGGCCGTTTCACCATCAGCCGTGACGATGCCCGTAACACGGTGTATCTG<br>CAAATGAACAGCTTGAAACCTGAAGATACGGCCGTGTATTACTGTAATGTGAACGTGGGCTTCGAG<br>TATTGGGGCCAAGGCACCCAGGTCACCGTCTCCAGCggatccTGA |
| FLAG-SKP2$_{2-147}$-vhhGFP4 | ATGggaggccttGATTACAAGGACGACGACGACAAGgctagcCACCGAAAGCACTTGCAGGAGATA<br>CCTGATCTTTCTAGCAATGTAGCAACTTCTTTTACGTGGGGGTGGATAGTAGTAAGACATCTGAA<br>CTGTTGTCCGGTATGGGGGTATCAGCTTTGGAGAAAGAAGAGCCTGACAGTGAGAACATACCGCAG<br>GAACTCCTGTCTAATCTCGGACATCCCGAATCCCCACCTAGGAAGAGGTTGAAGTCAAAAGGAAGT<br>GATAAAGATTTCGTCATTGTGCGAAGACCAAAACTGAACCGAGAAAATTTTCCTGGAGTTTCTTGG<br>GACTCCTTGCCTGACGAACTCCTTCTGGGTATATTTTCATGTTTGTGCCTGCCGGAACTGTTGAAA<br>GTTTCCGGCGTCTGTAAACGGTGGTATCGCCTCGCCTCCGATGAAAGCCTGTGGCAGACATTGGAC<br>CTGACAGGCAAGAACTTGAGCGGGTCAGGTAGTGGCtccggaGATCAAGTCCAACTGGTGGAGTCT<br>GGTGGCGCTTTGGTGCAGCCAGGTGGCTCTCTGCGTTTGTCCTGTGCCGCTTCTGGCTTCCCAGTG<br>AACCGCTATTCCATGCGCTGGTATCGCCAGGCTCCAGGCAAAGAGCGTGAGTGGGTAGCCGGTATG<br>TCCAGCGCGGGTGATCGTAGCTCCTATGAAGACTCCGTGAAGGGCCGTTTCACCATCAGCCGTGAC<br>GATGCCCGTAACACGGTGTATCTGCAAATGAACAGCTTGAAACCTGAAGATACGGCCGTGTATTAC<br>TGTAATGTGAACGTGGGCTTCGAGTATTGGGGCCAAGGCACCCAGGTCACCGTCTCCAGCggatcc<br>TGA |

| | |
|---|---|
| FLAG-vhhGFP4-VHL$_{152-213}$ | ATGggaggccttGATTACAAGGACGACGACGACAAGgctagcGATCAAGTCCAACTGGTGGAGTCT<br>GGTGGCGCTTTGGTGCAGCCAGGTGGCTCTCTGCGTTTGTCCTGTGCCGCTTCTGGCTTCCCAGTG<br>AACCGCTATTCCATGCGCTGGTATCGCCAGGCTCCAGGCAAAGAGCGTGAGTGGGTAGCCGGTATG<br>TCCAGCGCGGGTGATCGTAGCTCCTATGAAGACTCCGTGAAGGGCCGTTTCACCATCAGCCGTGAC<br>GATGCCCGTAACACGGTGTATCTGCAAATGAACAGCTTGAAACCTGAAGATACGGCCGTGTATTAC<br>TGTAATGTGAACGTGGGCTTCGAGTATTGGGGCCAAGGCACCCAGGTCACCGTCTCCAGCGGCAGT<br>GGTAGTGGCtccggaACTTTGCCGGTTTACACCCTGAAGGAGAGATGTCTCCAAGTTGTTCGCAGT<br>CTGGTCAAGCCTGAGAATTATCGACGCCTCGATATTGTAAGGTCTTTGTACGAAGATTTGGAAGAC<br>CATCCGAATGTTCAGAAGGACCTGGAGAGGCTTACACAGGAGAGAATCGCACATCAACGAATGGGT<br>GACGGATCCTGA |
| FLAG-CRBN$_{2-320}$-vhhGFP4 | ATGggaggccttGATTACAAGGACGACGACGACAAGgctagcGCGGGAGAGGGGGACCAACAGGAT<br>GCCGCACACAACATGGGAAACCACCTTCCATTGCTTCCTGCCGAATCTGAGGAGGAGGACGAGATG<br>GAAGTGGAAGACCAGGACTCTAAAGAGGCAAAAAAGCCGAATATAATAAACTTTGACACCTCTCTT<br>CCGACCTCCCATACTTATCTCGGCGCAGACATGGAGGAATTTCACGGGAGAACATTGCACGACGAC<br>GACTCCTGCCAAGTTATTCCGGTTCTGCCTCAGGTAATGATGATATTGATTCCGGGACAGACGTTG<br>CCACTTCAACTTTTCCATCCACAAGAAGTGTCAATGGTGCGAAACTTGATACAAAAGGACAGAACG<br>TTTGCCGTTCTTGCGTACAGTAATGTACAAGAGCGGGAAGCTCAGTTTGGCACCACGGCCGAAATC<br>TACGCATATAGAGAGGAACAAGATTTCGGTATTGAAATCGTTAAGGTTAAAGCCATAGGTAGGCAA<br>CGCTTTAAGGTGCTCGAGTTGAGAACCCAGAGCGACGGCATCCAGCAAGCCAAGGTACAGATTCTG<br>CCCGAATGCGTATTGCCTAGCACTATGAGCGCGGTGCAACTGGAAAGCCTCAATAAGTGCCAAATT<br>TTTCCATCTAAGCCAGTCAGCCGGGAAGACCAGTGTTCTTACAAATGGTGGCAGAAATACCAAAAG<br>CGGAAATTCCACTGTGCCAACCTGACGTCTTGGCCAAGGTGGCTCTACAGTCTTTATGATGCCGAA<br>ACGCTGATGGATCGGATTAAAAAGCAGCTCCGGGAGTGGGATGAAAACCTCAAAGATGATAGCCTT<br>CCCAGTAATCCGATTGACTTCAGTTATAGGGTAGCAGCCTGCCTCCCCATTGACGACGTACTCAGA<br>ATCCAACTTTTGAAGATTGGATCAGCCATTCAAAGACTGCGCTGTGAACTCGATATAATGAATAAG<br>TGTACATCTAGCGGGTCAGGTAGTGGCtccggaGATCAAGTCCAACTGGTGGAGTCTGGTGGCGCT<br>TTGGTGCAGCCAGGTGGCTCTCTGCGTTTGTCCTGTGCCGCTTCTGGCTTCCCAGTGAACCGCTAT<br>TCCATGCGCTGGTATCGCCAGGCTCCAGGCAAAGAGCGTGAGTGGGTAGCCGGTATGTCCAGCGCG<br>GGTGATCGTAGCTCCTATGAAGACTCCGTGAAGGGCCGTTTCACCATCAGCCGTGACGATGCCCGT<br>AACACGGTGTATCTGCAAATGAACAGCTTGAAACCTGAAGATACGGCCGTGTATTACTGTAATGTG<br>AACGTGGGCTTCGAGTATTGGGGCCAAGGCACCCAGGTCACCGTCTCCAGCggatccTGA |
| FLAG-DDB2$_{2-114}$-vhhGFP4 | ATGggaggccttGATTACAAGGACGACGACGACAAGgctagcGCTCCCAAAAAGAGGCCCGAGACC<br>CAGAAAACTTCAGAGATCGTTCTTCGCCCGAGGAATAAGAGGAGCCGAAGCCCACTCGAACTGGAA<br>CCCGAGGCTAAGAAGCTGTGTGCTAAAGGTAGTGGCCCTAGTAGACGGTGTGATAGCGATTGTCTG<br>TGGGTTGGCCTGGCGGGGCCTCAAATATTGCCGCCTTGTCGGTCTATCGTGCGGACCCTGCACCAG<br>CATAAGCTGGGCCGCGCAAGTTGGCCATCCGTGCAGCAGGGTCTTCAACAGAGCTTCCTTCACACA<br>CTGGATTCCTATAGAATTTTGCAAAAAGCAGCGCCTTTTGACAGACGCGCGAGCGGGTCAGGTAGT<br>GGCtccggaGATCAAGTCCAACTGGTGGAGTCTGGTGGCGCTTTGGTGCAGCCAGGTGGCTCTCTG<br>CGTTTGTCCTGTGCCGCTTCTGGCTTCCCAGTGAACCGCTATTCCATGCGCTGGTATCGCCAGGCT<br>CCAGGCAAAGAGCGTGAGTGGGTAGCCGGTATGTCCAGCGCGGGTGATCGTAGCTCCTATGAAGAC<br>TCCGTGAAGGGCCGTTTCACCATCAGCCGTGACGATGCCCGTAACACGGTGTATCTGCAAATGAAC<br>AGCTTGAAACCTGAAGATACGGCCGTGTATTACTGTAATGTGAACGTGGGCTTCGAGTATTGGGGC<br>CAAGGCACCCAGGTCACCGTCTCCAGCggatccTGA |
| FLAG-vhhGFP4-SOCS2$_{143-198}$ | ATGggaggccttGATTACAAGGACGACGACGACAAGgctagcGATCAAGTCCAACTGGTGGAGTCT<br>GGTGGCGCTTTGGTGCAGCCAGGTGGCTCTCTGCGTTTGTCCTGTGCCGCTTCTGGCTTCCCAGTG<br>AACCGCTATTCCATGCGCTGGTATCGCCAGGCTCCAGGCAAAGAGCGTGAGTGGGTAGCCGGTATG<br>TCCAGCGCGGGTGATCGTAGCTCCTATGAAGACTCCGTGAAGGGCCGTTTCACCATCAGCCGTGAC<br>GATGCCCGTAACACGGTGTATCTGCAAATGAACAGCTTGAAACCTGAAGATACGGCCGTGTATTAC<br>TGTAATGTGAACGTGGGCTTCGAGTATTGGGGCCAAGGCACCCAGGTCACCGTCTCCAGCGGCAGT<br>GGTAGTGGCtccggaCCCAGGAATGGGACTGTCCACTTGTATCTTACAAAACCGCTCTACACCTCA |

| | |
|---|---|
| | GCACCTTCATTGCAACACTTGTGTCGCCTGACCATTAACAAATGTACTGGGGCTATATGGGGCCTG<br>CCACTTCCGACACGCTTGAAAGATTATCTGGAAGAGTACAAGTTTCAGGTGGGATCCTGA |
| FLAG-vhhGFP4-ASB1$_{266-335}$ | ATGggaggccttGATTACAAGGACGACGACGACAAGgctagcGATCAAGTCCAACTGGTGGAGTCT<br>GGTGGCGCTTTGGTGCAGCCAGGTGGCTCTCTGCGTTTGTCCTGTGCCGCTTCTGGCTTCCCAGTG<br>AACCGCTATTCCATGCGCTGGTATCGCCAGGCTCCAGGCAAAGAGCGTGAGTGGGTAGCCGGTATG<br>TCCAGCGCGGGTGATCGTAGCTCCTATGAAGACTCCGTGAAGGGCCGTTTCACCATCAGCCGTGAC<br>GATGCCCGTAACACGGTGTATCTGCAAATGAACAGCTTGAAACCTGAAGATACGGCCGTGTATTAC<br>TGTAATGTGAACGTGGGCTTCGAGTATTGGGGCCAAGGCACCCAGGTCACCGTCTCCAGCGGCAGT<br>GGTAGTGGCtccggaGTTAAGTGGGAGAGCCTTGGACCCGAATCTAGGGGTCGCCGGAAAGTTGAC<br>CCGGAGGCGCTGCAAGTCTTCAAAGAAGCTAGAAGCGTTCCGCGAACCCTCCTCTGTCTTTGCCGA<br>GTGGCGGTCAGACGGGCGCTGGGAAAGCACAGGTTGCATCTTATTCCGTCCCTGCCTCTTCCAGAT<br>CCCATCAAGAAATTCTTGCTGCACGAAGGATCCTGA |
| FLAG-vhhGFP4-CHIP$_{128-303}$ | ATGggaggccttGATTACAAGGACGACGACGACAAGgctagcGATCAAGTCCAACTGGTGGAGTCT<br>GGTGGCGCTTTGGTGCAGCCAGGTGGCTCTCTGCGTTTGTCCTGTGCCGCTTCTGGCTTCCCAGTG<br>AACCGCTATTCCATGCGCTGGTATCGCCAGGCTCCAGGCAAAGAGCGTGAGTGGGTAGCCGGTATG<br>TCCAGCGCGGGTGATCGTAGCTCCTATGAAGACTCCGTGAAGGGCCGTTTCACCATCAGCCGTGAC<br>GATGCCCGTAACACGGTGTATCTGCAAATGAACAGCTTGAAACCTGAAGATACGGCCGTGTATTAC<br>TGTAATGTGAACGTGGGCTTCGAGTATTGGGGCCAAGGCACCCAGGTCACCGTCTCCAGCGGCAGT<br>GGTAGTGGCtccggaCGCCTGAATTTTGGAGACGATATTCCTTCTGCGCTCAGAATCGCTAAAAAG<br>AAGAGATGGAATAGTATTGAAGAGAGGCGCATCCACCAAGAGAGTGAGCTTCACTCCTACTTGAGT<br>AGGCTGATTGCTGCCGAAAGAGAACGAGAACTTGAGGAGTGCCAACGAAATCATGAGGGCGACGAA<br>GACGACTCACACGTAAGAGCCCAACAGGCTTGCATCGAAGCCAAGCACGACAAATACATGGCGGAC<br>ATGGACGAACTTTTTAGTCAAGTTGACGAAAAACGCAAAAAGCGCGATATACCGGATTACCTTTGC<br>GGTAAGATTTCTTTCGAGCTGATGCGCGAACCGTGCATTACACCTAGCGGGATCACGTACGACCGC<br>AAAGACATTGAGGAACACTTGCAACGGGTTGGGCACTTCGATCCTGTGACACGGTCACCGTTGACT<br>CAAGAACAACTCATACCGAACTTGGCAATGAAGGAGGTCATCGATGCTTTCATTTCTGAGAATGGC<br>TGGGTCGAGGACTATGGATCCTGA |
| FLAG-NS1$_{v1}$-SPOP$_{167-374}$ | ATGGATTACAAGGACGACGACGACAAGgctagcGGCAGTGTAAGTAGTGTGCCTACGAAACTTGAG<br>GTCGTGGCGGCTACTCCTACGAGCCTGTTGATTTCCTGGGATGCCCCAGCCGTGACAGTGGACTAC<br>TATGTCATTACATATGGCGAAACCGGCGGGAATAGTCCTGTTCAGAAATTCGAGGTCCCAGGTAGC<br>AAATCTACCGCCACGATTAGCGGGTTGAAACCAGGCGTGGATTATACCATTACGGTCTATGCTTGG<br>GGATGGCATGGGCAAGTCTATTACTATATGGGATCACCAATCTCCATTAACTACCGCACAGGTAGC<br>GGCAGCGGTtccggaAGCGTGAACATCTCCGGCCAGAACACAATGAACATGGTCAAGGTGCCCGAG<br>TGCAGACTGGCCGACGAGCTGGGAGGACTGTGGGAGAACTCCAGGTTTACCGACTGCTGCCTGTGC<br>GTGGCCGGCCAAGAGTTCCAAGCCCACAAAGCCATCCTGGCCGCTAGGTCCCCCGTGTTCAGCGCC<br>ATGTTCGAGCACGAGATGGAGGAGTCCAAGAAGAACAGAGTGGAGATTAACGATGTGGAGCCCGAG<br>GTGTTCAAAGAAATGATGTGCTTCATCTACACCGGCAAGGCCCCCAACCTGGATAAAATGGCCGAT<br>GACCTGCTGGCCGCCGCCGATAAGTACGCCCTGGAGAGACTGAAGGTGATGTGCGAGGACGCTCTG<br>TGTTCCAACCTGTCCGTGGAAAATGCCGCCGAGATCCTCATCCTGGCCGACCTGCATAGCGCCGAC<br>CAGCTGAAAACCCAGGCCGTGGACTTCATCAACTATCACGCTTCCGACGTGCTGGAGACCAGCGGA<br>TGGAAGAGCATGGTGGTGAGCCATCCCCATCTCGTGGCCGAAGCCTACAGGAGCCTGGCaAGCGCC<br>CAGTGTCCCTTTCTGGGCCCTCCCAGGAAGAGACTGAAACAGAGCTGA |
| FLAG-NS1$_{v2}$-SPOP$_{167-374}$ | ATGggaggccttGATTACAAGGACGACGACGACAAGgctagcGGCAGTGTAAGTAGTGTGCCTACG<br>AAACTTGAGGTCGTGGCGGCTACTCCTACGAGCCTGTTGATTTCCTGGGATGCCCCAGCCGTGACA<br>GTGGACTACTATGTCATTACATATGGCGAAACCGGCGGGAATAGTCCTGTTCAGAAATTCGAGGTC<br>CCAGGTAGCAAATCTACCGCCACGATTAGCGGGTTGAAACCAGGCGTGGATTATACCATTACGGTC<br>TATGCTTGGGGATGGCATGGGCAAGTCTATTACTATATGGGATCACCAATCTCCATTAACTACCGC<br>ACAGGTAGCGGCAGCGGTtccggaAGCGTGAACATCTCCGGCCAGAACACAATGAACATGGTCAAG<br>GTGCCCGAGTGCAGACTGGCCGACGAGCTGGGAGGACTGTGGGAGAACTCCAGGTTTACCGACTGC<br>TGCCTGTGCGTGGCCGGCCAAGAGTTCCAAGCCCACAAAGCCATCCTGGCCGCTAGGTCCCCCGTG<br>TTCAGCGCCATGTTCGAGCACGAGATGGAGGAGTCCAAGAAGAACAGAGTGGAGATTAACGATGTG |

| | |
|---|---|
| | GAGCCCGAGGTGTTCAAAGAAATGATGTGCTTCATCTACACCGGCAAGGCCCCCAACCTGGATAAA<br>ATGGCCGATGACCTGCTGGCCGCCGCCGATAAGTACGCCCTGGAGAGACTGAAGGTGATGTGCGAG<br>GACGCTCTGTGTTCCAACCTGTCCGTGGAAAATGCCGCCGAGATCCTCATCCTGGCCGACCTGCAT<br>AGCGCCGACCAGCTGAAAACCCAGGCCGTGGACTTCATCAACTATCACGCTTCCGACGTGCTGGAG<br>ACCAGCGGATGGAAGAGCATGGTGGTGAGCCATCCCCATCTCGTGGCCGAAGCCTACAGGAGCCTG<br>GCaAGCGCCCAGTGTCCCTTTCTGGGCCCTCCCAGGAAGAGACTGAAACAGAGCGGATCCTGA |
| FLAG-NS1-SPOP$_{mut}$ | ATGGATTACAAGGACGACGACGACAAGgctagcGGCAGTGTAAGTAGTGTGCCTACGAAACTTGAG<br>GTCGTGGCGGCTACTCCTACGAGCCTGTTGATTTCCTGGGATGCCCCAGCCGTGACAGTGGACTAC<br>TATGTCATTACATATGGCGAAACCGGCGGGAATAGTCCTGTTCAGAAATTCGAGGTCCCAGGTAGC<br>AAATCTACCGCCACGATTAGCGGGTTGAAACCAGGCGTGGATTATACCATTACGGTCTATGCTTGG<br>GGATGGCATGGGCAAGTCTATTACTATATGGGATCACCAATCTCCATTAACTACCGCACAGGTAGC<br>GGCAGCGGTtccggaAGCGTGAACATCTCCGGCCAGAACACAATGAACATGGTCAAGGTGCCCGAG<br>TGCAGACTGGCCGACGAGCTGGGAGGACTGTGGGAGAACTCCAGGTTTACCGACTGCTGCCTGTGC<br>GTGGCCGGCCAAGAGTTCCAAGCCCACAAAGCCATCCTGGCCGCTAGGTCCCCCGTGTTCAGCGCC<br>ATGTTCGAGCACGAGATGGAGGAGTCCAAGAAGAACAGAGTGGAGATTAACGATGTGGAGCCCGAG<br>GTGTTCAAAGAAATGATGTGCTTCATCTACACCGGCAAGGCCCCCAACCTGGATAAAATGGCCGAT<br>GACCTGCTGGCCGCCGCCGATAAGTACGCCCTGGAGAGACTGAAGGTGATGTGCGAGGACGCTCTG<br>TGTTCCAACCTGTCCGTGGAAAATTATCACGCTTCCGACGTGCTGGAGACCAGCGGATGGAAGAGC<br>ATGGTGGTGAGCCATCCCCATCTCGTGGCCGAAGCCTACAGGAGCCTGGCaAGCGCCCAGTGTCCC<br>TTTCTGGGCCCTCCCAGGAAGAGACTGAAACAGAGCTGA |
| FLAG-FN3-SPOP$_{167-374}$ | ATGGATTACAAGGACGACGACGACAAGgctagcGTGTCCGATGTGCCTAGGGACCTGGAAGTCGTT<br>GCTGCAACGCCCACTAGCCTGCTTATTTCCTGGGACGCTCCCGCCGTTACAGTCAGGTACTATAGG<br>ATAACTTATGGTGAAACCGGGGGTAACTCACCAGTCCAGGAGTTCACTGTACCCGGTTCAAAATCA<br>ACGGCGACAATCTCCGGGTTGAAACCTGGTGTAGACTATACCATCACGGTCTATGCCGTTACAGGA<br>CGCGGTGATTCACCCGCGTCTTCCAAACCCATTAGTATAAATTACCGCACGGGTAGCGGCAGCGGT<br>tccggaAGCGTGAACATCTCCGGCCAGAACACAATGAACATGGTCAAGGTGCCCGAGTGCAGACTG<br>GCCGACGAGCTGGGAGGACTGTGGGAGAACTCCAGGTTTACCGACTGCTGCCTGTGCGTGGCCGGC<br>CAAGAGTTCCAAGCCCACAAAGCCATCCTGGCCGCTAGGTCCCCCGTGTTCAGCGCCATGTTCGAG<br>CACGAGATGGAGGAGTCCAAGAAGAACAGAGTGGAGATTAACGATGTGGAGCCCGAGGTGTTCAAA<br>GAAATGATGTGCTTCATCTACACCGGCAAGGCCCCCAACCTGGATAAAATGGCCGATGACCTGCTG<br>GCCGCCGCCGATAAGTACGCCCTGGAGAGACTGAAGGTGATGTGCGAGGACGCTCTGTGTTCCAAC<br>CTGTCCGTGGAAAATGCCGCCGAGATCCTCATCCTGGCCGACCTGCATAGCGCCGACCAGCTGAAA<br>ACCCAGGCCGTGGACTTCATCAACTATCACGCTTCCGACGTGCTGGAGACCAGCGGATGGAAGAGC<br>ATGGTGGTGAGCCATCCCCATCTCGTGGCCGAAGCCTACAGGAGCCTGGCaAGCGCCCAGTGTCCC<br>TTTCTGGGCCCTCCCAGGAAGAGACTGAAACAGAGCTGA |
| FLAG-K27-SPOP$_{167-374}$ | ATGGATTACAAGGACGACGACGACAAGgctagcGATCTCGGAAAAAAATTGCTTGAGGCCGCTAGA<br>GCAGGCCAGGATGACGAAGTGCGGATTCTGATGGCAAACGGCGCTGACGTTAATGCGCATGATACC<br>TTTGGATTTACCCCACTCCATCTGGCGGCTCTTTACGGGCATTTGGAGATAGTAGAAGTGCTGCTG<br>AAAAATGGTGCAGACGTCAACGCAGACGATTCCTATGGGAGAACGCCCCTTCATTTGGCTGCTATG<br>CGAGGACATTTGGAGATCGTAGAAGTATTGCTGAAGTATGGCGCTGATGTCAACGCGGCTGACGAG<br>GAAGGCAGAACTCCTTTGCACCTGGCTGCGAAACGGGGTCACCTGGAAATAGTGGAAGTCCTTCTT<br>AAAAACGGCGCAGATGTGAACGCTCAGGACAAGTTTGGGAAGACAGCGTTTGACATAAGTATCGAC<br>AATGGAAACGAAGATTTGGCTGAAATTTTGCAGAAATTGGGCAGTGGTAGTGGCtccggaAGCGTG<br>AACATCTCCGGCCAGAACACAATGAACATGGTCAAGGTGCCCGAGTGCAGACTGGCCGACGAGCTG<br>GGAGGACTGTGGGAGAACTCCAGGTTTACCGACTGCTGCCTGTGCGTGGCCGGCCAAGAGTTCCAA<br>GCCCACAAAGCCATCCTGGCCGCTAGGTCCCCCGTGTTCAGCGCCATGTTCGAGCACGAGATGGAG<br>GAGTCCAAGAAGAACAGAGTGGAGATTAACGATGTGGAGCCCGAGGTGTTCAAAGAAATGATGTGC<br>TTCATCTACACCGGCAAGGCCCCCAACCTGGATAAAATGGCCGATGACCTGCTGGCCGCCGCCGAT<br>AAGTACGCCCTGGAGAGACTGAAGGTGATGTGCGAGGACGCTCTGTGTTCCAACCTGTCCGTGGAA<br>AATGCCGCCGAGATCCTCATCCTGGCCGACCTGCATAGCGCCGACCAGCTGAAAACCCAGGCCGTG<br>GACTTCATCAACTATCACGCTTCCGACGTGCTGGAGACCAGCGGATGGAAGAGCATGGTGGTGAGC |

| | |
|---|---|
| | CATCCCCATCTCGTGGCCGAAGCCTACAGGAGCCTGGCaAGCGCCCAGTGTCCCTTTCTGGGCCCT<br>CCCAGGAAGAGACTGAAACAGAGCTGA |
| FLAG-K27-SPOP$_{mut}$ | ATGGATTACAAGGACGACGACGACAAGgctagcGATCTCGGAAAAAAATTGCTTGAGGCCGCTAGA<br>GCAGGCCAGGATGACGAAGTGCGGATTCTGATGGCAAACGGCGCTGACGTTAATGCGCATGATACC<br>TTTGGATTTACCCCACTCCATCTGGCGGCTCTTTACGGGCATTTGGAGATAGTAGAAGTGCTGCTG<br>AAAAATGGTGCAGACGTCAACGCAGACGATTCCTATGGGAGAACGCCCCTTCATTTGGCTGCTATG<br>CGAGGACATTTGGAGATCGTAGAAGTATTGCTGAAGTATGGCGCTGATGTCAACGCGGCTGACGAG<br>GAAGGCAGAACTCCTTTGCACCTGGCTGCGAAACGGGGTCACCTGGAAATAGTGGAAGTCCTTCTT<br>AAAAACGGCGCAGATGTGAACGCTCAGGACAAGTTTGGGAAGACAGCGTTTGACATAAGTATCGAC<br>AATGGAAACGAAGATTTGGCTGAAATTTTGCAGAAATTGGGCAGTGGTAGTGGCtccggaAGCGTG<br>AACATCTCCGGCCAGAACACAATGAACATGGTCAAGGTGCCCGAGTGCAGACTGGCCGACGAGCTG<br>GGAGGACTGTGGGAGAACTCCAGGTTTACCGACTGCTGCCTGTGCGTGGCCGGCCAAGAGTTCCAA<br>GCCCACAAAGCCATCCTGGCCGCTAGGTCCCCCGTGTTCAGCGCCATGTTCGAGCACGAGATGGAG<br>GAGTCCAAGAAGAACAGAGTGGAGATTAACGATGTGGAGCCCGAGGTGTTCAAAGAAATGATGTGC<br>TTCATCTACACCGGCAAGGCCCCCAACCTGGATAAAATGGCCGATGACCTGCTGGCCGCCGCCGAT<br>AAGTACGCCCTGGAGAGACTGAAGGTGATGTGCGAGGACGCTCTGTGTTCCAACCTGTCCGTGGAA<br>AATTATCACGCTTCCGACGTGCTGGAGACCAGCGGATGGAAGAGCATGGTGGTGAGCCATCCCCAT<br>CTCGTGGCCGAAGCCTACAGGAGCCTGGCaAGCGCCCAGTGTCCCTTTCTGGGCCCTCCCAGGAAG<br>AGACTGAAACAGAGCTGA |
| FLAG-K27$_{mut}$-SPOP$_{167\text{-}374}$ | ATGGATTACAAGGACGACGACGACAAGgctagcGATCTCGGAAAAAAATTGCTTGAGGCCGCTAGA<br>GCAGGCCAGGATGACGAAGTGCGGATTCTGATGGCAAACGGCGCTGACGTTAATGCGCATGATACC<br>TTTGGATTTACCCCACTCCATCTGGCGGCTCTTTACGGGCATTTGGAGATAGTAGAAGTGCTGCTG<br>AAAAATGGTGCAGACGTCAACGCAGACGATTCCTATGGGgccACGCCCCTTCATTTGGCTGCTATG<br>CGAGGACATTTGGAGATCGTAGAAGTATTGCTGAAGTATGGCGCTGATGTCAACGCGGCTGACGAG<br>GAAGGCgccACTCCTTTGCACCTGGCTGCGAAAgccGGTCACCTGGAAATAGTGGAAGTCCTTCTT<br>AAAAACGGCGCAGATGTGAACGCTCAGGACAAGTTTGGGAAGACAGCGTTTGACATAAGTATCGAC<br>AATGGAAACGAAGATTTGGCTGAAATTTTGCAGAAATTGGGCAGTGGTAGTGGCtccggaAGCGTG<br>AACATCTCCGGCCAGAACACAATGAACATGGTCAAGGTGCCCGAGTGCAGACTGGCCGACGAGCTG<br>GGAGGACTGTGGGAGAACTCCAGGTTTACCGACTGCTGCCTGTGCGTGGCCGGCCAAGAGTTCCAA<br>GCCCACAAAGCCATCCTGGCCGCTAGGTCCCCCGTGTTCAGCGCCATGTTCGAGCACGAGATGGAG<br>GAGTCCAAGAAGAACAGAGTGGAGATTAACGATGTGGAGCCCGAGGTGTTCAAAGAAATGATGTGC<br>TTCATCTACACCGGCAAGGCCCCCAACCTGGATAAAATGGCCGATGACCTGCTGGCCGCCGCCGAT<br>AAGTACGCCCTGGAGAGACTGAAGGTGATGTGCGAGGACGCTCTGTGTTCCAACCTGTCCGTGGAA<br>AATGCCGCCGAGATCCTCATCCTGGCCGACCTGCATAGCGCCGACCAGCTGAAAACCCAGGCCGTG<br>GACTTCATCAACTATCACGCTTCCGACGTGCTGGAGACCAGCGGATGGAAGAGCATGGTGGTGAGC<br>CATCCCCATCTCGTGGCCGAAGCCTACAGGAGCCTGGCaAGCGCCCAGTGTCCCTTTCTGGGCCCT<br>CCCAGGAAGAGACTGAAACAGAGCTGA |
| FLAG-K55-SPOP$_{167\text{-}374}$ | ATGggaggccttGATTACAAGGACGACGACGACAAGgctagcGACCTCGGTAAGAAACTGTTGGAG<br>GCTGCGAGAGCAGGCCAAGATGACGAAGTAAGAATCCTTATGGCAAATGGTGCGGATGTCAACGCC<br>AACGATTCCGCAGGACACACTCCACTCCACCTTGCTGCTAAGCGAGGTCACCTTGAAATAGTCGAA<br>GTGCTGCTCAAACATGGAGCTGATGTCAACGCAATGGATAACACAGGGTTCACACCTCTCCATCTT<br>GCTGCTTTGAGGGGGCACCTCGAAATTGTTGAAGTTCTCCTGAAGAACGGGGCCGACGTTAATGCA<br>CAAGATCGCACAGGACGAACTCCACTCCATCTGGCTGCGAAACTGGGACATCTGGAGATTGTAGAG<br>GTACTTCTGAAGAACGGTGCAGACGTCAATGCGCAAGATAAATTTGGCAAAACTGCCTTTGACATT<br>TCTATCGACAACGGCAACGAGGATCTCGCTGAAATTCTGCAAAAGCTGGGCAGTGGTAGTGGCtcc<br>ggaAGCGTGAACATCTCCGGCCAGAACACAATGAACATGGTCAAGGTGCCCGAGTGCAGACTGGCC<br>GACGAGCTGGGAGGACTGTGGGAGAACTCCAGGTTTACCGACTGCTGCCTGTGCGTGGCCGGCCAA<br>GAGTTCCAAGCCCACAAAGCCATCCTGGCCGCTAGGTCCCCCGTGTTCAGCGCCATGTTCGAGCAC<br>GAGATGGAGGAGTCCAAGAAGAACAGAGTGGAGATTAACGATGTGGAGCCCGAGGTGTTCAAAGAA<br>ATGATGTGCTTCATCTACACCGGCAAGGCCCCCAACCTGGATAAAATGGCCGATGACCTGCTGGCC<br>GCCGCCGATAAGTACGCCCTGGAGAGACTGAAGGTGATGTGCGAGGACGCTCTGTGTTCCAACCTG |

| | |
|---|---|
| | TCCGTGGAAAATGCCGCCGAGATCCTCATCCTGGCCGACCTGCATAGCGCCGACCAGCTGAAAACC<br>CAGGCCGTGGACTTCATCAACTATCACGCTTCCGACGTGCTGGAGACCAGCGGATGGAAGAGCATG<br>GTGGTGAGCCATCCCCATCTCGTGGCCGAAGCCTACAGGAGCCTGGCaAGCGCCCAGTGTCCCTTT<br>CTGGGCCCTCCCAGGAAGAGACTGAAACAGAGCGGATCCTGA |
| FLAG-R11.1.6-SPOP$_{167-374}$ | ATGGATTACAAGGACGACGACGACAAGgctagcGCTACCGTAAAGTTCACTCATCAGGGAGAAGAA<br>AAGCAAGTTGATATATCTAAGATAAAATGGGTAATAAGATGGGGACAATATATCTGGTTCAAGTAC<br>GACGAAGATGGAGGTGCGAAGGGGTGGGGCTATGTGTCCGAAAAAGACGCTCCTAAAGAACTTTTG<br>CAGATGCTTAAAAAACGCGGCAGTGGTAGTGGCtccggaAGCGTGAACATCTCCGGCCAGAACACA<br>ATGAACATGGTCAAGGTGCCCGAGTGCAGACTGGCCGACGAGCTGGGAGGACTGTGGGAGAACTCC<br>AGGTTTACCGACTGCTGCCTGTGCGTGGCCGGCCAAGAGTTCCAAGCCCACAAAGCCATCCTGGCC<br>GCTAGGTCCCCCGTGTTCAGCGCCATGTTCGAGCACGAGATGGAGGAGTCCAAGAAGAACAGAGTG<br>GAGATTAACGATGTGGAGCCCGAGGTGTTCAAAGAAATGATGTGCTTCATCTACACCGGCAAGGCC<br>CCCAACCTGGATAAAATGGCCGATGACCTGCTGGCCGCCGCCGATAAGTACGCCCTGGAGAGACTG<br>AAGGTGATGTGCGAGGACGCTCTGTGTTCCAACCTGTCCGTGGAAAATGCCGCCGAGATCCTCATC<br>CTGGCCGACCTGCATAGCGCCGACCAGCTGAAAACCCAGGCCGTGGACTTCATCAACTATCACGCT<br>TCCGACGTGCTGGAGACCAGCGGATGGAAGAGCATGGTGGTGAGCCATCCCCATCTCGTGGCCGAA<br>GCCTACAGGAGCCTGGCaAGCGCCCAGTGTCCCTTTCTGGGCCCTCCCAGGAAGAGACTGAAACAG<br>AGCTGA |
| FLAG-RBD-CRD-SPOP$_{167-374}$ | ATGGATTACAAGGACGACGACGACAAGgctagcCCTTCCAAGACAAGCAACACAATTCGGGTCTTT<br>CTGCCGAACAAACAAAGGACTGTCGTCAATGTGAGGAATGGAATGTCTCTGCACGACTGTCTGATG<br>AAGGCGCTGAAGGTCAGGGGGCTCCAGCCCGAATGCTGCGCAGTGTTTAGGTTGCTTCATGAGCAT<br>AAGGGCAAAAAAGCACGGCTGGACTGGAATACTGACGCTGCCTCACTCATAGGCGAAGAGTTGCAG<br>GTCGATTTCTTGGACCATGTACCGCTTACAACTCATAATTTCGCTCGGAAAACCTTCCTTAAACTG<br>GCTTTTTGCGATATATGTCAAAAATTCCTTCTTAACGGGTTCCGCTGTCAGACTTGCGGATATAAG<br>TTTCATGAACATTGCTCCACAAAGGTGCCAACCATGTGCGTCGATTGGTCTAACATCAGACAGTTG<br>CTTTTGTTCCCCAATTCAACGATTGGGGATTCTGGCGTGCCTGCTCTCCCATCTCTCACTATGAGG<br>AGAATGAGAGAGTCCGGCAGTGGTAGTGGCtccggaAGCGTGAACATCTCCGGCCAGAACACAATG<br>AACATGGTCAAGGTGCCCGAGTGCAGACTGGCCGACGAGCTGGGAGGACTGTGGGAGAACTCCAGG<br>TTTACCGACTGCTGCCTGTGCGTGGCCGGCCAAGAGTTCCAAGCCCACAAAGCCATCCTGGCCGCT<br>AGGTCCCCCGTGTTCAGCGCCATGTTCGAGCACGAGATGGAGGAGTCCAAGAAGAACAGAGTGGAG<br>ATTAACGATGTGGAGCCCGAGGTGTTCAAAGAAATGATGTGCTTCATCTACACCGGCAAGGCCCCC<br>AACCTGGATAAAATGGCCGATGACCTGCTGGCCGCCGCCGATAAGTACGCCCTGGAGAGACTGAAG<br>GTGATGTGCGAGGACGCTCTGTGTTCCAACCTGTCCGTGGAAAATGCCGCCGAGATCCTCATCCTG<br>GCCGACCTGCATAGCGCCGACCAGCTGAAAACCCAGGCCGTGGACTTCATCAACTATCACGCTTCC<br>GACGTGCTGGAGACCAGCGGATGGAAGAGCATGGTGGTGAGCCATCCCCATCTCGTGGCCGAAGCC<br>TACAGGAGCCTGGCaAGCGCCCAGTGTCCCTTTCTGGGCCCTCCCAGGAAGAGACTGAAACAGAGC<br>TGA |
| FLAG-RBD-CRD-SPOP$_{mut}$ | ATGGATTACAAGGACGACGACGACAAGgctagcCCTTCCAAGACAAGCAACACAATTCGGGTCTTT<br>CTGCCGAACAAACAAAGGACTGTCGTCAATGTGAGGAATGGAATGTCTCTGCACGACTGTCTGATG<br>AAGGCGCTGAAGGTCAGGGGGCTCCAGCCCGAATGCTGCGCAGTGTTTAGGTTGCTTCATGAGCAT<br>AAGGGCAAAAAAGCACGGCTGGACTGGAATACTGACGCTGCCTCACTCATAGGCGAAGAGTTGCAG<br>GTCGATTTCTTGGACCATGTACCGCTTACAACTCATAATTTCGCTCGGAAAACCTTCCTTAAACTG<br>GCTTTTTGCGATATATGTCAAAAATTCCTTCTTAACGGGTTCCGCTGTCAGACTTGCGGATATAAG<br>TTTCATGAACATTGCTCCACAAAGGTGCCAACCATGTGCGTCGATTGGTCTAACATCAGACAGTTG<br>CTTTTGTTCCCCAATTCAACGATTGGGGATTCTGGCGTGCCTGCTCTCCCATCTCTCACTATGAGG<br>AGAATGAGAGAGTCCGGCAGTGGTAGTGGCtccggaAGCGTGAACATCTCCGGCCAGAACACAATG<br>AACATGGTCAAGGTGCCCGAGTGCAGACTGGCCGACGAGCTGGGAGGACTGTGGGAGAACTCCAGG<br>TTTACCGACTGCTGCCTGTGCGTGGCCGGCCAAGAGTTCCAAGCCCACAAAGCCATCCTGGCCGCT<br>AGGTCCCCCGTGTTCAGCGCCATGTTCGAGCACGAGATGGAGGAGTCCAAGAAGAACAGAGTGGAG<br>ATTAACGATGTGGAGCCCGAGGTGTTCAAAGAAATGATGTGCTTCATCTACACCGGCAAGGCCCCC<br>AACCTGGATAAAATGGCCGATGACCTGCTGGCCGCCGCCGATAAGTACGCCCTGGAGAGACTGAAG |

| | |
|---|---|
| | GTGATGTGCGAGGACGCTCTGTGTTCCAACCTGTCCGTGGAAAATTATCACGCTTCCGACGTGCTG<br>GAGACCAGCGGATGGAAGAGCATGGTGGTGAGCCATCCCCATCTCGTGGCCGAAGCCTACAGGAGC<br>CTGGCaAGCGCCCAGTGTCCCTTTCTGGGCCCTCCCAGGAAGAGACTGAAACAGAGCTGA |
| FLAG-RBD-SPOP$_{167\text{-}374}$ | ATGGATTACAAGGACGACGACGACAAGgctagcGGTTCCAAAACATCAAATACCATTCGCGTATTC<br>CTGCCTAATAAGCAGCGTACCGTAGTGAATGTACGCAATGGAATGAGTTTGCACGACTGTTTGATG<br>AAGGCTCTTAAAGTTCGTGGTCTTCAACCAGAGTGTTGTGCAGTATTTCGCCTTCTGCATGAACAT<br>AAAGGAAAGAAGGCTCGCTTGGATTGGAACACAGACGCCGCTTCCCTTATTGGCGAAGAGCTTCAG<br>GTAGATTTTTTGGGTAGCGGCAGCGGTtccggaAGCGTGAACATCTCCGGCCAGAACACAATGAAC<br>ATGGTCAAGGTGCCCGAGTGCAGACTGGCCGACGAGCTGGGAGGACTGTGGGAGAACTCCAGGTTT<br>ACCGACTGCTGCCTGTGCGTGGCCGGCCAAGAGTTCCAAGCCCACAAAGCCATCCTGGCCGCTAGG<br>TCCCCCGTGTTCAGCGCCATGTTCGAGCACGAGATGGAGGAGTCCAAGAAGAACAGAGTGGAGATT<br>AACGATGTGGAGCCCGAGGTGTTCAAAGAAATGATGTGCTTCATCTACACCGGCAAGGCCCCCAAC<br>CTGGATAAAATGGCCGATGACCTGCTGGCCGCCGCCGATAAGTACGCCCTGGAGAGACTGAAGGTG<br>ATGTGCGAGGACGCTCTGTGTTCCAACCTGTCCGTGGAAAATGCCGCCGAGATCCTCATCCTGGCC<br>GACCTGCATAGCGCCGACCAGCTGAAAACCCAGGCCGTGGACTTCATCAACTATCACGCTTCCGAC<br>GTGCTGGAGACCAGCGGATGGAAGAGCATGGTGGTGAGCCATCCCCATCTCGTGGCCGAAGCCTAC<br>AGGAGCCTGGCaAGCGCCCAGTGTCCCTTTCTGGGCCCTCCCAGGAAGAGACTGAAACAGAGCTGA |
| FLAG-RBD-SPOP$_{mut}$ | ATGGATTACAAGGACGACGACGACAAGgctagcGGTTCCAAAACATCAAATACCATTCGCGTATTC<br>CTGCCTAATAAGCAGCGTACCGTAGTGAATGTACGCAATGGAATGAGTTTGCACGACTGTTTGATG<br>AAGGCTCTTAAAGTTCGTGGTCTTCAACCAGAGTGTTGTGCAGTATTTCGCCTTCTGCATGAACAT<br>AAAGGAAAGAAGGCTCGCTTGGATTGGAACACAGACGCCGCTTCCCTTATTGGCGAAGAGCTTCAG<br>GTAGATTTTTTGGGTAGCGGCAGCGGTtccggaAGCGTGAACATCTCCGGCCAGAACACAATGAAC<br>ATGGTCAAGGTGCCCGAGTGCAGACTGGCCGACGAGCTGGGAGGACTGTGGGAGAACTCCAGGTTT<br>ACCGACTGCTGCCTGTGCGTGGCCGGCCAAGAGTTCCAAGCCCACAAAGCCATCCTGGCCGCTAGG<br>TCCCCCGTGTTCAGCGCCATGTTCGAGCACGAGATGGAGGAGTCCAAGAAGAACAGAGTGGAGATT<br>AACGATGTGGAGCCCGAGGTGTTCAAAGAAATGATGTGCTTCATCTACACCGGCAAGGCCCCCAAC<br>CTGGATAAAATGGCCGATGACCTGCTGGCCGCCGCCGATAAGTACGCCCTGGAGAGACTGAAGGTG<br>ATGTGCGAGGACGCTCTGTGTTCCAACCTGTCCGTGGAAAATTATCACGCTTCCGACGTGCTGGAG<br>ACCAGCGGATGGAAGAGCATGGTGGTGAGCCATCCCCATCTCGTGGCCGAAGCCTACAGGAGCCTG<br>GCaAGCGCCCAGTGTCCCTTTCTGGGCCCTCCCAGGAAGAGACTGAAACAGAGCTGA |
| FLAG-NS1-VHL$_{152\text{-}213}$ | ATGggaggccttGATTACAAGGACGACGACGACAAGgctagcGGCAGTGTAAGTAGTGTGCCTACG<br>AAACTTGAGGTCGTGGCGGCTACTCCTACGAGCCTGTTGATTTCCTGGGATGCCCCAGCCGTGACA<br>GTGGACTACTATGTCATTACATATGGCGAAACCGGCGGGAATAGTCCTGTTCAGAAATTCGAGGTC<br>CCAGGTAGCAAATCTACCGCCACGATTAGCGGGTTGAAACCAGGCGTGGATTATACCATTACGGTC<br>TATGCTTGGGGATGGCATGGGCAAGTCTATTACTATATGGGATCACCAATCTCCATTAACTACCGC<br>ACAGGTAGCGGCAGCGGTtccggaACTTTGCCGGTTTACACCCTGAAGGAGAGATGTCTCCAAGTT<br>GTTCGCAGTCTGGTCAAGCCTGAGAATTATCGACGCCTCGATATTGTAAGGTCTTTGTACGAAGAT<br>TTGGAAGACCATCCGAATGTTCAGAAGGACCTGGAGAGGCTTACACAGGAGAGAATCGCACATCAA<br>CGAATGGGTGACGGATCCTGA |
| FLAG-K27-VHL$_{152\text{-}213}$ | ATGggaggccttGATTACAAGGACGACGACGACAAGgctagcGATCTCGGAAAAAAATTGCTTGAG<br>GCCGCTAGAGCAGGCCAGGATGACGAAGTGCGGATTCTGATGGCAAACGGCGCTGACGTTAATGCG<br>CATGATACCTTTGGATTTACCCCACTCCATCTGGCGGCTCTTTACGGGCATTTGGAGATAGTAGAA<br>GTGCTGCTGAAAAATGGTGCAGACGTCAACGCAGACGATTCCTATGGGAGAACGCCCCTTCATTTG<br>GCTGCTATGCGAGGACATTTGGAGATCGTAGAAGTATTGCTGAAGTATGGCGCTGATGTCAACGCG<br>GCTGACGAGGAAGGCAGAACTCCTTTGCACCTGGCTGCGAAACGGGGTCACCTGGAAATAGTGGAA<br>GTCCTTCTTAAAAACGGCGCAGATGTGAACGCTCAGGACAAGTTTGGGAAGACAGCGTTTGACATA<br>AGTATCGACAATGGAAACGAAGATTTGGCTGAAATTTTGCAGAAATTGGGCAGTGGTAGTGGCtcc<br>ggaACTTTGCCGGTTTACACCCTGAAGGAGAGATGTCTCCAAGTTGTTCGCAGTCTGGTCAAGCCT<br>GAGAATTATCGACGCCTCGATATTGTAAGGTCTTTGTACGAAGATTTGGAAGACCATCCGAATGTT<br>CAGAAGGACCTGGAGAGGCTTACACAGGAGAGAATCGCACATCAACGAATGGGTGACGGATCCTGA |

| | |
|---|---|
| FLAG-R11.1.6-$VHL_{152-213}$ | ATGggaggccttGATTACAAGGACGACGACGACAAGgctagcGCTACCGTAAAGTTCACTCATCAG<br>GGAGAAGAAAAGCAAGTTGATATATCTAAGATAAAATGGGTAATAAGATGGGGACAATATATCTGG<br>TTCAAGTACGACGAAGATGGAGGTGCGAAGGGGTGGGGCTATGTGTCCGAAAAAGACGCTCCTAAA<br>GAACTTTTGCAGATGCTTAAAAAACGCGGCAGTGGTAGTGGCtccggaACTTTGCCGGTTTACACC<br>CTGAAGGAGAGATGTCTCCAAGTTGTTCGCAGTCTGGTCAAGCCTGAGAATTATCGACGCCTCGAT<br>ATTGTAAGGTCTTTGTACGAAGATTTGGAAGACCATCCGAATGTTCAGAAGGACCTGGAGAGGCTT<br>ACACAGGAGAGAATCGCACATCAACGAATGGGTGACGGATCCTGA |
| FLAG-RBD-CRD-$VHL_{152-213}$ | ATGggaggccttGATTACAAGGACGACGACGACAAGgctagcCCTTCCAAGACAAGCAACACAATT<br>CGGGTCTTTCTGCCGAACAAACAAAGGACTGTCGTCAATGTGAGGAATGGAATGTCTCTGCACGAC<br>TGTCTGATGAAGGCGCTGAAGGTCAGGGGGCTCCAGCCCGAATGCTGCGCAGTGTTTAGGTTGCTT<br>CATGAGCATAAGGGCAAAAAAGCACGGCTGGACTGGAATACTGACGCTGCCTCACTCATAGGCGAA<br>GAGTTGCAGGTCGATTTCTTGGACCATGTACCGCTTACAACTCATAATTTCGCTCGGAAAACCTTC<br>CTTAAACTGGCTTTTTGCGATATATGTCAAAAATTCCTTCTTAACGGGTTCCGCTGTCAGACTTGC<br>GGATATAAGTTTCATGAACATTGCTCCACAAAGGTGCCAACCATGTGCGTCGATTGGTCTAACATC<br>AGACAGTTGCTTTTGTTCCCCAATTCAACGATTGGGGATTCTGGCGTGCCTGCTCTCCCATCTCTC<br>ACTATGAGGAGAATGAGAGAGTCCGGCAGTGGTAGTGGCtccggaACTTTGCCGGTTTACACCCTG<br>AAGGAGAGATGTCTCCAAGTTGTTCGCAGTCTGGTCAAGCCTGAGAATTATCGACGCCTCGATATT<br>GTAAGGTCTTTGTACGAAGATTTGGAAGACCATCCGAATGTTCAGAAGGACCTGGAGAGGCTTACA<br>CAGGAGAGAATCGCACATCAACGAATGGGTGACGGATCCTGA |
| eGFP | ATGGTGAGCAAGGGCGAGGAGCTGTTCACCGGGGTGGTGCCCATCCTGGTCGAGCTGGACGGCGAC<br>GTAAACGGCCACAAGTTCAGCGTGTCCGGCGAGGGCGAGGGCGATGCCACCTACGGCAAGCTGACC<br>CTGAAGTTCATCTGCACCACCGGCAAGCTGCCCGTGCCCTGGCCCACCCTCGTGACCACCCTGACC<br>TACGGCGTGCAGTGCTTCAGCCGCTACCCCGACCACATGAAGCAGCACGACTTCTTCAAGTCCGCC<br>ATGCCCGAAGGCTACGTCCAGGAGCGCACCATCTTCTTCAAGGACGACGGCAACTACAAGACCCGC<br>GCCGAGGTGAAGTTCGAGGGCGACACCCTGGTGAACCGCATCGAGCTGAAGGGCATCGACTTCAAG<br>GAGGACGGCAACATCCTGGGGCACAAGCTGGAGTACAACTACAACAGCCACAACGTCTATATCATG<br>GCCGACAAGCAGAAGAACGGCATCAAGGTGAACTTCAAGATCCGCCACAACATCGAGGACGGCAGC<br>GTGCAGCTCGCCGACCACTACCAGCAGAACACCCCCATCGGCGACGGCCCCGTGCTGCTGCCCGAC<br>AACCACTACCTGAGCACCCAGTCCGCCCTGAGCAAAGACCCCAACGAGAAGCGCGATCACATGGTC<br>CTGCTGGAGTTCGTGACCGCCGCCGGGATCACTCTCGGCATGGACGAGCTGTACAAGTAA |
| eGFP-KRAS | ATGgctagcGTATCCAAAGGCGAGGAGTTGTTTACTGGGGTCGTGCCAATACTTGTCGAACTGGAT<br>GGCGACGTTAATGGTCACAAGTTTAGTGTTTCTGGGGAAGGTGAGGGGGATGCAACGTATGGGAAA<br>CTTACGTTGAAATTTATTTGTACGACCGGGAAACTCCCAGTCCCTTGGCCCACTCTTGTTACGACA<br>CTGACGTACGGCGTTCAGTGCTTTAGTAGATACCCAGACCATATGAAGCAACATGATTTCTTCAAA<br>AGTGCTATGCCGGAGGGCTATGTGCAAGAACGCACTATATTTTTCAAGGATGATGGGAACTATAAA<br>ACACGAGCGGAAGTTAAGTTTGAGGGCGATACGCTGGTGAATCGAATAGAACTGAAGGGTATTGAC<br>TTCAAAGAAGACGGGAACATATTGGGACATAAGCTCGAGTACAACTACAACTCTCACAATGTTTAT<br>ATTATGGCTGACAAGCAGAAGAATGGAATAAAGGTGAATTTTAAGATCAGGCACAACATTGAAGAT<br>GGTAGTGTACAATTGGCTGATCACTACCAACAGAACACACCGATCGGAGACGGACCAGTTTTGCTC<br>CCTGACAATCACTACCTGTCCACCCAGTCCGCCCTTTCAAAAGATCCGAATGAAAAGCGAGACCAC<br>ATGGTCCTCCTCGAGTTCGTGACGGCGGCGGGAATTACTTTGGGCATGGACGAACTCTACAAAGGA<br>TCCGGTAGTGGCtccggaACAGAATACAAACTGGTAGTCGTCGGAGCCGGAGGGGTAGGAAAATCC<br>GCCCTCACAATCCAGCTTATCCAGAACCATTTCGTTGACGAATACGATCCGACAATTGAAGACAGC<br>TATCGAAAACAGGTAGTGATAGACGGCGAGACCTGTCTTCTTGACATTCTTGATACAGCCGGTCAG<br>GAAGAATATTCAGCGATGCGGGACCAATACATGAGAACGGGAGAGGGGTTTCTCTGCGTATTTGCG<br>ATTAATAATACAAAGTCTTTTGAAGACATACACCACTACAGAGAGCAGATCAAACGTGTTAAGGAT<br>TCCGAAGATGTACCGATGGTTCTGGTTGGTAACAAATGCGACTTGCCATCAAGAACGGTGGACACA<br>AAACAAGCTCAGGACTTGGCCCGGAGCTACGGGATTCCTTTCATTGAGACTTCTGCCAAAACCAGG<br>CAGGGAGTAGACGACGCATTCTATACGCTCGTTCGAGAGATCCGCAAACATAAAGAGAAGATGAGT<br>AAGGACGGTAAGAAGAAGAAGAAGAAATCCAAGACAAAATGCGTCATAATGTGA |

| | |
|---|---|
| NanoLuc-HaloTag | ATGGTCTTCACACTCGAAGATTTCGTTGGGGACTGGCGACAGACAGCCGGCTACAACCTGGACCAA<br>GTCCTTGAACAGGGAGGTGTGTCCAGTTTGTTTCAGAATCTCGGGGTGTCCGTAACTCCGATCCAA<br>AGGATTGTCCTGAGCGGTGAAAATGGGCTGAAGATCGACATCCATGTCATCATCCCGTATGAAGGT<br>CTGAGCGGCGACCAAATGGGCCAGATCGAAAAAATTTTTAAGGTGGTGTACCCTGTGGATGATCAT<br>CACTTTAAGGTGATCCTGCACTATGGCACACTGGTAATCGACGGGGTTACGCCGAACATGATCGAC<br>TATTTCGGACGGCCGTATGAAGGCATCGCCGTGTTCGACGGCAAAAAGATCACTGTAACAGGGACC<br>CTGTGGAACGGCAACAAAATTATCGACGAGCGCCTGATCAACCCCGACGGCTCCCTGCTGTTCCGA<br>GTAACCATCAACGGAGTGACCGGCTGGCGGCTGTGCGAACGCATTCTGGCGggctcgagcggcGGA<br>TCCGAAATCGGTACTGGCTTTCCATTCGACCCCCATTATGTGGAAGTCCTGGGCGAGCGCATGCAC<br>TACGTCGATGTTGGTCCGCGCGATGGCACCCCTGTGCTGTTCCTGCACGGTAACCCGACCTCCTCC<br>TACGTGTGGCGCAACATCATCCCGCATGTTGCACCGACCCATCGCTGCATTGCTCCAGACCTGATC<br>GGTATGGGCAAATCCGACAAACCAGACCTGGGTTATTTCTTCGACGACCACGTCCGCTTCATGGAT<br>GCCTTCATCGAAGCCCTGGGTCTGGAAGAGGTCGTCCTGGTCATTCACGACTGGGGCTCCGCTCTG<br>GGTTTCCACTGGGCCAAGCGCAATCCAGAGCGCGTCAAAGGTATTGCATTTATGGAGTTCATCCGC<br>CCTATCCCGACCTGGGACGAATGGCCAGAATTTGCCCGCGAGACCTTCCAGGCCTTCCGCACCACC<br>GACGTCGGCCGCAAGCTGATCATCGATCAGAACGTTTTTATCGAGGGTACGCTGCCGATGGGTGTC<br>GTCCGCCCGCTGACTGAAGTCGAGATGGACCATTACCGCGAGCCGTTCCTGAATCCTGTTGACCGC<br>GAGCCACTGTGGCGCTTCCCAAACGAGCTGCCAATCGCCGGTGAGCCAGCGAACATCGTCGCGCTG<br>GTCGAAGAATACATGGACTGGCTGCACCAGTCCCCTGTCCCGAAGCTGCTGTTCTGGGGCACCCCA<br>GGCGTTCTGATCCCACCGGCCGAAGCCGCTCGCCTGGCCAAAAGCCTGCCTAACTGCAAGGCTGTG<br>GACATCGGCCCGGGTCTGAATCTGCTGCAAGAAGACAACCCGGACCTGATCGGCAGCGAGATCGCG<br>CGCTGGCTGTCTACTCTGGAGATTTCCGGTTAATGA |
| NanoLuc-KRAS | ATGGTCTTCACACTCGAAGATTTCGTTGGGGACTGGCGACAGACAGCCGGCTACAACCTGGACCAA<br>GTCCTTGAACAGGGAGGTGTGTCCAGTTTGTTTCAGAATCTCGGGGTGTCCGTAACTCCGATCCAA<br>AGGATTGTCCTGAGCGGTGAAAATGGGCTGAAGATCGACATCCATGTCATCATCCCGTATGAAGGT<br>CTGAGCGGCGACCAAATGGGCCAGATCGAAAAAATTTTTAAGGTGGTGTACCCTGTGGATGATCAT<br>CACTTTAAGGTGATCCTGCACTATGGCACACTGGTAATCGACGGGGTTACGCCGAACATGATCGAC<br>TATTTCGGACGGCCGTATGAAGGCATCGCCGTGTTCGACGGCAAAAAGATCACTGTAACAGGGACC<br>CTGTGGAACGGCAACAAAATTATCGACGAGCGCCTGATCAACCCCGACGGCTCCCTGCTGTTCCGA<br>GTAACCATCAACGGAGTGACCGGCTGGCGGCTGTGCGAACGCATTCTGGCGggctcgagcggcGGA<br>TCCGGTAGTGGCtccggaACCGAGTACAAGCTGGTTGTTGTAGGCGCAGGTGGCGTGGGGAAGAGT<br>GCTCTTACTATTCAGCTCATACAAAACCATTTCGTTGATGAATACGACCCCACTATAGAAGATAGC<br>TACCGGAAGCAAGTTGTAATCGACGGTGAAACCTGTCTGTTGGATATACTTGATACCGCAGGTCAG<br>GAGGAATACTCTGCCATGCGAGACCAATATATGAGGACTGGCGAGGGATTTCTTTGCGTATTCGCG<br>ATTAACAACACGAAGTCCTTTGAGGATATACACCACTACAGGGAACAGATAAAGCGGGTCAAAGAC<br>AGCGAAGACGTTCCGATGGTACTGGTGGGTAATAAGTGCGACCTGCCTTCACGCACAGTTGACACA<br>AAGCAGGCGCAAGATTTGGCTCGATCTTATGGCATCCCGTTCATAGAAACATCCGCTAAGACGAGG<br>CAGGGTGTAGATGACGCTTTTTATACGCTCGTCCGCGAAATACGCAAGCACAAGGAAAAGATGAGC<br>AAGGACGGCAAAAAAAAGAAGAAGAAGTCAAAAACTAAATGCGTTATCATGTGA |
| NanoLuc-HRAS | ATGGTCTTCACACTCGAAGATTTCGTTGGGGACTGGCGACAGACAGCCGGCTACAACCTGGACCAA<br>GTCCTTGAACAGGGAGGTGTGTCCAGTTTGTTTCAGAATCTCGGGGTGTCCGTAACTCCGATCCAA<br>AGGATTGTCCTGAGCGGTGAAAATGGGCTGAAGATCGACATCCATGTCATCATCCCGTATGAAGGT<br>CTGAGCGGCGACCAAATGGGCCAGATCGAAAAAATTTTTAAGGTGGTGTACCCTGTGGATGATCAT<br>CACTTTAAGGTGATCCTGCACTATGGCACACTGGTAATCGACGGGGTTACGCCGAACATGATCGAC<br>TATTTCGGACGGCCGTATGAAGGCATCGCCGTGTTCGACGGCAAAAAGATCACTGTAACAGGGACC<br>CTGTGGAACGGCAACAAAATTATCGACGAGCGCCTGATCAACCCCGACGGCTCCCTGCTGTTCCGA<br>GTAACCATCAACGGAGTGACCGGCTGGCGGCTGTGCGAACGCATTCTGGCGggctcgagcggcGGA<br>TCCGGTAGTGGCtccggaACCGAATACAAGCTGGTGGTCGTGGGTGCAGGCGGGGTTGGTAAGAGT<br>GCTCTCACGATTCAGCTTATTCAAAACCACTTTGTAGATGAGTATGATCCCACAATAGAGGATTCA<br>TATCGCAAACAAGTTGTGATTGATGGGGAAACCTGCCTTCTTGACATTCTTGACACCGCTGGCCAA<br>GAAGAGTATTCCGCAATGCGGGACCAGTATATGCGGACTGGCGAGGGATTCCTGTGCGTTTTCGCA |

| | |
|---|---|
| | ATAAACAATACCAAATCTTTTGAGGACATCCATCAATACAGAGAGCAGATTAAGAGAGTCAAAGAT<br>TCAGACGACGTGCCAATGGTCCTTGTCGGGAATAAATGTGACCTTGCAGCTAGAACGGTTGAGTCC<br>CGACAAGCCCAAGACCTTGCACGATCTTACGGTATCCCATACATAGAAACGTCCGCCAAGACGAGA<br>CAGGGCGTCGAGGACGCCTTTTACACACTCGTCAGGGAGATTCGACAACACAAGCTCAGGAAGCTC<br>AACCCACCAGATGAATCAGGCCCTGGATGTATGAGTTGCAAGTGTGTGTTGTCTTGA |
| NanoLuc-NRAS | ATGGTCTTCACACTCGAAGATTTCGTTGGGGACTGGCGACAGACAGCCGGCTACAACCTGGACCAA<br>GTCCTTGAACAGGGAGGTGTGTCCAGTTTGTTTCAGAATCTCGGGGTGTCCGTAACTCCGATCCAA<br>AGGATTGTCCTGAGCGGTGAAAATGGGCTGAAGATCGACATCCATGTCATCATCCCGTATGAAGGT<br>CTGAGCGGCGACCAAATGGGCCAGATCGAAAAAATTTTTAAGGTGGTGTACCCTGTGGATGATCAT<br>CACTTTAAGGTGATCCTGCACTATGGCACACTGGTAATCGACGGGGTTACGCCGAACATGATCGAC<br>TATTTCGGACGGCCGTATGAAGGCATCGCCGTGTTCGACGGCAAAAAGATCACTGTAACAGGGACC<br>CTGTGGAACGGCAACAAAATTATCGACGAGCGCCTGATCAACCCCGACGGCTCCCTGCTGTTCCGA<br>GTAACCATCAACGGAGTGACCGGCTGGCGGCTGTGCGAACGCATTCTGGCGggctcgagcggcGGA<br>TCCGGTAGTGGCtccggaACAGAGTACAAACTTGTAGTGGTCGGAGCCGGAGGCGTGGGAAAAGC<br>GCACTTACTATACAGCTTATCCAGAATCACTTTGTCGATGAGTACGACCCCACGATTGAAGATTCC<br>TATAGAAAGCAGGTTGTAATAGATGGGGAAACATGCCTCCTTGACATACTCGACACCGCCGGACAG<br>GAGGAATACAGTGCCATGCGAGACCAGTATATGCGGACCGGAGAAGGTTTCCTGTGTGTTTTTGCC<br>ATAAATAACTCCAAATCCTTTGCAGATATTAATCTCTACCGGGAACAAATAAAAAGAGTCAAGGAT<br>TCAGATGATGTACCAATGGTGCTGGTCGGTAATAAATGTGATCTTCCGACCCGGACTGTTGATACG<br>AAACAAGCCCACGAACTTGCTAAGTCTTATGGTATCCCCTTCATTGAGACCAGCGCAAAAACCCGA<br>CAAGGCGTAGAGGATGCCTTCTATACTTTGGTACGCGAGATCCGCCAGTATAGGATGAAGAAGCTG<br>AACTCATCAGATGACGGCACACAGGGTTGCATGGGGTTGCCGTGCGTTGTAATGTGA |
| NanoLuc-KRAS$^{G12D}$ | ATGGTCTTCACACTCGAAGATTTCGTTGGGGACTGGCGACAGACAGCCGGCTACAACCTGGACCAA<br>GTCCTTGAACAGGGAGGTGTGTCCAGTTTGTTTCAGAATCTCGGGGTGTCCGTAACTCCGATCCAA<br>AGGATTGTCCTGAGCGGTGAAAATGGGCTGAAGATCGACATCCATGTCATCATCCCGTATGAAGGT<br>CTGAGCGGCGACCAAATGGGCCAGATCGAAAAAATTTTTAAGGTGGTGTACCCTGTGGATGATCAT<br>CACTTTAAGGTGATCCTGCACTATGGCACACTGGTAATCGACGGGGTTACGCCGAACATGATCGAC<br>TATTTCGGACGGCCGTATGAAGGCATCGCCGTGTTCGACGGCAAAAAGATCACTGTAACAGGGACC<br>CTGTGGAACGGCAACAAAATTATCGACGAGCGCCTGATCAACCCCGACGGCTCCCTGCTGTTCCGA<br>GTAACCATCAACGGAGTGACCGGCTGGCGGCTGTGCGAACGCATTCTGGCGggctcgagcggcGGA<br>TCCGGTAGTGGCtccggaACCGAGTACAAGCTGGTTGTTGTAGGCGCAgacGGCGTGGGAAGAGT<br>GCTCTTACTATTCAGCTCATACAAAACCATTTCGTTGATGAATACGACCCCACTATAGAAGATAGC<br>TACCGGAAGCAAGTTGTAATCGACGGTGAAACCTGTCTGTTGGATATACTTGATACCGCAGGTCAG<br>GAGGAATACTCTGCCATGCGAGACCAATATATGAGGACTGGCGAGGGATTTCTTTGCGTATTCGCG<br>ATTAACAACACGAAGTCCTTTGAGGATATACACCACTACAGGGAACAGATAAAGCGGGTCAAAGAC<br>AGCGAAGACGTTCCGATGGTACTGGTGGGTAATAAGTGCGACCTGCCTTCACGCACAGTTGACACA<br>AAGCAGGCGCAAGATTTGGCTCGATCTTATGGCATCCCGTTCATAGAAACATCCGCTAAGACGAGG<br>CAGGGTGTAGATGACGCTTTTTATACGCTCGTCCGCGAAATACGCAAGCACAAGGAAAAGATGAGC<br>AAGGACGGCAAAAAAAAGAAGAAGAAGTCAAAAACTAAATGCGTTATCATGTGA |
| NanoLuc-KRAS$^{R135K}$ | ATGGTCTTCACACTCGAAGATTTCGTTGGGGACTGGCGACAGACAGCCGGCTACAACCTGGACCAA<br>GTCCTTGAACAGGGAGGTGTGTCCAGTTTGTTTCAGAATCTCGGGGTGTCCGTAACTCCGATCCAA<br>AGGATTGTCCTGAGCGGTGAAAATGGGCTGAAGATCGACATCCATGTCATCATCCCGTATGAAGGT<br>CTGAGCGGCGACCAAATGGGCCAGATCGAAAAAATTTTTAAGGTGGTGTACCCTGTGGATGATCAT<br>CACTTTAAGGTGATCCTGCACTATGGCACACTGGTAATCGACGGGGTTACGCCGAACATGATCGAC<br>TATTTCGGACGGCCGTATGAAGGCATCGCCGTGTTCGACGGCAAAAAGATCACTGTAACAGGGACC<br>CTGTGGAACGGCAACAAAATTATCGACGAGCGCCTGATCAACCCCGACGGCTCCCTGCTGTTCCGA<br>GTAACCATCAACGGAGTGACCGGCTGGCGGCTGTGCGAACGCATTCTGGCGggctcgagcggcGGA<br>TCCGGTAGTGGCtccggaACCGAGTACAAGCTGGTTGTTGTAGGCGCAGGTGGCGTGGGAAGAGT<br>GCTCTTACTATTCAGCTCATACAAAACCATTTCGTTGATGAATACGACCCCACTATAGAAGATAGC<br>TACCGGAAGCAAGTTGTAATCGACGGTGAAACCTGTCTGTTGGATATACTTGATACCGCAGGTCAG<br>GAGGAATACTCTGCCATGCGAGACCAATATATGAGGACTGGCGAGGGATTTCTTTGCGTATTCGCG |

| | |
|---|---|
| | ATTAACAACACGAAGTCCTTTGAGGATATACACCACTACAGGGAACAGATAAAGCGGGTCAAAGAC<br>AGCGAAGACGTTCCGATGGTACTGGTGGGTAATAAGTGCGACCTGCCTTCACGCACAGTTGACACA<br>AAGCAGGCGCAAGATTTGGCTaagTCTTATGGCATCCCGTTCATAGAAACATCCGCTAAGACGAGG<br>CAGGGTGTAGATGACGCTTTTTATACGCTCGTCCGCGAAATACGCAAGCACAAGGAAAAGATGAGC<br>AAGGACGGCAAAAAAAAGAAGAAGAAGTCAAAAACTAAATGCGTTATCATGTGA |
| NanoLuc-KRAS$^{G12C}$ | ATGGTCTTCACACTCGAAGATTTCGTTGGGGACTGGCGACAGACAGCCGGCTACAACCTGGACCAA<br>GTCCTTGAACAGGGAGGTGTGTCCAGTTTGTTTCAGAATCTCGGGGTGTCCGTAACTCCGATCCAA<br>AGGATTGTCCTGAGCGGTGAAAATGGGCTGAAGATCGACATCCATGTCATCATCCCGTATGAAGGT<br>CTGAGCGGCGACCAAATGGGCCAGATCGAAAAAATTTTTAAGGTGGTGTACCCTGTGGATGATCAT<br>CACTTTAAGGTGATCCTGCACTATGGCACACTGGTAATCGACGGGGTTACGCCGAACATGATCGAC<br>TATTTCGGACGGCCGTATGAAGGCATCGCCGTGTTCGACGGCAAAAAGATCACTGTAACAGGGACC<br>CTGTGGAACGGCAACAAAATTATCGACGAGCGCCTGATCAACCCCGACGGCTCCCTGCTGTTCCGA<br>GTAACCATCAACGGAGTGACCGGCTGGCGGCTGTGCGAACGCATTCTGGCGggctcgagcggcGGA<br>TCCGGTAGTGGCtccggaACCGAGTACAAGCTGGTTGTTGTAGGCGCAtGTGGCGTGGGAAGAGT<br>GCTCTTACTATTCAGCTCATACAAAACCATTTCGTTGATGAATACGACCCCACTATAGAAGATAGC<br>TACCGGAAGCAAGTTGTAATCGACGGTGAAACCTGTCTGTTGGATATACTTGATACCGCAGGTCAG<br>GAGGAATACTCTGCCATGCGAGACCAATATATGAGGACTGGCGAGGGATTTCTTTGCGTATTCGCG<br>ATTAACAACACGAAGTCCTTTGAGGATATACACCACTACAGGGAACAGATAAAGCGGGTCAAAGAC<br>AGCGAAGACGTTCCGATGGTACTGGTGGGTAATAAGTGCGACCTGCCTTCACGCACAGTTGACACA<br>AAGCAGGCGCAAGATTTGGCTCGATCTTATGGCATCCCGTTCATAGAAACATCCGCTAAGACGAGG<br>CAGGGTGTAGATGACGCTTTTTATACGCTCGTCCGCGAAATACGCAAGCACAAGGAAAAGATGAGC<br>AAGGACGGCAAAAAAAAGAAGAAGAAGTCAAAAACTAAATGCGTTATCATGTGA |
| NanoLuc-KRAS$^{G12V}$ | ATGGTCTTCACACTCGAAGATTTCGTTGGGGACTGGCGACAGACAGCCGGCTACAACCTGGACCAA<br>GTCCTTGAACAGGGAGGTGTGTCCAGTTTGTTTCAGAATCTCGGGGTGTCCGTAACTCCGATCCAA<br>AGGATTGTCCTGAGCGGTGAAAATGGGCTGAAGATCGACATCCATGTCATCATCCCGTATGAAGGT<br>CTGAGCGGCGACCAAATGGGCCAGATCGAAAAAATTTTTAAGGTGGTGTACCCTGTGGATGATCAT<br>CACTTTAAGGTGATCCTGCACTATGGCACACTGGTAATCGACGGGGTTACGCCGAACATGATCGAC<br>TATTTCGGACGGCCGTATGAAGGCATCGCCGTGTTCGACGGCAAAAAGATCACTGTAACAGGGACC<br>CTGTGGAACGGCAACAAAATTATCGACGAGCGCCTGATCAACCCCGACGGCTCCCTGCTGTTCCGA<br>GTAACCATCAACGGAGTGACCGGCTGGCGGCTGTGCGAACGCATTCTGGCGggctcgagcggcGGA<br>TCCGGTAGTGGCtccggaACCGAGTACAAGCTGGTTGTTGTAGGCGCAGtTGGCGTGGGAAGAGT<br>GCTCTTACTATTCAGCTCATACAAAACCATTTCGTTGATGAATACGACCCCACTATAGAAGATAGC<br>TACCGGAAGCAAGTTGTAATCGACGGTGAAACCTGTCTGTTGGATATACTTGATACCGCAGGTCAG<br>GAGGAATACTCTGCCATGCGAGACCAATATATGAGGACTGGCGAGGGATTTCTTTGCGTATTCGCG<br>ATTAACAACACGAAGTCCTTTGAGGATATACACCACTACAGGGAACAGATAAAGCGGGTCAAAGAC<br>AGCGAAGACGTTCCGATGGTACTGGTGGGTAATAAGTGCGACCTGCCTTCACGCACAGTTGACACA<br>AAGCAGGCGCAAGATTTGGCTCGATCTTATGGCATCCCGTTCATAGAAACATCCGCTAAGACGAGG<br>CAGGGTGTAGATGACGCTTTTTATACGCTCGTCCGCGAAATACGCAAGCACAAGGAAAAGATGAGC<br>AAGGACGGCAAAAAAAAGAAGAAGAAGTCAAAAACTAAATGCGTTATCATGTGA |
| NanoLuc-KRAS$^{Q61H}$ | ATGGTCTTCACACTCGAAGATTTCGTTGGGGACTGGCGACAGACAGCCGGCTACAACCTGGACCAA<br>GTCCTTGAACAGGGAGGTGTGTCCAGTTTGTTTCAGAATCTCGGGGTGTCCGTAACTCCGATCCAA<br>AGGATTGTCCTGAGCGGTGAAAATGGGCTGAAGATCGACATCCATGTCATCATCCCGTATGAAGGT<br>CTGAGCGGCGACCAAATGGGCCAGATCGAAAAAATTTTTAAGGTGGTGTACCCTGTGGATGATCAT<br>CACTTTAAGGTGATCCTGCACTATGGCACACTGGTAATCGACGGGGTTACGCCGAACATGATCGAC<br>TATTTCGGACGGCCGTATGAAGGCATCGCCGTGTTCGACGGCAAAAAGATCACTGTAACAGGGACC<br>CTGTGGAACGGCAACAAAATTATCGACGAGCGCCTGATCAACCCCGACGGCTCCCTGCTGTTCCGA<br>GTAACCATCAACGGAGTGACCGGCTGGCGGCTGTGCGAACGCATTCTGGCGggctcgagcggcGGA<br>TCCGGTAGTGGCtccggaACCGAGTACAAGCTGGTTGTTGTAGGCGCAGGTGGCGTGGGAAGAGT<br>GCTCTTACTATTCAGCTCATACAAAACCATTTCGTTGATGAATACGACCCCACTATAGAAGATAGC<br>TACCGGAAGCAAGTTGTAATCGACGGTGAAACCTGTCTGTTGGATATACTTGATACCGCAGGTCAt<br>GAGGAATACTCTGCCATGCGAGACCAATATATGAGGACTGGCGAGGGATTTCTTTGCGTATTCGCG |

| | |
|---|---|
| | ATTAACAACACGAAGTCCTTTGAGGATATACACCACTACAGGGAACAGATAAAGCGGGTCAAAGAC<br>AGCGAAGACGTTCCGATGGTACTGGTGGGTAATAAGTGCGACCTGCCTTCACGCACAGTTGACACA<br>AAGCAGGCGCAAGATTTGGCTCGATCTTATGGCATCCCGTTCATAGAAACATCCGCTAAGACGAGG<br>CAGGGTGTAGATGACGCTTTTTATACGCTCGTCCGCGAAATACGCAAGCACAAGGAAAAGATGAGC<br>AAGGACGGCAAAAAAAAGAAGAAGAAGTCAAAAACTAAATGCGTTATCATGTGA |
| FLAG-K19-SPOP$_{167-374}$ | ATGGATTACAAGGACGACGACGACAAGgctagcGACCTTGGAAAGAAACTTTTGGAAGCTGCCCGC<br>GCGGGGCAAGACGATGAAGTGCGAATATTGATGGCGAATGGCGCTGACGTCAACGCCAGCGACCGC<br>TGGGGTTGGACACCTCTGCACTTGGCCGCATGGTGGGGCCATCTTGAAATAGTAGAAGTTCTTCTG<br>AAGAGAGGGGCCGATGTTTCCGCTGCCGATTTGCACGGCCAATCCCCTCTCCACCTTGCGGCTATG<br>GTGGGACATTTGGAGATAGTCGAGGTGTTGCTTAAGTATGGGGCAGACGTCAACGCTAAAGACACT<br>ATGGGTGCAACGCCCTTGCACCTTGCAGCGCGCAGTGGTCACCTGGAGATCGTTGAAGAGTTGCTG<br>AAGAATGGAGCAGATATGAATGCTCAAGATAAGTTCGGTAAGACCACATTTGATATTTCCACGGAT<br>AATGGGAATGAGGACCTCGCCGAAATACTCCAGAAGTTGGGCAGTGGTAGTGGCtccggaAGCGTG<br>AACATCTCCGGCCAGAACACAATGAACATGGTCAAGGTGCCCGAGTGCAGACTGGCCGACGAGCTG<br>GGAGGACTGTGGGAGAACTCCAGGTTTACCGACTGCTGCCTGTGCGTGGCCGGCCAAGAGTTCCAA<br>GCCCACAAAGCCATCCTGGCCGCTAGGTCCCCCGTGTTCAGCGCCATGTTCGAGCACGAGATGGAG<br>GAGTCCAAGAAGAACAGAGTGGAGATTAACGATGTGGAGCCCGAGGTGTTCAAAGAAATGATGTGC<br>TTCATCTACACCGGCAAGGCCCCCAACCTGGATAAAATGGCCGATGACCTGCTGGCCGCCGCCGAT<br>AAGTACGCCCTGGAGAGACTGAAGGTGATGTGCGAGGACGCTCTGTGTTCCAACCTGTCCGTGGAA<br>AATGCCGCCGAGATCCTCATCCTGGCCGACCTGCATAGCGCCGACCAGCTGAAAACCCAGGCCGTG<br>GACTTCATCAACTATCACGCTTCCGACGTGCTGGAGACCAGCGGATGGAAGAGCATGGTGGTGAGC<br>CATCCCCATCTCGTGGCCGAAGCCTACAGGAGCCTGGCaAGCGCCCAGTGTCCCTTTCTGGGCCCT<br>CCCAGGAAGAGACTGAAACAGAGCTGA |
| NanoLuc-KRAS$^{H95Q}$ | ATGGTCTTCACACTCGAAGATTTCGTTGGGGACTGGCGACAGACAGCCGGCTACAACCTGGACCAA<br>GTCCTTGAACAGGGAGGTGTGTCCAGTTTGTTTCAGAATCTCGGGGTGTCCGTAACTCCGATCCAA<br>AGGATTGTCCTGAGCGGTGAAAATGGGCTGAAGATCGACATCCATGTCATCATCCCGTATGAAGGT<br>CTGAGCGGCGACCAAATGGGCCAGATCGAAAAAATTTTTAAGGTGGTGTACCCTGTGGATGATCAT<br>CACTTTAAGGTGATCCTGCACTATGGCACACTGGTAATCGACGGGGTTACGCCGAACATGATCGAC<br>TATTTCGGACGGCCGTATGAAGGCATCGCCGTGTTCGACGGCAAAAAGATCACTGTAACAGGGACC<br>CTGTGGAACGGCAACAAAATTATCGACGAGCGCCTGATCAACCCCGACGGCTCCCTGCTGTTCCGA<br>GTAACCATCAACGGAGTGACCGGCTGGCGGCTGTGCGAACGCATTCTGGCGggctcgagcggcGGA<br>TCCGGTAGTGGCtccggaACCGAGTACAAGCTGGTTGTTGTAGGCGCAGGTGGCGTGGGAAGAGT<br>GCTCTTACTATTCAGCTCATACAAAACCATTTCGTTGATGAATACGACCCCACTATAGAAGATAGC<br>TACCGGAAGCAAGTTGTAATCGACGGTGAAACCTGTCTGTTGGATATACTTGATACCGCAGGTCAG<br>GAGGAATACTCTGCCATGCGAGACCAATATATGAGGACTGGCGAGGGATTTCTTTGCGTATTCGCG<br>ATTAACAACACGAAGTCCTTTGAGGATATACACCAgTACAGGGAACAGATAAAGCGGGTCAAAGAC<br>AGCGAAGACGTTCCGATGGTACTGGTGGGTAATAAGTGCGACCTGCCTTCACGCACAGTTGACACA<br>AAGCAGGCGCAAGATTTGGCTCGATCTTATGGCATCCCGTTCATAGAAACATCCGCTAAGACGAGG<br>CAGGGTGTAGATGACGCTTTTTATACGCTCGTCCGCGAAATACGCAAGCACAAGGAAAAGATGAGC<br>AAGGACGGCAAAAAAAAGAAGAAGAAGTCAAAAACTAAATGCGTTATCATGTGA |
| NanoLuc-KRAS$^{H95L}$ | ATGGTCTTCACACTCGAAGATTTCGTTGGGGACTGGCGACAGACAGCCGGCTACAACCTGGACCAA<br>GTCCTTGAACAGGGAGGTGTGTCCAGTTTGTTTCAGAATCTCGGGGTGTCCGTAACTCCGATCCAA<br>AGGATTGTCCTGAGCGGTGAAAATGGGCTGAAGATCGACATCCATGTCATCATCCCGTATGAAGGT<br>CTGAGCGGCGACCAAATGGGCCAGATCGAAAAAATTTTTAAGGTGGTGTACCCTGTGGATGATCAT<br>CACTTTAAGGTGATCCTGCACTATGGCACACTGGTAATCGACGGGGTTACGCCGAACATGATCGAC<br>TATTTCGGACGGCCGTATGAAGGCATCGCCGTGTTCGACGGCAAAAAGATCACTGTAACAGGGACC<br>CTGTGGAACGGCAACAAAATTATCGACGAGCGCCTGATCAACCCCGACGGCTCCCTGCTGTTCCGA<br>GTAACCATCAACGGAGTGACCGGCTGGCGGCTGTGCGAACGCATTCTGGCGggctcgagcggcGGA<br>TCCGGTAGTGGCtccggaACCGAGTACAAGCTGGTTGTTGTAGGCGCAGGTGGCGTGGGAAGAGT<br>GCTCTTACTATTCAGCTCATACAAAACCATTTCGTTGATGAATACGACCCCACTATAGAAGATAGC<br>TACCGGAAGCAAGTTGTAATCGACGGTGAAACCTGTCTGTTGGATATACTTGATACCGCAGGTCAG |

| | |
|---|---|
| | GAGGAATACTCTGCCATGCGAGACCAATATATGAGGACTGGCGAGGGATTTCTTTGCGTATTCGCG<br>ATTAACAACACGAAGTCCTTTGAGGATATACACCtCTACAGGGAACAGATAAAGCGGGTCAAAGAC<br>AGCGAAGACGTTCCGATGGTACTGGTGGGTAATAAGTGCGACCTGCCTTCACGCACAGTTGACACA<br>AAGCAGGCGCAAGATTTGGCTCGATCTTATGGCATCCCGTTCATAGAAACATCCGCTAAGACGAGG<br>CAGGGTGTAGATGACGCTTTTTATACGCTCGTCCGCGAAATACGCAAGCACAAGGAAAAGATGAGC<br>AAGGACGGCAAAAAAAAGAAGAAGAAGTCAAAAACTAAATGCGTTATCATGTGA |
